# FluoMALDI Imaging: An Integrated Multimodality Platform for Biologically Informed Spatial-Omics

**DOI:** 10.64898/2026.11.15.766883

**Authors:** Tae-Hun Hahm, Michael L. Piacentino, Waqar Ahmed, Maxime A. Siegler, Dalton R. Brown, Caitlin M. Tressler, Juan C. Troncoso, Arvind P. Pathak, Swati Tanwar, Ishan Barman, Vasiliki Mahairaki, Kristine Glunde

**Affiliations:** Applied Imaging Mass Spectrometry Core, Johns Hopkins University School of Medicine, Baltimore, MD, 21287 USA; Russell H. Morgan Department of Radiology and Radiological Science, Johns Hopkins University School of Medicine, Baltimore, MD, 21287 USA; Department of Cell Biology, Johns Hopkins University School of Medicine, Baltimore, MD, 21205 USA; Department of Genetic Medicine, Johns Hopkins University School of Medicine, Baltimore, MD, 21205 USA; Department of Chemistry, Johns Hopkins University, Baltimore, MD, 21218 USA; Department of Pathology, Division of Neuropathology, Johns Hopkins University School of Medicine, Baltimore, MD, 21205 USA; Department of Neurology, Johns Hopkins University School of Medicine, Baltimore, MD, 21231 USA; Department of Mechanical Engineering, Johns Hopkins University, Baltimore, MD, 21218, USA; Sidney Kimmel Comprehensive Cancer Center, Johns Hopkins University School of Medicine, Baltimore, MD, 21231 USA

## Abstract

FluoMALDI imaging integrates fluorescence microscopy and matrix-assisted laser desorption/ionization mass spectrometry imaging (MALDI-MSI) of tissues and cells, leveraging matrix-enhanced fluorescence signals to enable spatially guided functional molecular discovery. Here we report three broadly applicable FluoMALDI paradigms that link how genes, proteins, and organelles shape spatial molecular composition. These advances are achieved by integrating immuno-, HaloTag-, and dye-based labels into a unified FluoMALDI imaging framework. These approaches enabled us to interrogate cell-specific lipidomic compositions in human hippocampal vasculature, neural crest cells in chick embryo tissue, and human induced pluripotent stem cell-derived astrocytes. We also determined that fluorophore-MALDI matrix co-crystals resulted in FluoMALDI-induced fluorescence enhancements. Single-crystal X-ray crystallography showed that their co-crystallization produced localized, finite π-π interactions between two fluorophore molecules, or a fluorophore and a matrix molecule, throughout the crystal lattice. Together, these findings provide structural insight into the basis of fluorescence enhancement and support FluoMALDI as a broadly applicable multimodality imaging platform for biologically informed, cell-specific spatial-omics.

## Introduction

Biological function is governed by the spatial organization of molecules within tissues and cells. Fluorescence microscopy provides high-resolution maps of specific biomolecules, while matrix-assisted laser desorption/ionization mass spectrometry imaging (MALDI-MSI) offers label-free, multiplexed molecular profiling^1,2^. The ability to integrate these complementary modalities for the same specimen in a single workflow would enable direct interrogation of molecularly defined structures within their native spatial metabolic and lipidomic environments^3,4^. However, conventional multimodality imaging workflows typically require separate adjacent tissue sections and complex post hoc image alignment^5,6,7^, which frequently introduces spatial distortion and registration errors^1,2^. Existing multimodality platforms also remain limited in their compatibility with diverse molecular labeling strategies, their potential to enhance the fluorescence signal, and their applicability across different tissue types and MALDI ionization modes. A single-specimen approach that simultaneously enhances fluorescence signals and is compatible with a wide array of labels, while enabling mass spectrometric molecular profiling has remained elusive.

Hypothesis-driven biological investigations often require the selective labeling of cell types, tissue structures, gene expression states, or organelles to provide physiological and spatial context for molecular measurements. This is commonly achieved through immunofluorescence, an antibody-based approach linking molecular measurements to cellular identity^8^. While this approach is powerful for extracellular antigens, it is limited for intracellular antigens, because permeabilization with detergents can extract lipids and other metabolites from the sample. To circumvent this challenge, genetic labeling using HaloTag reporters provides a powerful alternative means to selectively identify and visualize unique cell populations, gene expression programs, and proteins of interest^9^. By linking fluorescence to genetically encoded biological features, these approaches establish direct connections between cellular identity and function, enabling hypothesis-driven interrogation of spatial molecular landscapes^9^. In particular, HaloTag labeling combines genetic specificity with the flexibility of synthetic fluorophores, offering bright and tunable signals that are well suited to FluoMALDI imaging^3,4^. Fluorescent organelle labeling uses small-molecule dyes that selectively accumulate within specific subcellular compartments, including nuclei or actin filaments, providing a versatile means of defining subcellular architecture^10^.

Here, we introduce FluoMALDI imaging, a multimodality workflow in which MALDI matrix co-crystallization simultaneously enhances the fluorescence signal while preserving mass spectrometry imaging capabilities within the same tissue section or cells, for interrogating specific cell types, tissue structures, gene expression states, and organelles within their biological context. We discovered that MALDI matrix application enhances, rather than degrades, fluorescence upon co-crystallization with the fluorophore^3,4^. Since both modalities interrogate the same physical substrate, FluoMALDI imaging requires simple rigid body image co-registration, eliminating geometric distortions associated with acquisition of images of sequential tissue sections^3,4^. We developed FluoMALDI imaging as a versatile and broadly applicable platform that integrates a wide range of fluorescence-labeling strategies. These include organic dyes such as SYTOX Green (dye-FluoMALDI); phalloidin or antibodies conjugated to Alexa Fluor 488, 568, or 647 (immuno-FluoMALDI); and genetically encoded HaloTag labeling with tetramethylrhodamine (TMR) (HaloTag-FluoMALDI). This platform is compatible with matrices optimized for both positive-ion (α-cyano-4-hydroxycinnamic acid, CHCA) and negative-ion (norharmane, nH) mode MALDI-MSI, enabling spatial molecular profiling across various biological systems. The main steps involved in the immuno-, HaloTag-, and dye-FluoMALDI imaging workflow are shown in **Fig. 1**.

**Figure 1:**
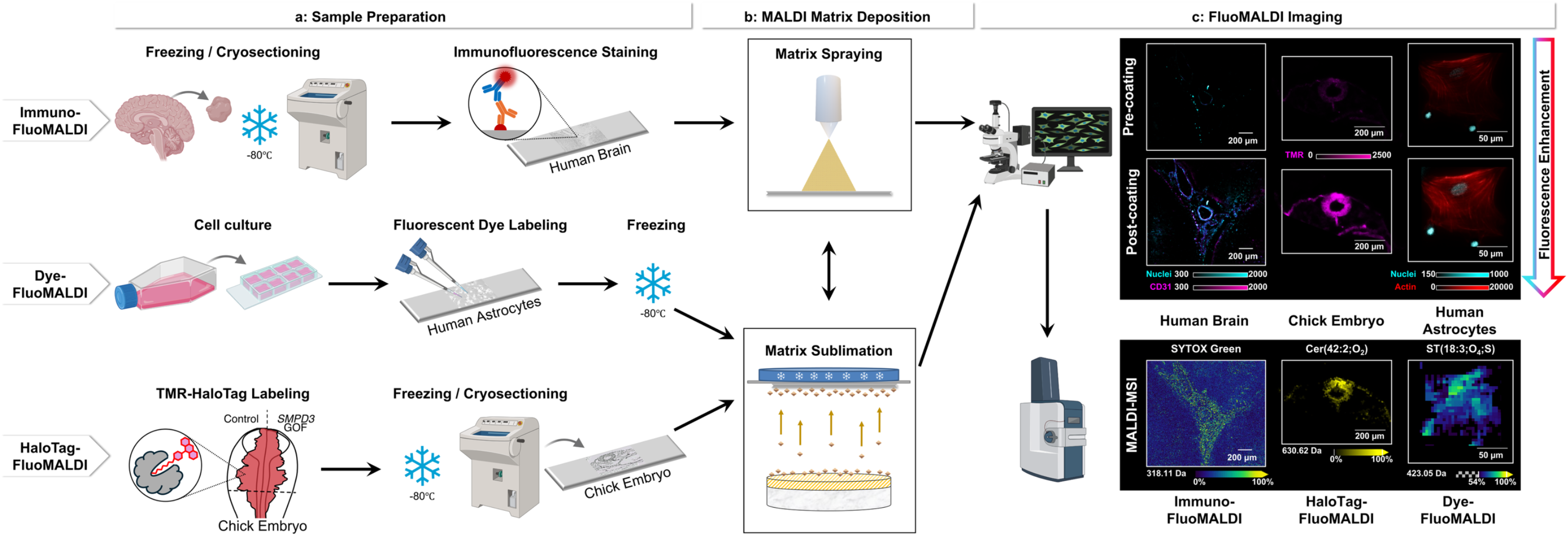
Overview of the main steps in immuno-, HaloTag- and dye-based FluoMALDI imaging workflows. Rows show the immuno- (top), dye- (middle) and HaloTag-FluoMALDI (bottom) workflows, and arrows indicate the order of steps. (**a**) Sample preparation. Human brain tissue is frozen, cryosectioned onto poly-L-lysine-coated indium tin oxide (ITO) glass slides and labeled by immunofluorescence with fluorophore-conjugated antibodies (immuno-FluoMALDI). Human astrocytes are cultured in flasks, seeded onto chamber slides, labeled with fluorescent dyes and then frozen (dye-FluoMALDI). Chick embryos are labeled with HaloTag and a TMR ligand and then frozen and cryosectioned onto poly-L-lysine-coated ITO glass slides (HaloTag-FluoMALDI), with the control side and the *SMPD3* gain-of-function (GOF) side of the embryo indicated. Labeling follows freezing in immuno-FluoMALDI but precedes freezing in dye- and HaloTag-FluoMALDI. (**b**) MALDI matrix deposition. The matrix (CHCA in the examples shown) is applied uniformly by automated spraying (immuno-FluoMALDI example) or sublimation (dye- and HaloTag-FluoMALDI examples). The double-headed arrow indicates that either method can be applied. (**c**) FluoMALDI imaging. The same specimens are imaged by slide-scanning fluorescence microscopy before (pre-coating) and after (post-coating) matrix deposition, which enhances the fluorescence signal (top, quantified in Figs. 2–4), and then by MALDI-MSI (bottom). Columns show human brain (immuno-FluoMALDI), chick embryo (HaloTag-FluoMALDI) and human astrocytes (dye-FluoMALDI). MALDI ion images show the nuclear stain SYTOX Green (*m/z* 318.11), Cer(42:2;O_2_) (*m/z* 630.62) and the putative sulfated steroid ST(18:3;O_4_;S) (*m/z* 423.05). Fluorescence channels are nuclei (SYTOX Green, cyan), CD31 (Alexa Fluor 647, magenta), TMR-HaloTag (magenta) and F-actin (phalloidin-Alexa Fluor 647, red). Color bars show fluorescence display ranges (arbitrary units) and relative MALDI-MSI intensity (%). Scale bars: 200 μm for human brain and chick embryo, 50 μm for human astrocytes. This workflow acquires fluorescence and MALDI-MSI data from the same specimen while increasing the fluorescence signal. Detailed workflows are shown in Figs. 2–4.

We demonstrated the utility of FluoMALDI imaging across three distinct biological paradigms, human hippocampal vasculature, chick embryo neural crest, and induced pluripotent stem cell (iPSC)-derived astrocytes. This showcases its broad applicability as a versatile single-specimen multimodality imaging platform that enables biologically informed, cell-specific spatial-omics.

## Results

### Immuno-FluoMALDI imaging of blood vessels in the human brain

Vascular dysfunction is implicated in Alzheimer’s disease and temporal lobe epilepsy, but the lipid composition of individual human hippocampal vessels remains poorly characterized^11^. The hippocampus is well suited for such analyses because its highly defined cytoarchitecture and vascular specialization enable spatially and cell-resolved interrogation of molecular phenotypes. The hippocampal trisynaptic circuit, which connects the entorhinal cortex, dentate gyrus (DG), cornu ammonis 3 (CA3), and CA1 through the perforant path, mossy fibers, and Schaffer collaterals, serves as a canonical neural pathway essential for learning, memory formation, and spatial navigation^11^. Distinct hippocampal subregions exhibit unique vascular architectures and metabolic requirements^11^. Notably, CA1 is supplied by terminal branches of the hippocampal artery with limited collateral circulation, making it particularly susceptible to ischemic injury^11^. We therefore asked whether immuno-FluoMALDI can assign lipid species to CD31-defined vascular compartments in human hippocampal tissue. To address this question, we developed immuno-FluoMALDI imaging to achieve spatially resolved, cell-specific lipid mapping in human hippocampal tissue.

The immuno-FluoMALDI imaging workflow begins with immunofluorescence staining for vascular markers, followed by fluorescence image acquisition, MALDI matrix application, fluorescence re-imaging to evaluate the FluoMALDI effect, and finally MALDI-MSI acquisition (**Fig. 2a** and **Supplementary Fig. S1a**). This single-section, sequential approach enables the direct mapping of lipid molecular signatures to immunofluorescence-defined vascular structures, providing cell-specific spatial context at subcellular resolution.

**Figure 2:**
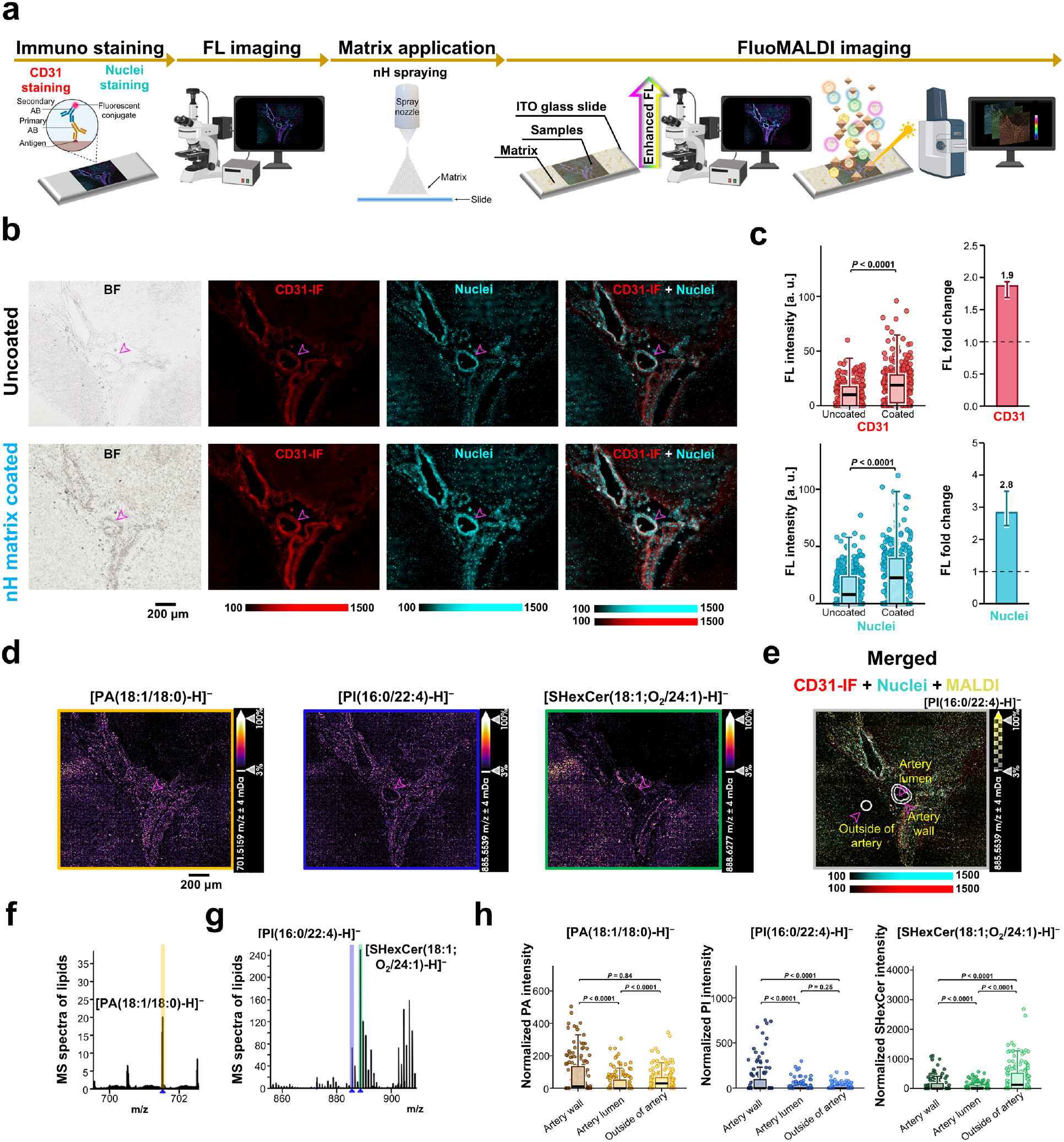
Immuno-FluoMALDI imaging of human brain tissue sections using nH matrix. (**a**) Workflow for immuno-FluoMALDI imaging and analysis. A human brain tissue section was mounted on a slide and subjected to immunofluorescence (IF) labeling. Optical microscopy was performed to compare the fluorescence of uncoated and coated tissue sections after nH matrix was applied by spray deposition. (b) Brightfield (BF) and fluorescence images of a human hippocampal section before (uncoated) and after coating with 0.9 µg/mm² nH. BF shows tissue morphology without a chromogen, and after coating also the matrix layer. CD31 was detected with an Alexa Fluor Plus 647-conjugated secondary antibody (CD31-IF, red) and nuclei with SYTOX Green (cyan), and both are visible only in the fluorescence channels. Magenta arrowheads mark the hippocampal artery analyzed in this study (a sulcal artery in the hippocampal fissure adjacent to CA1). Other CD31-positive vessels in the field of view were not analyzed. Scale bar: 200 μm. (**c**) Fluorescence intensity (a.u.) of CD31-IF (top) and nuclei (bottom) in the artery region before and after nH coating, with fold change (right, ratio of medians, 95% bootstrap confidence interval, the dashed line marks 1). Each point is one pixel (one section, one donor). Boxes show the median and interquartile range, and whiskers show 1.5 times the interquartile range. Statistical comparisons used two-sided Mann-Whitney U tests. (**d**) MALDI MS images in negative ion mode were obtained from the region of interest (ROI) containing hippocampal arteries (magenta arrows) in panel b. The MALDI images display PA(18:1/18:0) at *m/z***^−^**701.52, PI(16:0/22:4) at *m/z***^−^** 885.55, and SHexCer(18:1;O_2_/24:1) at *m/z***^−^** 888.63. Scale bars: 200 μm. (**e**) Co-registered post-coating fluorescence data (CD31-IF in red, nuclei in cyan) and MALDI imaging data of PI(16:0/22:4) (intensity scale converted to yellow color in overlayed image). Arrows indicate ROIs used for quantification. (**f** and **g**) Average MALDI MSI spectra from the MALDI-imaged ROI with color-coded labels for *m/z* signals shown as MALDI images in d. (**h**) TIC-normalized signal intensity of PA(18:1/18:0), PI(16:0/22:4) and SHexCer(18:1;O_2_/24:1) in artery wall, lumen and outside of artery (regions in e). Each point is one pixel (one artery, one section, one donor). Boxes show the median and interquartile range, and whiskers show 1.5 times the interquartile range. Kruskal–Wallis tests were followed by two-sided Dunn tests with Holm adjustment, and adjusted P values are shown. The hotspot intensity cutoff for pixel selection was defined by the hotspot quantile setting in the SCiLS Lab Settings dialog, configured at 99.0%. Exact P values and effect sizes are in Source Data.

Sections were labeled for endothelial CD31 (Alexa Fluor 647) and nuclei (SYTOX Green) and then coated with nH or CHCA matrix on adjacent serial sections (**Fig. 2b** and **Supplementary Fig. S1b**). MALDI matrix application substantially enhanced immunofluorescence intensity. CD31 immunofluorescence increased 1.9-fold with nH matrix (*P* < 0.0001; **Fig. 2c**) and 17.2-fold with CHCA matrix (*P* < 0.0001; **Supplementary Fig. S1c**), transforming faint vascular signals into well-defined vessel structures. Nuclear fluorescence showed comparable enhancement: 2.8-fold with nH (*P* < 0.0001) and 5.1-fold with CHCA (*P* < 0.0001) (**Fig. 2b, c** and **Supplementary Fig. S1b, c**). Vascular structures and endothelial cell boundaries remained well distinguishable after coating.

Although both matrices enhanced fluorescence, the optimal matrix–fluorophore pairing varied with experimental context (**Fig. 2** and **Supplementary Fig. S1**). In dried droplet experiments, nH demonstrated greater enhancement than CHCA for A647 (**Supplementary Fig. S2**), whereas on tissue sections, CHCA provided superior enhancement for both A647 and SYTOX Green (**Fig. 2** and **Supplementary Fig. S1**), suggesting that the tissue microenvironment may modulate matrix–fluorophore interactions.

### Vascular lipid compartmentalization mapped at single-vessel resolution

MALDI-MSI acquisition at ∼5 μm lateral spatial resolution revealed distinct phospholipid and sphingolipid distributions within hippocampal artery structures and the surrounding microenvironments (**Fig. 2d**). Phosphatidic acid (PA 18:1/18:0, [M–H]⁻ at *m/z* 701.52) was higher in the artery wall and in the surrounding tissue than in the lumen (**Fig. 2d**, **h**). Phosphatidylinositol (PI; 16:0/22:4, [M–H]⁻ at *m/z* 885.55) exhibited strong enrichment throughout the artery wall^12–14^, with elevated signal in the endothelial layer which colocalized with high CD31 immunofluorescence (**Fig. 2e**).

Sulfated hexosylceramide (SHexCer 18:1;O₂/24:1, [M–H]⁻ at *m/z* 888.63, C24:1 sulfatide) was highest in the tissue surrounding the artery and lower in the artery wall and lumen (**Fig. 2d**, **h**), a distribution in agreement with a perivascular glial or myelin-associated source. Sulfatide is a major myelin lipid, and about 17% of mature myelinating oligodendrocytes in the neocortex, hippocampal CA1 and cerebellar cortex lie in contact with blood vessels, mostly capillaries, as shown in mouse models^15^. All three lipid species (PA, PI, and SHexCer) were identified using negative-ion mode imaging parallel reaction monitoring-parallel accumulation-serial fragmentation (iprm-PASEF), which enabled structural assignments through ion mobility-resolved fragmentation analysis (**Supplementary Fig. S3**).

Complementary experiments using CHCA matrix in positive-ion mode revealed additional lipid compartmentalization within vascular structures (**Supplementary Fig. S1d**). Phosphatidic acid PA(16:0/20:4) ([M+H]⁺, *m/z* 697.48) was lowest in the arterial lumen and higher in the artery wall and the surrounding tissue, whereas sphingomyelin SM(d18:1/20:3) ([M+H]⁺, *m/z* 753.59) was highest in the surrounding tissue, intermediate in the artery wall and lowest in the lumen (adjusted *P* < 0.0001 for each pair, **Supplementary Fig. S1h**). Both lipid species were identified using positive-ion mode iprm-PASEF (**Supplementary Fig. S4**).

Notably, the nuclear stain SYTOX Green was also detected as a potassium adduct ([M+K]⁺, *m/z* 318.11) in the MALDI-MSI data, demonstrating that fluorescent probes themselves can serve as internal landmarks for registration that are detectable by both imaging modalities. Dried-droplet MALDI MS analysis confirmed that this signal originated from the fluorescent dye rather than matrix interference, and its colocalization with nuclear fluorescence (**Supplementary Fig. S1b, d, e** and **Supplementary Fig. S5**) validated MALDI-detected fluorescent probes as internal registration landmarks.

Overlaying the enhanced CD31 immunofluorescence and nuclear staining image with the MALDI-MSI-derived PI(16:0/22:4) ion image demonstrated the precise spatial registration between these two imaging modalities with FluoMALDI imaging (**Fig. 2e**). The merged data revealed PI enrichment specifically within the CD31-positive arterial wall, compared to lower signal in both the artery lumen and the surrounding tissue (**Fig. 2h**).

Mass spectra from this region demonstrated distinct lipid profiles (**Fig. 2f, g**). Signal intensities differed among the three compartments for each lipid (Kruskal–Wallis test, *P* < 0.0001, **Fig. 2h**). In Dunn tests with Holm adjustment, PA(18:1/18:0) was higher in the artery wall than in the lumen (*P* < 0.0001) and did not differ from the surrounding tissue (*P* = 0.84). PI(16:0/22:4) was higher in the artery wall than in the lumen and the surrounding tissue (*P* < 0.0001 for each). SHexCer(18:1;O2/24:1) was highest in the surrounding tissue (*P* < 0.0001 versus wall and lumen), which could be explained with a glial or myelin-associated source^15,16^.

This single-section workflow avoided the registration step between serial sections, and simplified image co-registration. Notably, MALDI-MSI signal integrity was largely preserved between unstained and immunostained arterial regions in positive and negative ion modes, with only slight attenuation observed for PA species in negative ion mode (**Supplementary Fig. S6**). For brain vasculature, this single-specimen precision enabled comparative analysis across hippocampal subregions with their distinct vascular features, enabling future studies into disease-specific alterations of the brain vasculature such as those observed in Alzheimer’s disease and temporal lobe epilepsy.

### HaloTag-FluoMALDI imaging of migrating neural crest cells in chick embryos

Embryonic development requires spatiotemporally coordinated morphogenetic events, and the neural crest exemplifies this paradigm as a transient stem cell population that undergoes epithelial-to-mesenchymal transition (EMT) and extensive migration to build the craniofacial skeleton and peripheral nervous system^17,18^. Dysregulation of neural crest development is at the root of many congenital disorders such as Hirschsprung disease and neuroblastoma^17^. Genetic studies demonstrated that SRY-box transcription factor 10 (SOX10)-driven sphingomyelin phosphodiesterase 3 (*SMPD3*) expression generates the plasma membrane lipid ceramide (Cer) which is functionally required for neural crest EMT^19,20^; however, reliance on pan-ceramide antibodies left both the identity of the relevant ceramide species and their spatial localization unresolved. To simultaneously map genetic cell identities and their lipid composition in developing embryos, we integrated a genetically encoded HaloTag-TMR labeling strategy with MALDI-MSI on the same tissue section within the FluoMALDI microscopy workflow (**Fig. 3a**). The HaloTag system, a self-labeling enzyme tag that forms a covalent bond with synthetic ligands through nucleophilic displacement^21,22^, provided stable, irreversible marking of genetically defined cell populations. Genetic labeling with the HaloTag system in combination with the TMR fluorophore as ligand of choice offers an extended π–electron system capable of sustaining FluoMALDI enhancements. Genetic labeling can also be achieved with fluorescent protein reporters such as tdTomato^23^; however, matrix coating quenched rather than enhanced this fluorescent protein signal (**Supplementary Fig. S7**). We therefore used genetic labeling with a HaloTag expression vector and TMR to directly visualize *SMPD3*-driven ceramide production in neural crest cells.

**Figure 3:**
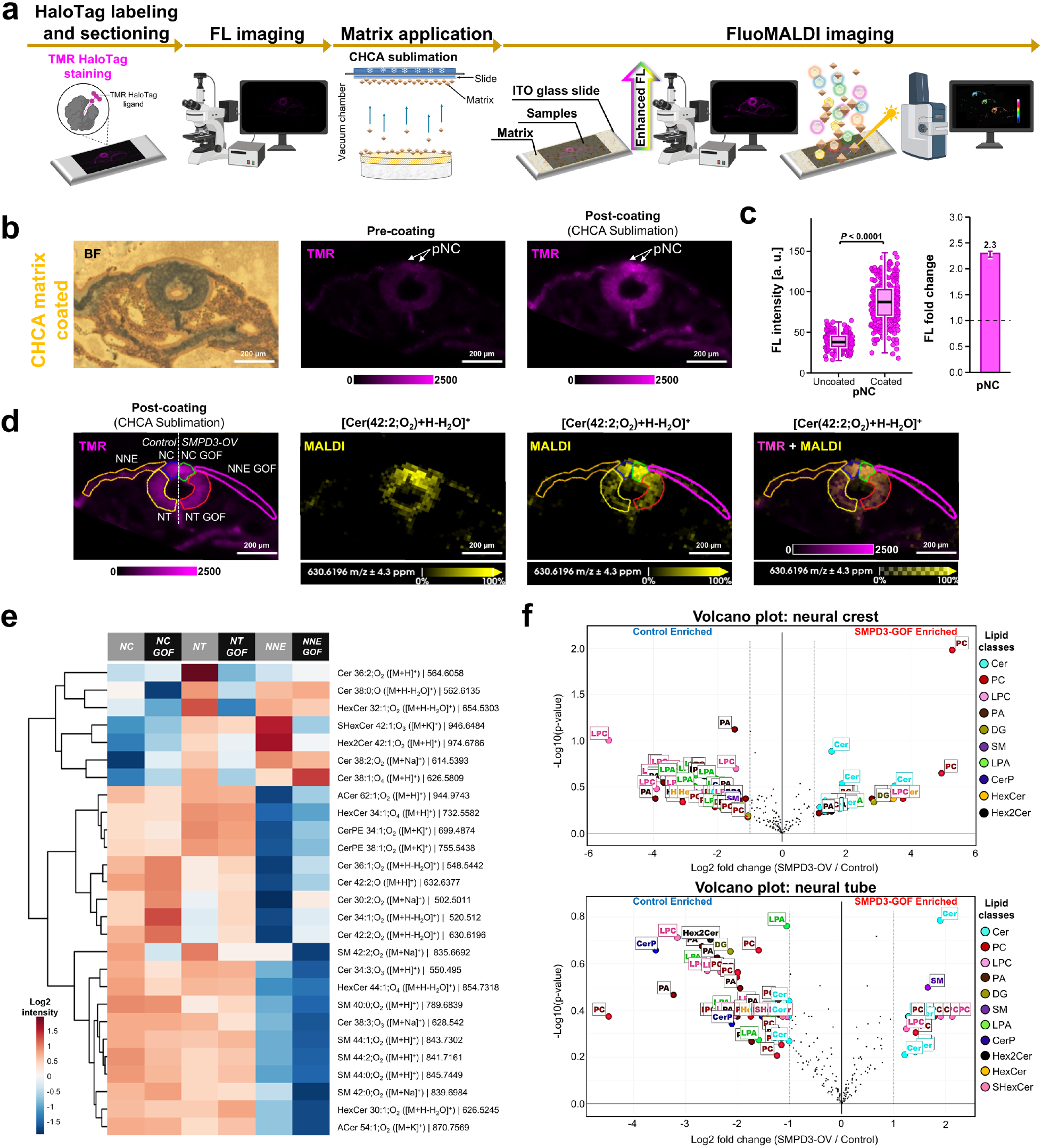
HaloTag-FluoMALDI imaging reveals lipid remodeling in embryonic neural crest cells overexpressing *SMPD3*. (**a**) Workflow for HaloTag-FluoMALDI-MSI analysis. Neural crest cells in the chick embryo were genetically labeled using the HaloTag method. Embryo tissue was sectioned and mounted on slides. Optical microscopy was performed to compare the fluorescence intensity of uncoated and CHCA matrix-coated tissue sections. (**b**) Brightfield (BF) and TMR red fluorescence images (shown in magenta) of HaloTag-TMR-labeled neural crest cells before and after CHCA sublimation coating, demonstrating significant enhancement in the red fluorescence channel. Scale bars: 200 μm. (**c**) Fluorescence intensity (a.u.) of the pre-migratory neural crest (pNC) before and after CHCA sublimation. Boxes show the median and interquartile range, and whiskers show 1.5 times the interquartile range. Statistical comparisons used two-sided Mann-Whitney U tests. The bar shows the fold change in median intensity (coated divided by uncoated, 95% bootstrap confidence interval), and the dashed line marks 1. Exact *P* values and effect sizes are in Source Data. (**d**) HaloTag-TMR fluorescence images illustrating annotated ROI during embryo development: control neural crest (NC) is indicated by the blue line, control neural tube (NT) by the yellow line, control non-neural ectoderm (NNE) by the orange line, *SMPD3*-overexpressing (OV) gain-of-function (GOF) NC by the green line, *SMPD3* GOF NT by the red line, and *SMPD3* GOF NNE by the magenta line. These images are co-registered with the corresponding MALDI images showing ceramide Cer(42:2;O_2_) in yellow. Scale bars: 200 μm. (**e**) Heatmap of Z-scores of log2-transformed MALDI-MSI signal intensities for sphingomyelin (SM) and ceramide (Cer) species within the outlined and labeled tissue regions are shown in rows, following the same color coding of tissue regions as in panel d. Rows represent individual lipid species identified by lipid class, acyl chain composition, detected ion adduct, and *m/z* value and are hierarchically clustered based on abundance patterns. This heatmap highlights the decreased production of SM in the *SMPD3* GOF NC compared to the control. (**f**) Volcano plot showing Cer enrichment in *SMPD3* GOF regions compared to control regions.

*SMPD3* encodes neutral sphingomyelinase 2 (nSMase2), a Mg²⁺-dependent phosphodiesterase that hydrolyzes sphingomyelin to ceramide and phosphocholine at the cytosolic membrane leaflet^24,25^, and is spatiotemporally upregulated at the onset of neural crest EMT, where ceramide production drives endocytosis and activation of Wnt and BMP signaling required for migration^20^. To directly visualize where and to what extent ceramide production is spatially amplified *in vivo*, we expressed a Citrine fluorescent protein fused to the HaloTag7 enzyme (Citrine-HaloTag) under control of the neural crest-specific *TFAP2A E1* enhancer^26^; this strategy allows for restricted expression of the HaloTag enzyme to the neural crest lineage, together with Citrine expression as a live readout of transfection efficiency (**Supplementary Fig. S8**). We performed *ex ovo* electroporation of chicken embryos at Hamburger–Hamilton stage 4 (HH4) to transfect the *TFAP2A E1*::Citrine-HaloTag plasmid together with a gain-of-function (GOF) construct driving a FLAG-tagged nSMase2 enzyme in all transfected cells (*SMPD3-FLAG*) on the embryo’s right side, and the same construct lacking the *SMPD3-FLAG* coding sequence on the left as a contralateral control^22^. Since endogenous *SMPD3* transcription normally initiates at approximately HH8 and remains elevated throughout migratory stages under direct control of the neural crest gene regulatory network^20^, transfection of *SMPD3-FLAG* at HH4 resulted in broader nSMase2 expression ahead of its endogenous onset. After transfection and culturing to specified migratory neural crest stages, we labeled neural crest cells by incubating with the HaloTag TMR ligand live, before fixation and sectioning. Taking advantage of the bilateral symmetry of avian embryos and contralateral delivery system, this approach allows for HaloTag-mediated detection of neural crest cells *in situ*, together with the simultaneous assessment of sphingolipid metabolites in unperturbed conditions (left side) compared to the effects of *SMPD3* GOF on the contralateral side (right side).

Following CHCA matrix sublimation (∼1.5 µg/mm²), TMR-HaloTag fluorescence intensity in neural crest cells increased 2.3-fold compared with pre-coating levels (95% bootstrap confidence interval, *P* < 0.0001, **Fig. 3b,c**). A similar enhancement trend was observed in pure TMR droplets after CHCA sublimation (5-fold; **Supplementary Fig. S8e**), supporting that this fluorescence enhancement was attributable to the matrix coating rather than tissue-specific factors. Following matrix coating, fluorescence imaging was first performed to identify HaloTag-labeled neural crest cells. MALDI-MSI was then conducted on the same section, revealing elevated ceramide Cer(42:2;O_2_) ([M+H−H_2_O]⁺, *m/z* 630.62) levels specifically in the neural crest cell region (**Fig. 3d**), as confirmed by on-tissue MS/MS (**Supplementary Fig. S9**), which is consistent with elevated SMPD3 expression resulting in increased ceramide release^19,20^. Consistent with our immuno-FluoMALDI imaging results, the HaloTag-FluoMALDI-MSI approach demonstrated that fluorescence scanning under standard conditions did not compromise subsequent MALDI imaging quality or spectral integrity. Specifically, Cer and sphingomyelin (SM) signals were well preserved when comparing unstained and HaloTag-stained chick embryo tissue sections analyzed in positive ion mode with CHCA matrix. Only 4 out of 30 lipid species showed a slight decrease in signal intensity, with no statistically significant reduction overall (**Supplementary Fig. S10**).

Spatial lipid analysis revealed various enriched Cer species in pre-migratory neural crest (NC) compared to the adjacent neural tube (**Fig. 3d, e**), consistent with active sphingolipid remodeling at the onset of EMT previously studied^19^. In addition to elevated *SMPD3* expression in the NC^19^, mRNA sequencing data also showed elevated expression of diacylglycerol O-acyltransferase 2 (*DGAT2*; acylceramide (ACer) synthesis), fatty acid synthase (*FASN*), and long-chain fatty acid-CoA ligase 3 (*ACSL3*; myristate production) in NC (**Supplementary Fig. S11**)^19^, linking active lipid metabolic reprogramming to the onset of developmental cell migration.

### SMPD3-mediated sphingolipid remodeling in the neural crest

Consistent with enhanced nSMase2 activity, spatial lipidomics revealed higher ceramide and lower sphingomyelin levels in *SMPD3*-overexpressing NC cells. Heatmap visualization of SM and Cer species revealed a shift from SM to Cer species (**Fig. 3e**). Hierarchical clustering identified a distinct group of ceramide species such as Cer(42:2;O_2_), Cer(34:1;O_2_), and Cer(36:1;O_2_), that exhibited consistently elevated abundance in *SMPD3* GOF regions across the transfected NC, neural tube (NT), and non-neural ectoderm (NNE). This Cer accumulation was accompanied by reciprocal SM depletion, with the most pronounced reduction observed in NC (**Fig. 3e, f**). Individual replicates are shown in Supplementary **Fig. S12**. This reciprocal relationship, in which elevated Cer levels coincide with reduced SM levels, is consistent with enhanced nSMase2 activity in *SMPD3* GOF versus control NC.

Among the ceramide species elevated by *SMPD3* overexpression, Cer(42:2;O₂) (also referred to as C24:1 or Cer d18:1/24:1) showed the most pronounced increase (**Fig. 3d**). This very long-chain monounsaturated Cer, containing the nervonic acid (C24:1 Δ15) acyl chain, exhibited the largest fold-increase (about 4-fold) among ceramide species in *SMPD3* overexpressing NC compared to NC controls (**Fig. 3f**). Interestingly, we also observed phosphatidylcholine species enriched after *SMPD3* overexpression (**Fig. 3f**); since nSMase2-mediated SM hydrolysis releases phosphocholine in addition to Cer, this accumulation may stimulate more PC production accounting for this compositional shift. These findings demonstrate that the HaloTag-FluoMALDI imaging approach enables the detection of spatially targeted perturbation experiments to measure shifts in substrate and product equilibrium of specific cell types within tissue.

### Ectodermal sphingolipid heterogeneity

Spatial lipidomics revealed inherent differences in sphingolipid profiles among NC, NT, and NNE that provide biological context for interpreting *SMPD3* overexpression phenotypes (**Fig. 3d, e**). Even in control embryos, NC and NT cells exhibited elevated Cer intensities compared to NNE cells, consistent with the established requirement for nSMase2 activity during neural crest specification and EMT^20^.

Regional profiling confirmed that NC tissue contains the highest Cer abundance and the lowest SM levels among ectodermal derivatives, consistent with active nSMase2-mediated hydrolysis during EMT (**Fig. 3e, f**). These region-specific lipid profiles are consistent with the membrane remodeling requirements of migratory NC versus non-migratory populations we previously hypothesized from functional perturbation experiments^20^. By enhancing HaloTag-TMR fluorescence while preserving lipid identity within the same section, HaloTag-FluoMALDI imaging links molecular phenotypes to HaloTag-labeled regions of the same section.

### Dye-FluoMALDI and immuno-FluoMALDI imaging of organelles in human iPSCS-derived astrocytes

Astrocytes compartmentalize lipid metabolism at the organelle level, with nuclear sphingolipid signaling regulating apolipoprotein E (ApoE) production, Golgi trafficking building membrane and lipoprotein cholesterol, and actin-associated lipid domains driving process motility for synaptic and vascular ensheathment^27^. Disruption of this organization contributes to disease. Saturated lipid-laden astrocytes turn neurotoxic in neurodegeneration^28^, and lipid-accumulating reactive astrocytes worsen epilepsy^29^. Organelle-resolved lipid imaging is therefore essential to link these subcellular pools to astrocyte dysfunction. We therefore developed FluoMALDI imaging with human iPSC-derived astrocytes cultured directly in chamber slides and labeled with organelle-specific fluorescent probes. Two labeling strategies were employed. For dye-FluoMALDI alone, cells were stained with SYTOX Green to label nuclei and Phalloidin–A635 to label filamentous actin. For combined dye- and immuno-FluoMALDI, a three-probe approach was used consisting of SYTOX Green for nuclei, an A568-conjugated antibody targeting the Golgi apparatus, and Phalloidin–A647 for filamentous actin. Fluorescence images were acquired both before and after CHCA matrix deposition by sublimation, followed by MALDI-MSI acquisition without physical displacement of the sample (**Fig. 4a** and **Supplementary Fig. S13a**). CHCA sublimation produced statistically significant fluorescence enhancement for all probes (*P* < 0.0001 for all comparisons; **Fig. 4b, c** and **Supplementary Fig. S13b, c**). SYTOX Green showed the greatest enhancement, with a 7-fold increase in median fluorescence intensity (95% bootstrap confidence interval, **Fig. 4b,c**). Golgi and actin probes showed more modest enhancements of 1.9-fold and 2.0-fold, respectively.

**Figure 4:**
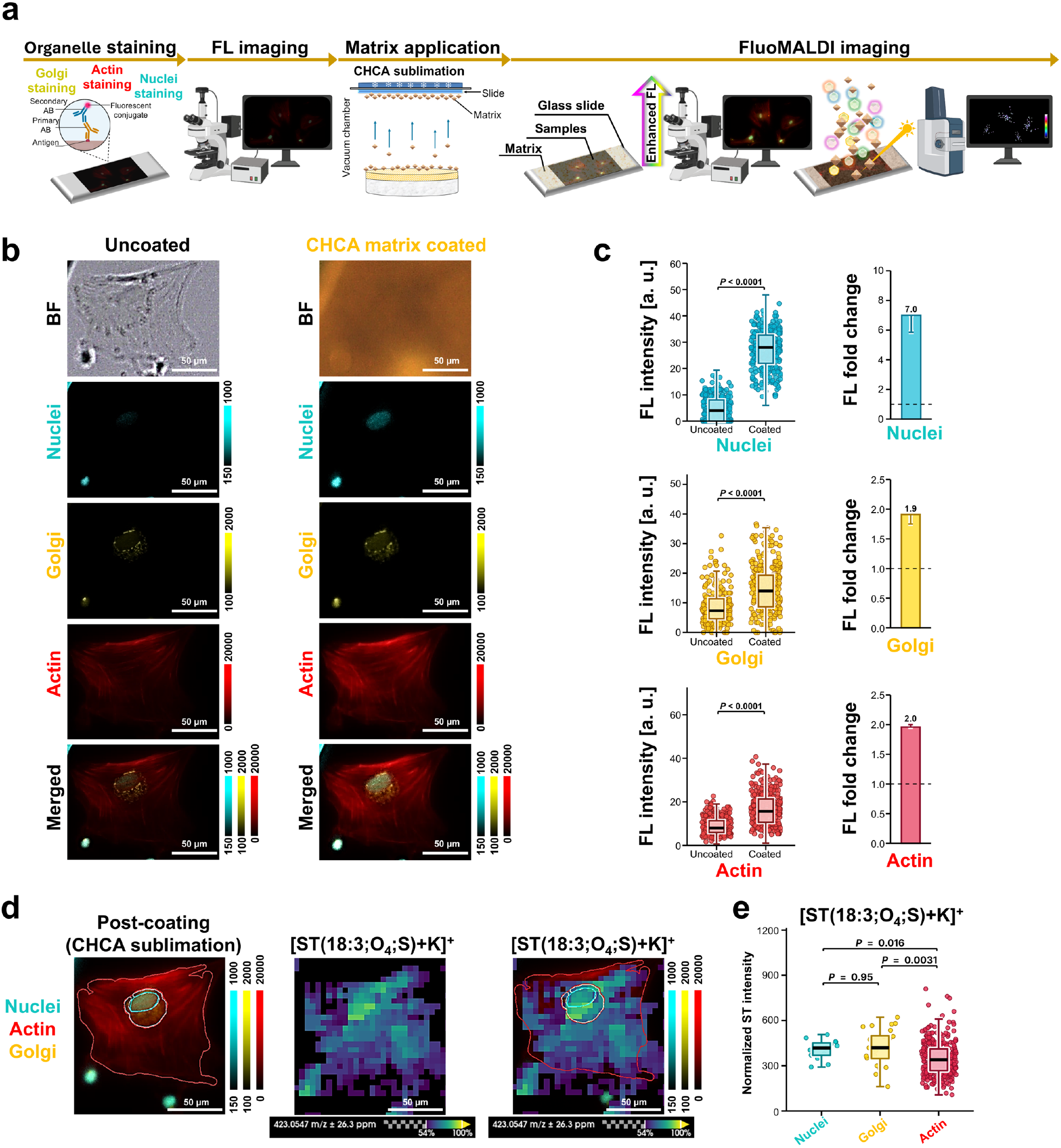
Three-channel dye- and immuno-FluoMALDI imaging of human astrocytes with subcellular organelle staining. (**a**) Workflow for dye- and immuno-FluoMALDI analysis. Human iPSC-derived astrocytes were cultured on chamber slides and labeled with organelle-specific fluorescent dyes. Optical microscopy was performed to compare the fluorescence intensity of uncoated and coated cells. CHCA matrix was applied by sublimation. (**b**) Representative brightfield (BF) and fluorescence images of human astrocytes before (left column, uncoated) and after (right column, CHCA matrix-coated) matrix deposition. Fluorescence channels are displayed as follows: nuclei (SYTOX Green) in cyan, Golgi apparatus (A568-conjugated antibody) in yellow, and F-actin (Phalloidin-Alexa Fluor 647) in red. Bottom row shows merged multichannel images. Scale bars: 50 μm. (**c**) Fluorescence intensity (a.u.) of nuclei, Golgi and actin before and after CHCA sublimation (left) and corresponding fold change (right, ratio of medians, 95% bootstrap confidence interval, the dashed line marks 1). Each point is one pixel. Boxes show the median and interquartile range, and whiskers show 1.5 times the interquartile range. Statistical comparisons used two-sided Mann-Whitney U tests, *P* < 0.0001 for all channels. (**d**) Integrated fluorescence and MALDI-MSI of astrocytes. Left: post-coating fluorescence image showing subcellular compartments (nuclei in cyan, F-actin in red, Golgi in yellow). Center: MALDI ion image of sulfated C18 steroid ST(18:3;O_4_;S) at *m/z* 423.0547 ([M+K]⁺). Right: co-registered overlay of fluorescence microscopy data (subcellular boundaries indicated by colored outlines) with MALDI-MSI data. (**e**) Normalized MALDI-MSI intensity of ST(18:3;O4;S) in the nuclei, Golgi and actin regions defined by the co-registered fluorescence data. Boxes show the median and interquartile range, and whiskers show 1.5 times the interquartile range. Kruskal–Wallis test, *P* = 0.00021, followed by two-sided Dunn tests with Holm adjustment (adjusted *P* shown). Effect sizes are in Source Data.

Parallel experiments using sublimated nH on identically stained astrocytes produced comparable organelle-specific signal amplification (**Supplementary Fig. S14** and **Supplementary Fig. S15**). Nuclei fluorescence exhibited approximately 5-fold enhancement, while Golgi apparatus fluorescence showed 2.7- to 7.8-fold signal increases depending on the Alexa fluorophore used (A568 or A488, respectively). Similarly, fluorescently labeled actin filaments demonstrated a 1.9- to 2.7-fold amplification depending on the fluorophore used (A647 or A635, respectively). Enhancement was further validated in isolated dye preparations, where purified dried droplets of cytoskeletal probes (Phalloidin–A635/647), immunofluorescence conjugates (anti-mouse A488/568), and organelle-dye (SYTOX Green) all showed substantial fluorescence amplification following matrix sublimation (**Supplementary Fig. S16–S18**), indicating that the phenomenon arises from intrinsic dye–matrix photophysical interactions rather than cellular context-dependent effects.

Co-registered MALDI-MSI of the same field of view revealed the spatial distribution of the putative sulfated C18 steroid species ST(18:3;O_4_;S), detected as its potassium adduct [M+K]⁺ at *m/z* 423.0547 (**Fig. 4d**). We also detected several ceramide species, including Cer, acyl ceramides (ACer), and ceramide phosphates (CerP), which were detected as proton adducts [M+H]⁺ (**Supplementary Fig. S13d**). Overlay of the MALDI ion image with fluorescence-defined organelle boundaries revealed a spatially heterogeneous ST(18:3;O_4_;S) distribution within individual astrocytes (**Fig. 4d**, right). Quantitative analysis of normalized MALDI signal intensities from nuclei-, Golgi- and actin-defined regions of interest showed that ST(18:3;O4;S) differed among regions (Kruskal–Wallis test, *P* = 0.00021). In Dunn tests with Holm adjustment, the signal was lower in the actin-defined region than in the nuclear region (*P* = 0.016) and the Golgi region (*P* = 0.0031), with a median about 17% lower than nuclei and 19% lower than Golgi, and it did not differ between the nuclear and Golgi regions (*P* = 0.95) (**Fig. 4e**). The subcellular resolution afforded by dye- and immuno-FluoMALDI imaging allowed MALDI-MSI signals to be assigned to fluorescence-defined regions of the same cell. ST(18:3;O4;S) was relatively depleted in the actin-defined cell region compared with the nuclear and Golgi regions, with the limitation that the actin region is the cell area outside the nucleus and Golgi and that MALDI pixels are similar in size to these organelles.

Since fluorescence microscopy resolves subcellular structures that remain below the effective spatial resolution of current commercial MALDI-MSI instruments, dye- and immuno-FluoMALDI enable direct spatial assignment of lipid signals to physically co-registered organelle boundaries within the same tissue section, eliminating reliance on computational inference. This approach enables nucleus, Golgi, and actin-region-assigned lipid measurements in single cells on commercially available instrumentation. This capability matters because organelle-specific lipid dysregulation, including Golgi fragmentation in Alzheimer’s disease and amyotrophic lateral sclerosis, is increasingly being recognized as a driver rather than a consequence of neurodegeneration^30^.

### Crystal packing of matrix–fluorophore co-crystals

To probe the structural origins of MALDI matrix-enhanced fluorescence, we selected rhodamine B and A488 for co-crystallographic analysis with a MALDI matrix because they represent the two foundational xanthene-derived fluorophore lineages used throughout this study. Rhodamine B serves as a structural analog to the TMR framework utilized for HaloTag labeling^21,22^, while A488 is a sulfonated derivative of rhodamine 110, structurally representative of the Alexa Fluor dyes (A488, A568) deployed for immunolabeling^31,32^. Both are long-established, extensively characterized xanthene fluorophores that provide a well-studied photophysical reference for interpreting the resulting co-crystal structures^33^.

Single-crystal X-ray structures of co-crystallized matrix–fluorophore pairs were experimentally determined to characterize the molecular packing arrangement of CHCA with rhodamine B, and nH with A488 (**Fig. 5** and **Supplementary Fig. S19,** and **S20**). Both co-crystals contained localized, finite π–π interactions rather than an extended, repeating π–π stack (**Fig. 5**). In the CHCA–rhodamine B co-crystal, rhodamine B molecules formed a parallel π–π interaction with each other (interplanar distance 3.647 Å; 0° interplanar angle), and CHCA molecules independently formed a separate parallel π–π interaction (3.415 Å; 0°) (**Fig. 5a, b**). CHCA did not form π–π interactions with rhodamine B and instead formed hydrogen-bonds to bridge the rhodamine B pairs (**Supplementary Fig. S20a)**. The nH–A488 crystal contained two geometrically distinct types of A488···nH π–π interactions. A near-parallel interaction (interplanar angle 6.3°) occurred at each end of a finite A488···nH···nH···nH···nH···A488 motif, and a separate, more inclined interaction (21.2°) occurred elsewhere in the structure (**Fig. 5c** and **Supplementary Fig. S20c**). In the motif, successive interplanar angles are 6.3°, 5.6°, 0°, 5.6° and 6.3°, and the two central, parallel nH molecules are 3.313 Å apart (**Fig. 5d**). π–π interactions in both co-crystals were stabilized by an accompanying hydrogen-bonding network (**Supplementary Videos S1, S2** and **Supplementary Table S1** and **S2**). Single-component crystals of nH and CHCA without fluorophore showed no comparable π–π interactions (**Supplementary Videos S3, S4)**, indicating that these interactions require the presence of the fluorophore and co-crystallization, rather than reflecting an intrinsic matrix property.

**Figure 5:**
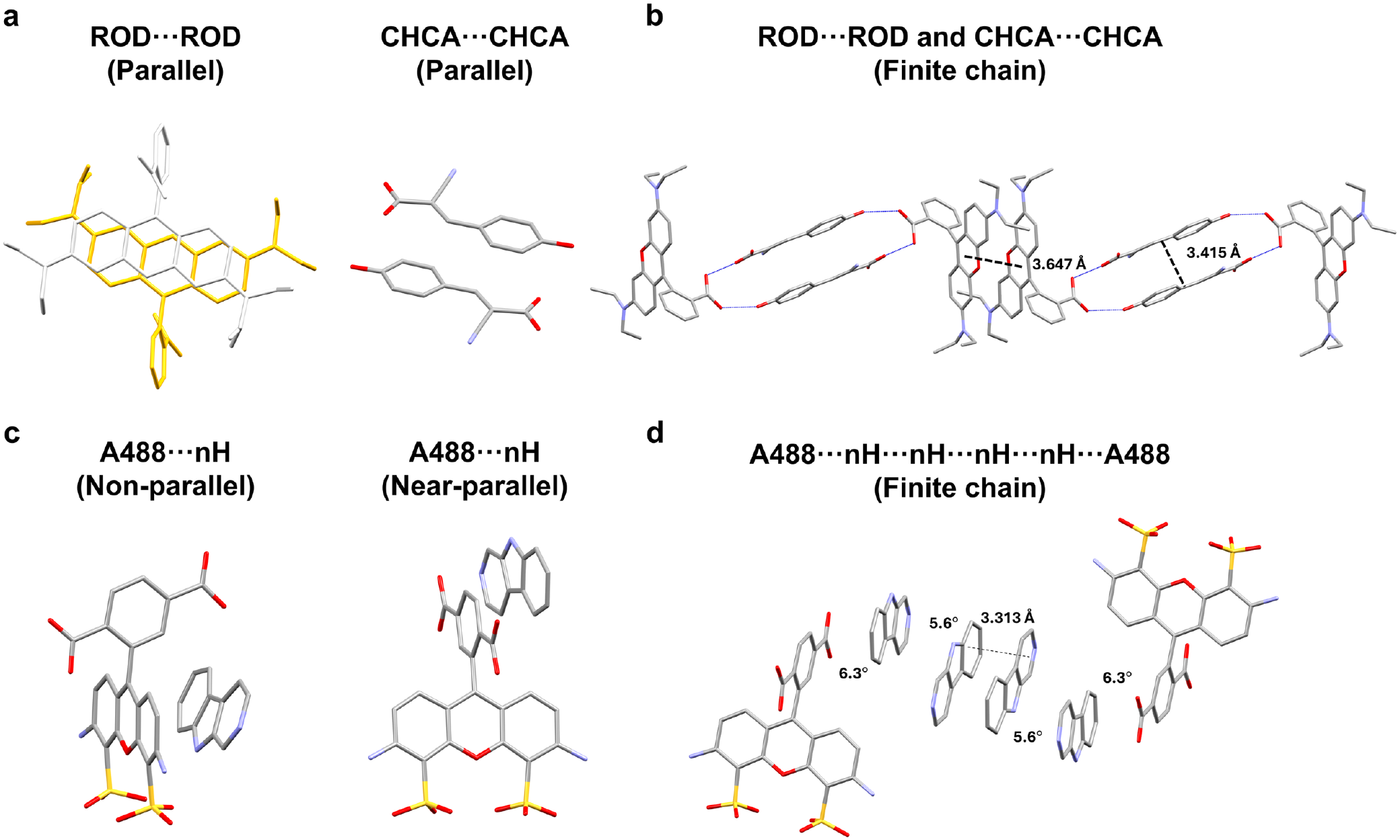
Quantitative geometry of π–π interactions in matrix–fluorophore co-crystals. (**a**) CHCA–rhodamine B (ROD) co-crystal showing parallel ROD···ROD and CHCA···CHCA π–π interactions, with an interplanar angle of 0° for both contacts. (**b**) Offset view of the ROD···ROD and CHCA···CHCA interactions, showing interplanar distances of 3.647 Å and 3.415 Å, respectively. (**c**) nH–A488 co-crystal showing two geometrically distinct A488···nH π–π interactions, a non-parallel interaction with an interplanar angle of 21.2° and a near-parallel offset interaction with an interplanar angle of 6.3°. (**d)** Complete finite A488···nH···nH···nH···nH···A488 motif, showing an interplanar distance of 3.313 Å between the two central, parallel nH molecules and interplanar angles of 6.3°, 5.6°, 0°, 5.6°, and 6.3° between successive molecular planes. The motif terminates at both ends and does not propagate through the crystal lattice. Disorder, hydrogen-bond interactions, hydrogen atoms, and lattice water molecules are omitted for clarity.

### MALDI matrix effects on fluorescence scattering

Because matrix deposition occurs directly on the specimen and could potentially scatter light and broaden the point spread function (PSF), potentially resulting in blurring of fluorescence images, we measured the full width at half maximum (FWHM) of sub-resolution fluorescent beads before and after MALDI matrix coating to test whether the resolution needed to observe subcellular structures is preserved. PSF measurements using 200-nm-diameter fluorescent microspheres assessed whether matrix application compromised the spatial resolution of fluorescence imaging. FWHM values were quantified across green, red, and far-red channels before and after sublimation of CHCA or nH at exposure times of 20 and 50 ms. Fluorescent beads remained clearly resolved across all spectral channels following matrix application (**Supplementary Fig. S21**). Post-matrix FWHM values were 0.46 ± 0.08 µm (green), 0.48 ± 0.08 µm (red), and 0.54 ± 0.04 µm (far-red), corresponding to increases of 4.9–49.5% relative to pre-matrix conditions (n = 1646, 1021, and 1950 beads per channel, respectively; mean ± s.d.). Although matrix-induced light scattering did slightly increase apparent bead diameter, subcellular morphology was largely preserved, and organelle boundaries remained clearly distinguishable in post-coating fluorescence images (**Fig. 4b, Supplementary Fig. S13–15**).

## Discussion

We developed FluoMALDI imaging, in which matrix deposited prior to fluorescence imaging enhances, rather than quenches fluorescent signals, enabling amplified co-registered fluorescence and MALDI-MSI from the same tissue section and cells. Since both modalities interrogate the identical physical substrate, image co-registration reduces to a simple rigid transformation rather than deformable warping^3,4^, eliminating the geometric distortions inherent to serial-section workflows^1,2^. This allows molecular profiles to be extracted directly from genetically labeled cells, immunolabeled structures, or organelle-delimited regions, shifting spatial lipidomics from region-level to compartment- and organelle-resolved analysis, anchored to verified gene or protein expression^21^. Validated across dye-, immuno-, and HaloTag-FluoMALDI imaging, the FluoMALDI-MSI approach collectively spans three widely used fluorescent labeling strategies used in cell, developmental, and neurobiology studies. This establishes FluoMALDI imaging as a broadly applicable platform for spatial omics, which allows for hypothesis-driven biological investigations that require selective labeling of cell types, tissue structures, gene expression states, or organelles.

The biological applications presented here illustrate the capabilities of FluoMALDI imaging to generate insights inaccessible to either modality alone. Immuno- and dye-FluoMALDI mapping of human hippocampal vasculature revealed compartment-specific lipid distributions at single-vessel resolution, with PA(18:1/18:0) enrichment in arterial walls and SHexCer(18:1;O_2_/24:1), also known as C24:1 sulfatide, highest in the perivascular region. The spatial agreement between SHexCer(18:1;O_2_/24:1) and perivascular zones is consistent with the established association of oligodendrocyte processes with vascular basement membranes, as demonstrated quantitatively in mouse neocortex where approximately 17% of mature myelinating oligodendrocytes maintain direct vascular contact^15^. Positive-ion mode acquisition with CHCA matrix revealed complementary sphingolipid and glycerophospholipid partitioning within vascular structures. SM(d18:1/20:3) and PA(16:0/20:4) were both lowest in the arterial lumen, and PA(18:1/18:0) was also lower in the lumen than in the artery wall in negative-ion mode. These data show that the lipid composition of the lumen differs from that of the wall and the surrounding tissue.

Immuno-FluoMALDI imaging was able to resolve lipid distributions at the neurovascular interface and combined with enhanced fluorescence imaging and detailed morphological visualization, provides a valuable tool for investigating lipid-mediated mechanisms underlying selective neuronal vulnerabilities. Immuno-FluoMALDI imaging leverages standard fluorophore-conjugated antibodies rather than specialized barcoded or isotope-tagged reagents, allowing direct integration into existing workflows. Given the widespread use of immunofluorescence in research and development, immuno-FluoMALDI imaging allows endogenous lipids and metabolites to be mapped directly to protein-defined structures on the same section using standard antibody-based workflows, a capability that previously required separate specimens or dedicated multiplex platforms.

HaloTag-FluoMALDI imaging allowed us to investigate neural crest cells within chick embryos and revealed direct spatial evidence that nSMase2 activity from elevated *SMPD3* expression in overexpression experiments remodeled the cellular sphingolipid composition in the neural crest. Cer species were elevated and SM species were reduced in *SMPD3* overexpressing neural crest cells, with Cer(42:2;O₂), a very long chain ceramide comprising 42 total carbons and two degrees of unsaturation, as the predominantly enriched product. This reciprocal shift is consistent with nSMase2’s broad substrate specificity toward SM species across a range of N-acyl chain lengths, as supported by the reduction of Cer species in *SMPD3*-deficient models^24^. Long chain ceramides promote membrane curvature through acyl chain interdigitation and drive the formation of Cer-enriched platforms that cluster signaling receptors, thereby enhancing endocytosis-mediated Wnt and BMP pathway activation during EMT^20,24^. Genetically engineered fluorescent reporters, including chemigenetic systems such as HaloTag, are widely used in protein localization, protein function, and gene promoter activity studies in developmental and cell biology^9^. HaloTag-FluoMALDI imaging extends HaloTag-based approaches by directly mapping lipids to genetically defined cell populations on the same tissue section.

By combining dye-FluoMALDI (SYTOX Green for nuclei, phalloidin for actin) with immuno-FluoMALDI targeting of the Golgi apparatus, we extended hypothesis-driven spatial analyte mapping to the subcellular level in cultured human iPSCs-derived astrocytes. Fluorescent organelle labeling with dye- and immuno-FluoMALDI revealed a higher abundance of the sulfated C18 steroid species ST(18:3;O_4_;S) in the Golgi and nucleus as compared to the actin-delineated peripheral membrane domains, with no significant difference detected between the Golgi and nuclear compartments. Astrocytes synthesize neuroactive steroids, including estradiol and progesterone^34,35^, which influence neuronal excitability and synaptic plasticity. Moreover, progesterone-derived neurosteroids such as allopregnanolone exert modulatory effects on GABA_A_ receptors.

Steroid hormones are synthesized predominantly in peripheral steroidogenic tissues. However, the brain is also capable of de novo neurosteroid synthesis, and steroid activity and bioavailability can be further regulated by reversible sulfation and desulfation^36^. Cytosolic sulfotransferases catalyze steroid sulfation, whereas steroid sulfatase (STS), an endoplasmic reticulum (ER)-associated enzyme, catalyzes the hydrolysis of steroid sulfates, providing a mechanism for regulating the balance between sulfated and unconjugated steroid species^35–37^. In our spatial lipidomic analysis, the sulfated C18 steroid species ST(18:3;O_4_;S) was relatively depleted from actin-delineated peripheral cellular domains but showed comparatively greater abundance in Golgi-associated regions. This spatial distribution suggests compartmentalized handling of this sulfated steroid species within astrocytes and may reflect intracellular trafficking, metabolism, storage, or secretory processing associated with steroid homeostasis. Coupling organelle-specific fluorescent markers with FluoMALDI-MSI in the same astrocyte preparation enabled us to map lipid distributions to morphologically defined astrocytic domains. Moreover, the successful application of dye- and immuno-FluoMALDI imaging to astrocytes, using organelle-specific probes targeting the nucleus, Golgi apparatus, and actin cytoskeleton demonstrates the versatility of this approach for spatial lipidome mapping across diverse neural cell populations.

The localized π–π interactions between two fluorophore molecules or a fluorophore and a matrix molecule in MALDI matrix-fluorophore co-crystals, which were identified by single-crystal X-ray crystallography, likely contribute to the excitonic coupling associated with the observed fluorescence enhancement. These results are consistent with aromatic π–π interactions, which are not arranged in an extended stacking pattern in these co-crystals, not inherently quenching fluorescence, with the photophysical outcome instead depending on the specific supramolecular geometry and coupling regime of the interacting fluorophores^38^. Rhodamine B and A488 each formed close π–π interactions upon co-crystallization with MALDI matrices, whereas matrix-only crystals showed no comparable π–π interactions, implicating co-crystallization itself, rather than a specific fluorophore–matrix pairing, as the structural trigger for fluorescence enhancement. Analysis of matrix-induced fluorescence scattering revealed unchanged PSF symmetry and centroid position following matrix application, confirming that matrix deposition preserves the spatial fidelity needed for subcellular FluoMALDI imaging.

Systematic evaluation of matrix and fluorophore combinations revealed that co-crystallization-mediated enhancement is not limited to free dyes. Fluorophores conjugated to antibodies (∼150 kDa) and HaloTag (∼33 kDa) retained enhancement despite substantial steric constraints, which likely indicates that matrix access to the fluorophore’s aromatic core contributes to the FluoMALDI effect. Optimal matrix choice nonetheless depended on fluorophore identity and imaging context. While nH outperformed CHCA for A647 in dried droplets, this relationship reversed on tissue sections, where CHCA provided superior enhancement for both A647 and SYTOX Green. This reversal may reflect tissue-specific effects on co-crystallization dynamics. Taken together, these results provide a first matrix and fluorophore matching framework, analogous to established matrix selection principles for different MALDI-MSI analyte classes, to guide future FluoMALDI-MSI experimental design.

Emerging in-source fluorescence, transmission-geometry MALDI imaging platforms achieve MSI pixel sizes below 1×1 µm² by integrating a fluorescence microscope, which shares the same optical path and coordinate system with their transmission-mode laser optics for MALDI imaging, for direct image co-registration^39^. A related transmission-geometry ambient ionization platform has pushed lipid imaging to sub-micron spatial resolutions^40^. The fluorescence signal enhancement afforded by matrix-driven co-crystallization in FluoMALDI offers a practical route for improving the precision of fluorescence landmark localization in in-source fluorescence MALDI imaging platforms, increasing co-registration fidelity. Independent of ongoing instrumentation advances, the cross-system biological insights presented here, spanning neurovascular lipid architecture, developmental ceramide-species dynamics, and subcellular steroid localization, demonstrate that hypothesis-driven integration of morphology and molecular composition can uncover previously inaccessible spatial biology. By combining hypothesis-driven fluorescent labeling of defined cell populations, anatomical structures, and subcellular organelles with spatially resolved molecular imaging, FluoMALDI imaging places omics measurements within a biologically meaningful spatial context. Leveraging the intrinsic fluorescence-enhancing effects of matrix co-crystallization, FluoMALDI imaging provides a streamlined, single-specimen approach to high-resolution spatial omics that is broadly accessible to laboratories with conventional fluorescence microscopy and MALDI-MSI capabilities. This single-specimen efficiency is particularly valuable for irreplaceable biological materials, including archival human tissue, patient derived iPSC lines, and genetically defined embryos that often cannot be recollected. In contrast, serial-section workflows consume limited tissue samples across multiple sections and may fail to capture small, discontinuous structures, such as individual capillaries or migratory cells that are absent from adjacent sections. By extracting morphological and molecular information from the same tissue section, FluoMALDI imaging maximizes the information obtained from scarce specimens, preserving precious material for replication and follow-up studies rather than parallel serial-section analyses.

## Online Methods

### Chemicals and materials

All solvents and trifluoroacetic acid (TFA) were purchased from Sigma-Aldrich (St. Louis, MO) unless otherwise noted. Norharmane (nH), α-cyano-4-hydroxycinnamic acid (CHCA), glycine, and goat serum were also obtained from Sigma-Aldrich. Paraformaldehyde (PFA), donkey anti-goat IgG (H+L) highly cross-adsorbed secondary antibody conjugated to Alexa Fluor™ Plus 647, goat anti-mouse IgM (heavy chain) cross-adsorbed secondary antibody conjugated to Alexa Fluor™ 568, goat anti-mouse IgG (H+L) Alexa Fluor 488, SYTOX™ Green, Alexa Fluor™ 488, phalloidin labeling probes (Alexa Fluor^TM^ 635 and 647), and FluoSpheres™ Carboxylate-Modified Microspheres (yellow-green, red, dark red) were purchased from Thermo Fisher Scientific (Waltham, MA). The primary anti-CD31 antibody was obtained from R&D Systems (Minneapolis, MN), and the anti-GM130 antibody was obtained from Abcam (Cambridge, MA). Citrate buffer was purchased from Biolegend (San Diego, CA). Purified recombinant tdTomato (Discosoma sp. red fluorescent protein) containing an N-terminal His tag and expressed in *E. coli* was obtained from OriGene (Rockville, MD). HaloTag® biotin tetramethylrhodamine (TMR) ligand was purchased from Promega (Madison, WI). Eight well chamber slides were purchased from ibidi (Fitchburg, WI). All solvents and reagents, which were of analytical grade, were employed without further purification.

### Human brain tissue collection and immunofluorescence staining

Postmortem human brain tissue was obtained according to protocols approved by the JHU Institutional Review Board and processed following established neuropathological protocols as previously reported^41^. Upon autopsy, brains were hemisected along the sagittal plane, with one hemisphere fixed in 10% neutral buffered formalin (NBF; pH 7.4, Sigma-Aldrich, cat. no. HT501128) for subsequent histological examination, and the contralateral hemisphere was cut into coronal slabs. Tissue blocks of selected brain regions including hippocampus were snap-frozen in liquid nitrogen and stored at -80°C for molecular analysis. The remaining tissue slabs were frozen on chilled aluminum plates and dry ice and stored at -80°C. Frozen hippocampal samples were cryosectioned at 20 µm thickness using a CM1860 UV Cryostat (Leica Biosystems, Wetzlar, Germany) and thaw-mounted onto poly-L-lysine-coated indium tin oxide (ITO) slides (Delta Technologies, Loveland, CO). Sections were stored in vacuum-sealed slide mailers at -80 °C until use.

For immunofluorescence^7^, sections were fixed in freshly prepared 4% PFA for 15 min at room temperature, subjected to antigen retrieval in 10 mM citrate buffer (pH 6.0, 95 °C, 1h), and washed in 0.01 M phosphate-buffered saline (PBS, pH 7.2). Subsequently, the sections were blocked in a solution containing 20% goat serum, 1% bovine serum albumin (BSA), 0.3 M glycine, and 0.01 M PBS (pH 7.2) for 24 h at 4 °C. The sections were then incubated with primary anti-CD31 antibody (1:20; R&D Systems, Minneapolis, MN) for 24 h at 4 °C, followed by donkey anti-goat IgG (H+L) secondary antibody conjugated to Alexa Fluor™ Plus 647 (1:100; Invitrogen, Carlsbad, CA) for 3 h at 37 °C. Nuclei were counterstained with 1 μM SYTOX Green for 2 min, and sections were washed in PBS for 5 min with shaking at 70 rpm.

### Construct design and cloning

PCR amplification and NEBuilder Hi-Fi DNA Assembly (NEB) were used to produce an in-frame fusion of Citrine fluorescent protein with the HaloTag7 coding sequence (synthesized by GeneArt, Thermo Fisher Scientific), separated by an -SGGGGS-linker sequence, and assembled into the pTK plasmid harboring the chicken *TFAP2A E1* enhancer sequence to drive neural crest-specific expression^26^. The resulting pTK *TFAP2A E1*::Citrine-HaloTag constructs were verified by nanopore sequencing (Plasmidsaurus, Louisville, KY) before endotoxin-free plasmid preparation (Zymo Research, Irvine, CA) and injection.

### Chicken embryology and sample preparation

Fertilized white leghorn chicken eggs were purchased commercially (University of Connecticut Poultry Unit, Storrs, CT). Eggs were cultured at 38°C in humidified incubators to reach Hamburger-Hamilton-stage 4 (HH4), then harvested for *ex ovo* culture and electroporation as previously described^22^. Endotoxin-free plasmid DNA constructs were injected, and the embryonic epiblast was transfected by electroporation using five pulses of 5.8 V for 50 ms at 100 ms intervals. Plasmids were injected at 2 µg/µl concentrations for pCI H2B-RFP and pCI SMPD3-FLAG^20,42^, and 3 µg/µl concentration for pTK *TFAP2A E1*::Citrine-HaloTag. Embryos were then cultured in albumin to the desired stages, at which point embryos with strong electroporation efficiency were selected for processing. Embryos were washed twice in Ringer’s solution, then stained for 1 hour at 37°C with 5% CO_2_ in 5 µM HaloTag TMR Ligand (Promega, Madison, WI) solution in DMEM supplemented with 10% Fetal Bovine Serum and 1% penicillin and streptomycin. Stained embryos were then washed twice with Ringer’s solution, fixed for 15 min with 4% PFA solution in PBS, rinsed, then flash frozen for 5 min dry in cryomolds on the cryobar in an HM525 NX cryostat (Epredia, Kalamazoo, MI), then stored at -80°C. For cryosectioning, molten gelatin solution (7.5% gelatin and 15% sucrose in PBS) was added to the top of each embryo on the cryobar and allowed to freeze for 5 min, then transferred to -80°C overnight. The following day, gelatin-covered embryo blocks were sectioned at an 18 µm thickness onto poly-L-lysine-coated ITO slides, then stored under vacuum at -20°C until processed for FluoMALDI-MSI analysis.

### iPSCs differentiation to astrocytes

Human astrocytes were differentiated from a control iPSC line as previously reported^43^. Neural progenitor cells (NPCs) were seeded onto Matrigel-coated culture plates at a density of 15,000 cells cm⁻² and cultured in astrocyte differentiation medium (ScienCell Research Laboratories, Carlsbad, CA, cat. no. 1801) supplemented with 2% fetal bovine serum and astrocyte growth supplement (ScienCell Research Laboratories, cat. no. 1852). Cells were maintained under these conditions for 30 days with medium replacement every other day. After 30 days of differentiation, astrocytes were passaged at a ratio of 1:3 and further expanded under the same culture conditions for up to 120 days.

### Fluorescence labeling of iPSCs-derived astrocytes

Astrocytes were seeded into 8-well chamber slides at a density of 1 × 10⁴ cells per well and cultured until reaching an appropriate confluence. For immunofluorescence staining, cells were washed three times with PBS and fixed with 4% (w/v) PFA in PBS (pH 7.4) for 15 min at room temperature. Fixed cells were washed three times with PBS and permeabilized with 0.2% (v/v) Triton X-100 in PBS for 5 min. Non-specific binding was blocked by incubation with 5% (v/v) goat serum in PBS for 1 h at 4 °C.

For Golgi apparatus visualization, samples were incubated overnight at 4 °C with primary anti-GM130 antibody (Abcam, ab169276; 1:400 dilution) to label a Golgi marker. Following primary antibody incubation, cells were washed three to four times with PBS and incubated with secondary antibodies Alexa Fluor 488 goat anti-mouse IgG (Invitrogen, A11001; 1:250 dilution) or Alexa Fluor 568 goat anti-mouse IgM (Invitrogen, A21043; 1:250 dilution) for 1 h at room temperature. Filamentous actin was visualized using Alexa Fluor 635- or Alexa Fluor 647-conjugated phalloidin (1:100 dilution), and nuclei were counterstained with SYTOX Green (1 μM).

Following staining, all samples were washed two to three times with PBS. Residual buffer was removed, and samples were dried for 1 min under a stream of nitrogen gas. To preserve cellular architecture, samples were snap-frozen on a freezing boat floated on liquid nitrogen. Cell slides were stored in vacuum-sealed slide mailers at -80°C for no more than one week prior to FluoMALDI-MSI analysis.

### MALDI matrix application

Slides were equilibrated to ambient temperature in a vacuum desiccator before matrix application. For sublimation, matrix was deposited using the HTX SubliMATE^TM^ system (HTX Technologies, Chapel Hill, NC) with 15 mg of nH or 20 mg of CHCA. The SubliMATE^TM^ lid was pre-cooled with an ice pack to maintain the slide surface at reduced temperature. Preheating temperature was 185 °C for CHCA, and sublimation heating temperatures were 200 °C for nH and 300 °C for CHCA at 0.40 torr. Sublimation proceeded for 5 min. For spray deposition, CHCA or nH matrix (10 mg/mL in 70% acetonitrile (ACN) with 0.1% TFA) was applied uniformly using an HTX M3+ sprayer (HTX Technologies) with the following parameters: temperature of 70 °C, nozzle velocity of 1,000 mm/min, flow rate of 60 µl/min, 3 passes, track spacing of 2 mm, drying time of 2 s between passes, nitrogen gas pressure of 10 psi. Matrix coating density was calculated automatically by the sprayer software as:

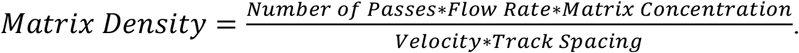

### Fluorescence image acquisition and analysis

Fluorescence images were acquired before and after matrix deposition using slide-scanning fluorescence microscopy (SSFM) with an Olympus Slideview VS200 slide scanner (Evident Corp., Waltham, MA) at the Johns Hopkins Applied Imaging Mass Spectrometry (AIMS) Core facility.

Three filter sets were used: FITC (474/27 nm excitation, 525/45 nm emission), TRITC (545/23 nm excitation; 609/54 nm emission), and Cy5 (635/18 nm excitation; 680/42 nm emission). All images were acquired with a 20× objective lens. For pre- and post-matrix coating comparisons, a fixed exposure time of 1 ms was applied across all fluorescence channels. For cell imaging, channel-specific exposure times were set as follows: 30 ms for Cy5 (actin), 10 ms for TRITC (Golgi), 10 ms for FITC (Golgi), and 1 ms for FITC (nuclei). This standardized acquisition protocol facilitated high-resolution fluorescence imaging across multiple spectral channels.

Fluorescence images were processed using Fiji (ImageJ)^44^. For dried fluorophore droplet experiments, circular regions of interest (ROIs) were drawn within each droplet interior on corresponding brightfield images, excluding edge drying artifacts. For tissue and cell experiments, pixel-wise fluorescence intensities with spatial coordinates were extracted from each ROI using a custom JavaScript macro in Fiji. For Fig. 2c, the artery region was outlined on the CD31-IF image, and the same ROI was applied to the uncoated and coated images. Box plots with individual pixel values were generated in R.

Fluorescence images are displayed with identical, linear display ranges for uncoated and coated conditions within each figure panel, as indicated by the colorbars. No gamma adjustment, background subtraction, deconvolution, denoising or non-linear contrast change was applied to any image. The term enhancement refers to the matrix-induced increase in fluorescence intensity and not to digital image processing.

### Quantification of Fluorescence Intensity

Fluorescence intensity was measured within defined ROIs and compared between matrix-coated and uncoated control conditions. Mean pixel fluorescence intensity (MPFI) was calculated for each ROI, and fold change was quantified as previously published^4^:

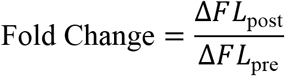

where:

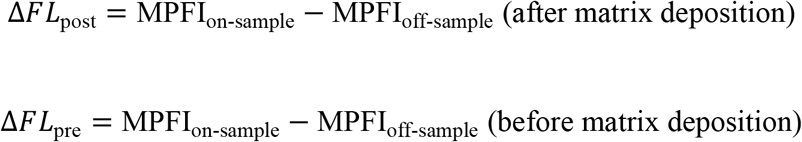

In these equations, MPFI represents the mean pixel fluorescence intensity expressed in arbitrary units (a.u.), on-sample (e.g., on-fluorophore droplet) refers to the ROI positioned on the sample (e.g., droplet), and off-sample (e.g., off-droplet) refers to the ROI positioned outside the sample (e.g., droplet) area for background correction (**Supplementary Figs. S2, S7, S8e, S16–S18**). This approach enabled precise quantification of fluorescence intensity changes induced by matrix deposition while accounting for background signal variations.

For pixel-matched comparisons of fluorescence intensities between uncoated and matrix-coated conditions (**Figs. 2**–**4** and **Supplementary Figs. S1, S13–S15**), pixel-wise fluorescence intensities with spatial coordinates were extracted from the same ROI in the uncoated and coated images using a custom JavaScript macro in Fiji. Because the images were registered and every pixel was measured before and after coating, the two conditions were compared pixel by pixel with a two-sided Wilcoxon signed-rank test. The pixels of the two conditions were treated as independent samples and compared with a two-sided Mann–Whitney U test. Fold change was calculated as the ratio of the median fluorescence intensity of coated pixels to that of uncoated pixels, and its 95% confidence interval was obtained by bootstrap resampling of pixels.

### MALDI mass spectrometry imaging

Matrix-coated tissue sections were analyzed on a timsTOF fleX mass spectrometer (Bruker Daltonics, Bremen, Germany) equipped with a dual ESI/MALDI source and a 10 kHz smartbeam 3D Nd: YAG laser (λ = 355 nm) at the Johns Hopkins AIMS Core facility. Height adjustment and laser focus tuning were performed before each acquisition. Mass calibration was conducted using electrospray ionization (ESI) of Agilent ESI-L Tune Mix, followed by calibration with red phosphorus to achieve a mass error of <1 ppm. MALDI imaging parameters were optimized in qTOF mode to enhance signal intensity by adjusting ion optics, laser intensity, and laser focus. In the MALDI-1 mode, a primary laser directly irradiated the sample to ionize molecules. Key acquisition settings were as follows: MALDI plate offset of 50 V, deflection 1 delta of 70 V, funnel 1 RF of 400 Vpp, funnel 2 RF of 400 Vpp, multipole RF of 400 Vpp, collision cell energy of 10 eV, collision RF of 750 Vpp, quadrupole ion energy of 5 eV, low mass cutoff of *m/z* 20, pre-TOF transfer time of 70 µs, prepulse storage time of 10 µs, N_2_ source pressure of ∼2 mbar.

Imaging was performed in both positive- and negative-ion modes, covering a mass-to-charge (*m/z*) range of 20–1,000 Da. Two raster configurations were used: (1) a 5 μm raster width with 25 laser shots per pixel, and (2) a 10 μm raster width with 600 laser shots per pixel. For both configurations, the laser was operated at a frequency of 1 kHz with the laser power set to 60-90%. On-tissue tandem mass spectrometry (MS/MS) was performed for the ions at *m/z* 630.62 Da in positive-ion mode (**Supplementary Fig. S9**). Tandem MS experiments were performed with nitrogen as the collision gas, using a sum of 5 spectra with 600 shots per pixel across the *m/z* range of 20–1,000 Da. The laser power was set to 90% with an isolation window of ±1 Da and a collision energy of 40 eV. MALDI imaging data were imported into SCiLS Lab (version (v) 2027a, Bruker Daltonics) for visualization and analysis, focusing on average spectra within the MS imaging region. Average intensity was determined by the spectra from the ROI normalized to total ion count (TIC). Tandem MS spectra were processed and analyzed using dataAnalysis (v 5.3.236, Bruker Daltonics).

### iprm-PASEF acquisition, data processing, and data analysis

MALDI trapped ion mobility spectrometry (TIMS) MS1 imaging was performed on the timsTOF fleX mass spectrometer (Bruker Daltonics, Bremen, Germany) operating in positive- or negative-ion reflector mode, with pixel sizes of 100 µm × 100 µm, acquiring 300 laser shots per pixel at a frequency of 10 kHz. The resulting imaging data were processed in the SCiLS Lab software (Bruker Daltonics) using TIC normalization, with bin widths of 4 ppm for *m/z* and 0.01 V·s/cm² for 1/K_0_. Spectral features were identified using T-ReX3 (2 × 2 spatial smoothing, 50% minimum pixel coverage, 1% relative intensity threshold), supplemented by manual peak selection as needed. Targeted MS2 acquisition was performed using the iprm-PASEF module in timsControl 7.0 (Bruker Daltonics). iprm-PASEF imaging measurements were acquired in prm-PASEF positive-ion reflector mode using identical spatial resolution and laser parameters as the MS1 acquisition, with an *m/z* range of 20–1,000, a 1/K_0_ range of 0.40–1.72 V·s/cm², and a ramp time of 300 ms. Instrument tuning parameters were optimized as follows: collision RF of 750 Vpp, ion energy of 5 eV, transfer time of 75 µs, pre-pulse storage of 5 µs. The iprm-PASEF imaging method targeted three negative or two positive precursor ions simultaneously within predefined m/z and 1/K_0_ ranges at a collision energy of 40 eV. MS2 spectra were averaged across all pixels in DataAnalysis, and fragment ion images were visualized in SCiLS Lab.

### Lipid identification

Lipid annotation targeted 26 lipid classes previously reported in primate brain tissue^45^, including phosphatidylethanolamine (PE), phosphatidylcholine (PC), their ether-linked species (PE-O, PC-O), phosphatidylglycerol (PG), phosphatidylserine (PS), phosphatidic acid (PA), phosphatidylinositol (PI), lysophosphatidylcholine (LPC), lysophosphatidylethanolamine (LPE), their ether-linked species (LPC-O, LPE-O), lysophosphatidylserine (LPS), lysophosphatidic acid (LPA), ceramide (Cer), acylceramide (ACer), ceramide phosphate (CerP), sphingomyelin (SM), hexosylceramide (HexCer), dihexosylceramide (Hex2Cer), sulfatide (SHexCer), galactosylceramide (GalCer), diacylglycerol (DG), dimyristoylphosphatidylcholine (DMPC), sulfates, and cholesterol.

Lipid class definitions followed the LIPID MAPS comprehensive classification system^45^, with three modifications: free fatty acid subclasses were consolidated into a single class, the Cer class was expanded to include dihydroceramides and phytoceramides, and the HexCer class encompassed both galactosylceramides and glucosylceramides.

The annotation workflow consisted of the following steps. First, a custom reference database was constructed containing theoretical *m/z* values for the 26 lipid classes, accounting for variations in acyl chain length and degree of unsaturation. Mass spectrometry peaks from our dataset were then matched against this database using a mass tolerance of 10 ppm, with fixed adduct assignments for each lipid class.

Additionally, publicly available MALDI-MSI datasets were retrieved from the METASPACE repository^46^ and searched for corresponding lipid classes using a false discovery rate (FDR) threshold of <20%.

### Embryo spatial lipidomics data acquisition and analysis

Lipidomic intensity data were acquired from triplicates (n = 3) of *SMPD3* gain-of-function (GOF) and control samples across three embryonic tissue lineages: neural crest, neural tube, and non-neural ectoderm. Lipid features were identified by *m/z* and lipid class assignment. Features with zero intensity across all biological observations were excluded, no imputation was applied, and a pseudocount (+1) was added only for logarithmic transformations.

Sphingolipid profiles were compared between *SMPD3* GOF and control conditions across the three embryonic tissue lineages. TIC-normalized intensities from three replicates per condition were processed for heatmap display using custom R scripts (v 4.5) with the pheatmap package. Only SM and Cer species with non-zero intensities across all samples were retained. Replicate intensities within each condition-tissue combination were averaged, log2-transformed (pseudocount +1), and Z-score–normalized across samples for each lipid species independently. Hierarchical clustering of lipid species displayed in rows used Euclidean distance with complete linkage. Columns were organized by tissue lineage, with control and *SMPD3* GOF conditions paired within each group. Heatmaps used a diverging red-white-blue palette (RdBu, RColorBrewer).

Volcano plots were generated for each tissue lineage separately, with ΔE (log2 fold change GOF/control) on the x-axis and −log_10_( *P*) on the y-axis. Vertical lines demarcate the fold-change threshold at |ΔE| = 1. Points exceeding this threshold were colored by lipid class, and remaining points were rendered in grey. Point size scaled with mean intensity [(µ_GOF_ + µ_Ctrl_)/2]. Top-ranked species satisfying both |ΔE| ≥ 1 and *P* < 0.05 were labeled and ranked by a combined score [|ΔE| × −log_10_(*P*)], with non-overlapping placement (ggrepel).

### Co-crystal preparation

For CHCA–rhodamine B co-crystals, CHCA and rhodamine B were combined at a 1:1.25 molar ratio and dissolved in 50% toluene. The mixture was left to evaporate slowly at room temperature. Deep red-violet needle-shaped crystals formed after approximately one week. For nH–A488 co-crystals, equimolar amounts of nH and A488 were dissolved in 50% dichloromethane and allowed to evaporate at room temperature. Amber-orange rod-shaped crystals formed after approximately three weeks.

### Single Crystal X-ray crystallography

All reflection intensities were measured at 110.00(10) K on a Rigaku XtaLAB Synergy R diffractometer equipped with a rotating-anode X-ray source (Cu Kα radiation, λ = 1.54178 Å) and a HyPix-6000HE detector (Rigaku, Tokyo, Japan). Temperature was controlled by a Cryostream 1000 system (Oxford Cryosystems). Data collection, cell refinement, and data reduction were performed in CrysAlisPro (v 1.171.42.49, Rigaku, 2022).

### Structure determination and refinement

Structures were solved using SHELXT-2018/2 and refined on *F*^2^ using SHELXL-2019/3^47^. An analytical numeric absorption correction based on a multifaceted crystal model was applied in CrysAlisPro. Crystallographic data have been deposited with the Cambridge Crystallographic Data Centre (CCDC 2293689^4^ for CHCA-Rhodamine B and 2588717 for nH-A488). Additional refinement details are provided in the Supplementary Information.

### Point spread function (PSF) measurement

Lateral point spread function (PSF) measurements were performed using sub-resolution fluorescent beads imaged on a Nikon Ti2 microscope equipped with a 100× objective. Raw TIFF images were exported and analyzed using a custom Python-based analysis pipeline (NumPy, SciPy, Pandas). Individual beads were automatically detected and cropped into local ROIs. Each bead intensity distribution was fit to a two-dimensional Gaussian model to extract the standard deviations of σ_x_ and σ_y_ along the lateral axes.

The full width at half maximum (FWHM) was calculated from the fitted Gaussian parameters using FWHM = 2√(2 ln 2)·σ ≈ 2.355σ for both x and y directions. Pixel dimensions were converted to nanometers using the calibrated pixel size of the imaging system. Beads with poor fits or aberrant dimensions were excluded using a 1.5× interquartile range (IQR) outlier criterion applied independently to FWHM_x_ and FWHM_y_ distributions. Reported values represent the mean ± standard deviation of the cleaned dataset. For summary analysis, a combined FWHM metric was defined as (FWHM_x_ + FWHM_y_)/2.

Statistical comparisons between matrix-deposition groups and bead controls were performed using the two-sided Mann–Whitney U test (SciPy) for each exposure condition of 20 ms and 50 ms, with Holm–Bonferroni correction for multiple comparisons. Significance thresholds: * *P* < 0.05, \*\**P* < 0.01, \*\*\**P* < 0.001.

### Statistical analysis and software

Statistical analyses were performed in R (v 4.5) using ggplot2 (v 4.0.3), dplyr (v 1.1.0), readxl (v 1.4.2), uwot (v 0.1.14), geometry (v 0.4.6.1) and plotly (v 4.10.1), and in JASP (v 0.96.0). Pixel-level comparisons (Fig. 2c,h, 3c and 4c,e and the corresponding Supplementary Figures) used two-sided nonparametric tests. Two conditions were compared with the Mann–Whitney U test for independent samples or the Wilcoxon signed-rank test for matched pixels. Three groups were compared with the Kruskal–Wallis test, followed by Dunn tests with Holm adjustment for all pairwise comparisons. Effect sizes were Cliff’s δ (Mann–Whitney U and Dunn comparisons) or the matched-pairs rank-biserial correlation (signed-rank test), each with a 95% bootstrap confidence interval (2,000 resamples), and shifts between groups were summarized with the Hodges–Lehmann estimator. The Mann–Whitney U and Kruskal–Wallis tests compare distributions, and they compare medians only when the distributions have similar shapes, so differences are described as differences between distributions unless the shapes were similar. Box plots show the median, the interquartile range and whiskers extending to the most extreme values within 1.5 times the interquartile range. P values are exact to two significant figures, values below 0.0001 are displayed as *P* < 0.0001 in the figures, and exact values are given in Source Data. Pixels from one section are not independent biological replicates, so pixel-level *P* values describe the imaged section and are reported together with effect sizes. No statistical methods were used to predetermine sample sizes. Experiments were not randomized, and investigators were not blinded during experiments or outcome assessments.

## Reporting summary

Further information on research design is available in Reporting Summary linked to this article.

## Data availability

The FluoMALDI-MSI data generated in this study are available from the corresponding author upon reasonable request.

## Author contributions

T.-H.H. and K.G. conceived and designed the study. T.-H.H., M.L.P., W.A., J.C.T., D.R.B., C.M.T., and S.T. prepared samples. Fluorescence staining was performed by T.-H.H., M.L.P., A.P.P., and W.A., and fluorescence microscopy was conducted by T.-H.H., M.L.P., W.A., and S.T. X-ray crystallography was performed by M.A.S. and T.-H.H. T.-H.H. performed MALDI-MSI acquisition and data processing. Image analysis using SCiLS Lab, FIJI, Python, and R was carried out by T.-H.H., M.L.P., and S.T. T.-H.H. designed the figures with input from M.L.P., S.T., I.B., V.M., and K.G. T.-H.H. wrote the manuscript under the supervision of K.G. All authors reviewed and approved the final manuscript.

## Acknowledgements

We would like to acknowledge the instruments and expertise of the Johns Hopkins Applied Imaging Mass Spectrometry (AIMS) Core Facility at the Johns Hopkins University School of Medicine, which were critical in advancing this work. The authors further acknowledge funding from the National Institutes of Health (NIH) of the United States of America, grants R01 CA264901 (to K.G.), R01 EB037743 (to K.G.), P30 CA006973 Shared Resource for Mass Spectrometry Molecular Imaging and Multi-Omics (to K.G.), S10 OD030500 (to K.G.), S10 OD030352 (to M.A.S.), R35 GM155257 (to M.L.P.), R35 GM149272 (to I.B.), K99 GM166955 (to T.-H.H.). We thank The Richman Family Precision Medicine Center of Excellence in Alzheimer’s Disease at Johns Hopkins University for funding support.

## References

1. Tuck, M., Grélard, F., Blanc, L. & Desbenoit, N. MALDI-MSI towards multimodal imaging: challenges and perspectives. Front. Chem. 10, 904688 (2022).

2. Yang, E. et al. RaMALDI: enabling simultaneous Raman and MALDI imaging of the same tissue section. Biosens. Bioelectron. 239, 115597 (2023).

3. Hahm, T.-H. et al. MALDI matrix sublimation versus spray coating in the FluoMALDI imaging pipeline. Anal. Chim. Acta 1376, 344610 (2025).

4. Yang, E. et al. FluoMALDI microscopy: matrix co-crystallization simultaneously enhances fluorescence and MALDI imaging. Adv. Sci. 10, 2304343 (2023).

5. Bourceau, P. et al. Visualization of metabolites and microbes at high spatial resolution using MALDI mass spectrometry imaging and in situ fluorescence labeling. Nat. Protoc. 18, 3050–3079 (2023).

6. Liang, Z., Guo, Y., Sharma, A., McCurdy, C. R. & Prentice, B. M. A multi-modal image fusion workflow incorporating MALDI imaging mass spectrometry and microscopy for the study of small pharmaceutical compounds. Anal. Chem. 96, 11869–11880 (2024).

7. Hahm, T. -H. et al. Quantitative MALDI imaging of aspirin metabolites in mouse models of triple-negative breast cancer. Theranostics 15, 9359–9374 (2025).

8. Hickey, J. W. et al. Spatial mapping of protein composition and tissue organization: a primer for multiplexed antibody-based imaging. Nat. Methods 19, 284–295 (2022).

9. Pellanda, D., Dressler, E. & Frei, M. S. Chemigenetic fluorescent biosensors in biological imaging - New trends and advances. Curr. Opin. Chem. Biol. 93, 102707 (2026).

10. Lin, J., Yang, K. & New, E. J. Strategies for organelle targeting of fluorescent probes. Org. Biomol. Chem. 19, 9339–9357 (2021).

11. Kiernan, J. A. Anatomy of the temporal lobe. Epilepsy Res. Treat. 2012, 176157 (2012).

12. Liu, Y., Su, Y. & Wang, X. Phosphatidic acid-mediated signaling. Adv. Exp. Med. Biol. 991, 159–176 (2013).

13. Cazzolli, R., Shemon, A. N., Fang, M. Q. & Hughes, W. E. Phospholipid signalling through phospholipase D and phosphatidic acid. IUBMB Life 58, 457–461 (2006).

14. Martin, T. F. J. Phosphoinositide lipids as signaling molecules: common themes for signal transduction, cytoskeletal regulation, and membrane trafficking. Annu. Rev. Cell Dev. Biol. 14, 231–264 (1998).

15. Palhol, J. S. C. et al. Direct association with the vascular basement membrane is a frequent feature of myelinating oligodendrocytes in the neocortex. Fluids Barriers CNS 20, 24 (2023).

16. Yokoo, H. et al. Anti-human Olig2 antibody as a useful immunohistochemical marker of normal oligodendrocytes and gliomas. Am. J. Pathol. 164, 1717–1725 (2004).

17. Vega-Lopez, G. A., Cerrizuela, S., Tribulo, C. & Aybar, M. J. Neurocristopathies: New insights 150 years after the neural crest discovery. Dev. Biol. 444, S110–S143 (2018).

18. Mancini, A. E., Jenne, M. P. & Piacentino, M. L. Membrane lipid metabolism as a regulatory frontier in neural crest biology: Roles for sphingolipids, cholesterol, and lipid rafts. Dev. Dyn. 1–26 (2026).

19. Piacentino, M. L., Fasse, A. J., Camacho-Avila, A., Grabylnikov, I. & Bronner, M. E. SMPD3 expression is spatially regulated in the developing embryo by SOXE factors. Dev. Biol. 506, 31–41 (2024).

20. Piacentino, M. L., Hutchins, E. J., Andrews, C. J. & Bronner, M. E. Temporal changes in plasma membrane lipid content induce endocytosis to regulate developmental epithelial-to-mesenchymal transition. Proc. Natl. Acad. Sci. U.S.A. 119, e2212879119 (2022).

21. Los, G. V. et al. HaloTag: a novel protein labeling technology for cell imaging and protein analysis. ACS Chem. Biol. 3, 373–382 (2008).

22. Williams, R. M. & Sauka-Spengler, T. Ex ovo electroporation of early chicken embryos. STAR Protoc. 2, 100424 (2021).

23. Shaner, N. et al. Improved monomeric red, orange and yellow fluorescent proteins derived from *Discosoma* sp. red fluorescent protein. Nat. Biotechnol. 22, 1567–1572 (2004).

24. Shamseddine, A. A., Airola, M. V. & Hannun, Y. A. Roles and regulation of neutral sphingomyelinase-2 in cellular and pathological processes. Adv. Biol. Regul. 57, 24–41 (2015).

25. Airola, M. V., et al. Structure of human nSMase2 reveals an interdomain allosteric activation mechanism for ceramide generation. Proc. Natl. Acad. Sci. U.S.A. 114, E5549–E5558 (2017).

26. Hovland, A. S. et al. Pluripotency factors are repurposed to shape the epigenomic landscape of neural crest cells. Dev. Cell 57, 2257–2272.e5 (2022).

27. Komai, M. et al. Nuclear SphK2/S1P signaling is a key regulator of ApoE production and Aβ uptake in astrocytes. J. Lipid Res. 65, 100510 (2024).

28. Guttenplan, K. A. et al. Neurotoxic reactive astrocytes induce cell death via saturated lipids. Nature 599, 102–107 (2021).

29. Chen, Z.-P. et al. Lipid-accumulated reactive astrocytes promote disease progression in epilepsy. Nat. Neurosci. 26, 542–554 (2023).

30. Joshi, G., Bekier, M. E. 2nd & Wang, Y. Golgi fragmentation in Alzheimer’s disease. Front. Neurosci. 9, 340 (2015).

31. Panchuk-Voloshina, N. et al. Alexa Fluor dyes, a series of new, fluorescent dyes that yield exceptionally bright, photostable conjugates. J. Histochem. Cytochem. 47, 1179–1188 (1999).

32. Eggeling, C. Fluorophore conjugates for single molecule work. In Single Molecule Spectroscopy in Chemistry, Physics and Biology (eds. Rigler, R., Orrit, M. & Röhlich, T.) 35–51 (Springer, Berlin, Heidelberg, 2011).

33. Beija, M., Afonso, C. A. M. & Martinho, J. M. G. Synthesis and applications of Rhodamine derivatives as fluorescent probes. Chem. Soc. Rev. 38, 2410–2433 (2009).

34. Brann, D. W., et al. Brain-derived estrogen and neural function. Neurosci. Biobehav. Rev. 132, 793–817 (2022).

35. Garcia-Segura, L. M. & Melcangi, R. C. Steroids and glial cell function. Glia 54, 485–498 (2006).

36. Vitku, J. et al. Steroid sulfation in neurodegenerative diseases. Front. Mol. Biosci. 9, 839887 (2022).

37. Zwain, I. H. & Yen, S. S. C. Neurosteroidogenesis in astrocytes, oligodendrocytes, and neurons of cerebral cortex of rat brain. Endocrinology 140, 3843–3852 (1999).

38. Kasha, M., Rawls, H. R. & El-Bayoumi, M. A. The exciton model in molecular spectroscopy. Pure Appl. Chem. 11, 371–392 (1965).

39. Potthoff, A., et al. Spatial biology using single-cell mass spectrometry imaging and integrated microscopy. Nat. Commun. 16, 9129 (2025).

40. Young, R. S. E. et al. Subcellular mass spectrometry imaging of lipids and nucleotides using transmission geometry ambient laser desorption and plasma ionisation. Nat. Commun. 16, 9130 (2025).

41. Chang, K. et al. Loss of TDP-43 splicing repression occurs early in the aging population and is associated with Alzheimer’s disease neuropathologic changes and cognitive decline. Acta Neuropathol. 147, 4 (2023).

42. Megason, S. G. & McMahon, A. P. A mitogen gradient of dorsal midline Wnts organizes growth in the CNS. Development 129, 2087–2098 (2002).

43. Tcw, J. et al. An efficient platform for astrocyte differentiation from human induced pluripotent stem cells. Stem Cell Rep. 9, 600–614 (2017).

44. Schindelin, J. et al. Fiji: an open-source platform for biological-image analysis. Nat. Methods 9, 676–682 (2012).

45. Fahy, E. et al. Update of the LIPID MAPS comprehensive classification system for lipids. J. Lipid Res. 50, S9–S14 (2009).

46. Palmer, A. et al. FDR-controlled metabolite annotation for high-resolution imaging mass spectrometry. Nat. Methods 14, 57–60 (2017).

47. Sheldrick, G. M. Crystal structure refinement with SHELXL. Acta Crystallogr. C Struct. Chem. 71, 3–8 (2015).

