## Supplementary Information for "FluoMALDI Imaging: An Integrated Multimodality Platform for Biologically Informed Spatial-Omics"

**Supplementary Information:** Figures S1 to S21 and Tables S1 and S2.

Tae-Hun Hahm[a,b], Michael L. Piacentino[c], Waqar Ahmed[d], Maxime A. Siegler[e], Dalton R. Brown[a,b], Caitlin M. Tressler[a,b], Juan C. Troncoso[f,g], Arvind P. Pathak[b], Swati Tanwar[h], Ishan Barman[b,h], Vasiliki Mahairaki[d], Kristine Glunde[a,b,f,i,*]

[a]Applied Imaging Mass Spectrometry Core, Johns Hopkins University School of Medicine, Baltimore, MD, 21287 USA; [b]Russell H. Morgan Department of Radiology and Radiological Science, Johns Hopkins University School of Medicine, Baltimore, MD, 21287 USA; [c]Department of Cell Biology, Johns Hopkins University School of Medicine, Baltimore, MD, 21205 USA; [d]Department of Genetic Medicine, Johns Hopkins University School of Medicine, Baltimore, MD, 21205 USA; [e]Department of Chemistry, Johns Hopkins University, Baltimore, MD, 21218 USA; [f]Department of Pathology, Division of Neuropathology, Johns Hopkins University School of Medicine, Baltimore, MD, 21205 USA; [g]Department of Neurology, Johns Hopkins University School of Medicine, Baltimore, MD, 21231 USA; [h]Department of Mechanical Engineering, Johns Hopkins University, Baltimore, MD, 21218, USA; [i]Sidney Kimmel Comprehensive Cancer Center, Johns Hopkins University School of Medicine, Baltimore, MD, 21231 USA.

*Corresponding author:
Kristine Glunde, Ph.D.
Professor of Radiology, Oncology and Pathology
Director, Applied Imaging Mass Spectrometry (eAIMS)
The Johns Hopkins University School of Medicine
Russell H. Morgan Department of Radiology and Radiological Science
Division of Cancer Imaging Research
720 Rutland Avenue
Traylor Building, Room 203
Baltimore, MD 21205, U.S.A.


### Supplementary Figures

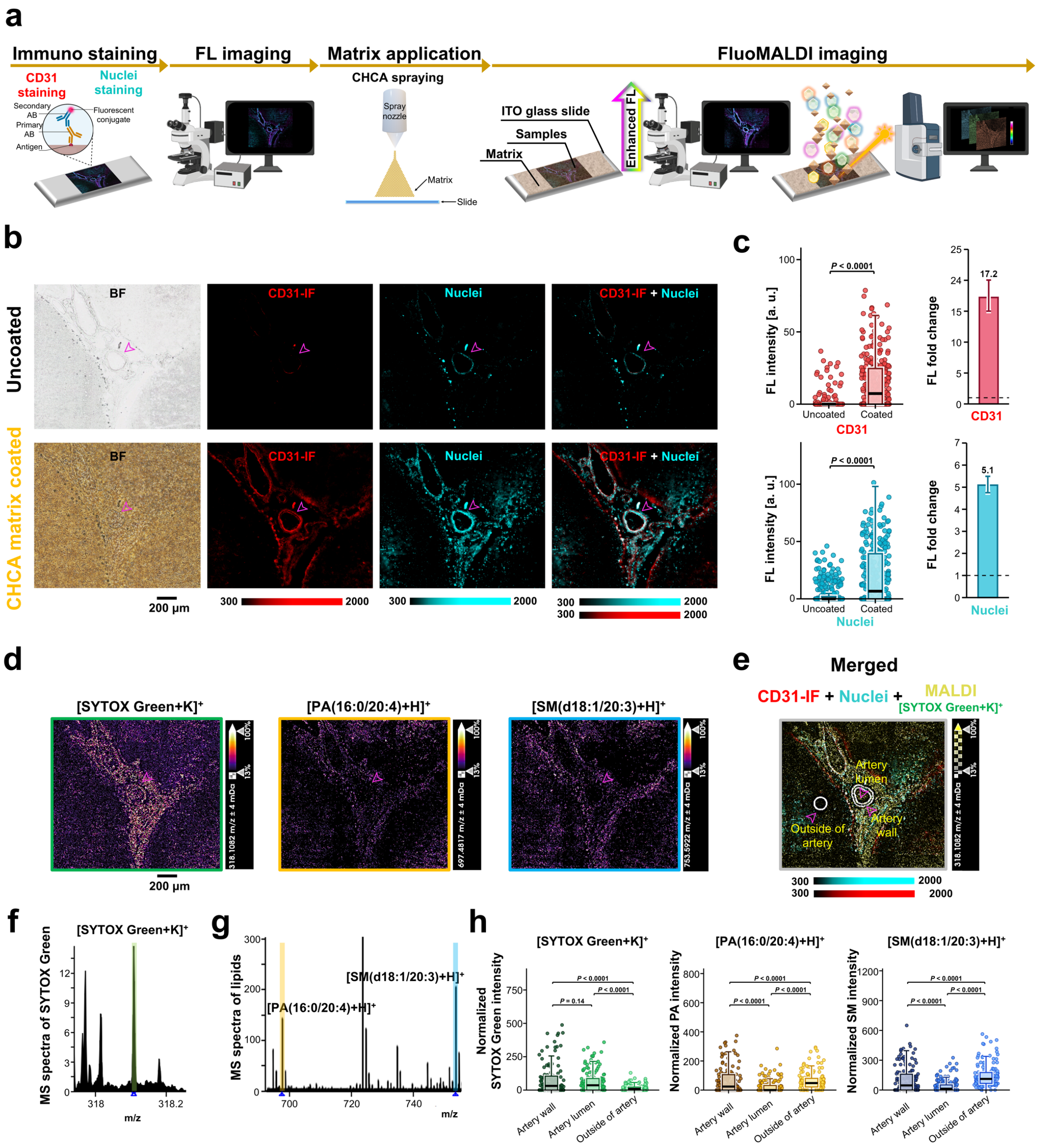

**Supplementary Fig. S1: Immuno-FluoMALDI imaging of human brain tissue sections using CHCA matrix.** (**a**) Workflow for Immuno-FluoMALDI analysis. A human brain tissue section was mounted on a slide and subjected to immunofluorescence (IF) labeling. Optical microscopy was performed to compare the fluorescence of uncoated and coated tissue sections after CHCA matrix was applied by spray deposition. (**b**) Brightfield (BF) and fluorescence images of a human hippocampal section before (uncoated) and after coating with 0.9 µg/mm² CHCA. BF shows tissue morphology without a chromogen, and after coating also the matrix layer. CD31 was detected with an Alexa Fluor Plus 647-conjugated secondary antibody (CD31-IF, red) and nuclei with SYTOX Green (cyan), and both are visible only in the fluorescence channels. Magenta arrowheads mark the hippocampal artery analyzed in this study (a sulcal artery in the hippocampal fissure adjacent to CA1). Other CD31-positive vessels in the field of view were not analyzed. Scale bar: 200 µm. (**c**) Fluorescence intensity (a.u.) of CD31-IF (top) and nuclei (bottom) in the artery region before and after CHCA coating, with fold change (right, ratio of medians, 95% bootstrap confidence interval, the dashed line marks 1). Each point is one pixel (one section, one donor). Boxes show the median and interquartile range, and whiskers show 1.5 times the interquartile range. Statistical comparisons used two-sided Mann-Whitney U tests. (**d**) MALDI MS images in positive ion mode were obtained from the region of interest (ROI) containing hippocampal arteries (magenta arrows) in panel b. The MALDI images display SYTOX Green at $m/z^+$ 318.11, PA(16:0/20:4) at $m/z^+$ 697.48, and SM(d18:1/20:3) at $m/z^+$ 753.59. Scale bars: 200 µm. (**e**) Co-registered enhanced fluorescence microscopy data (CD31-IF in red, nuclei in cyan) and MALDI imaging data of SytoxGreen (intensity scale converted to yellow color in overlaid image). Arrows point out ROIs for quantification. (**f** and **g**) Average MALDI MSI spectra from the MALDI-imaged ROI with color-coded labels for *m/z* signals shown as MALDI images in d. (**h**) TIC-normalized signal intensity of SYTOX Green ($[M+K]^+$), PA(16:0/20:4) and SM(d18:1/20:3) in artery wall, lumen and outside of artery (regions in e). Each point is one pixel (one artery, one section, one donor). Boxes show the median and interquartile range, and whiskers show 1.5 times the interquartile range. Kruskal–Wallis tests were followed by two-sided Dunn tests with Holm adjustment, and adjusted P values are shown. The hotspot intensity cutoff for pixel selection was defined by the hotspot quantile setting in the SCiLS Lab Settings dialog, configured at 99.0%. Exact P values and effect sizes are in Source Data.

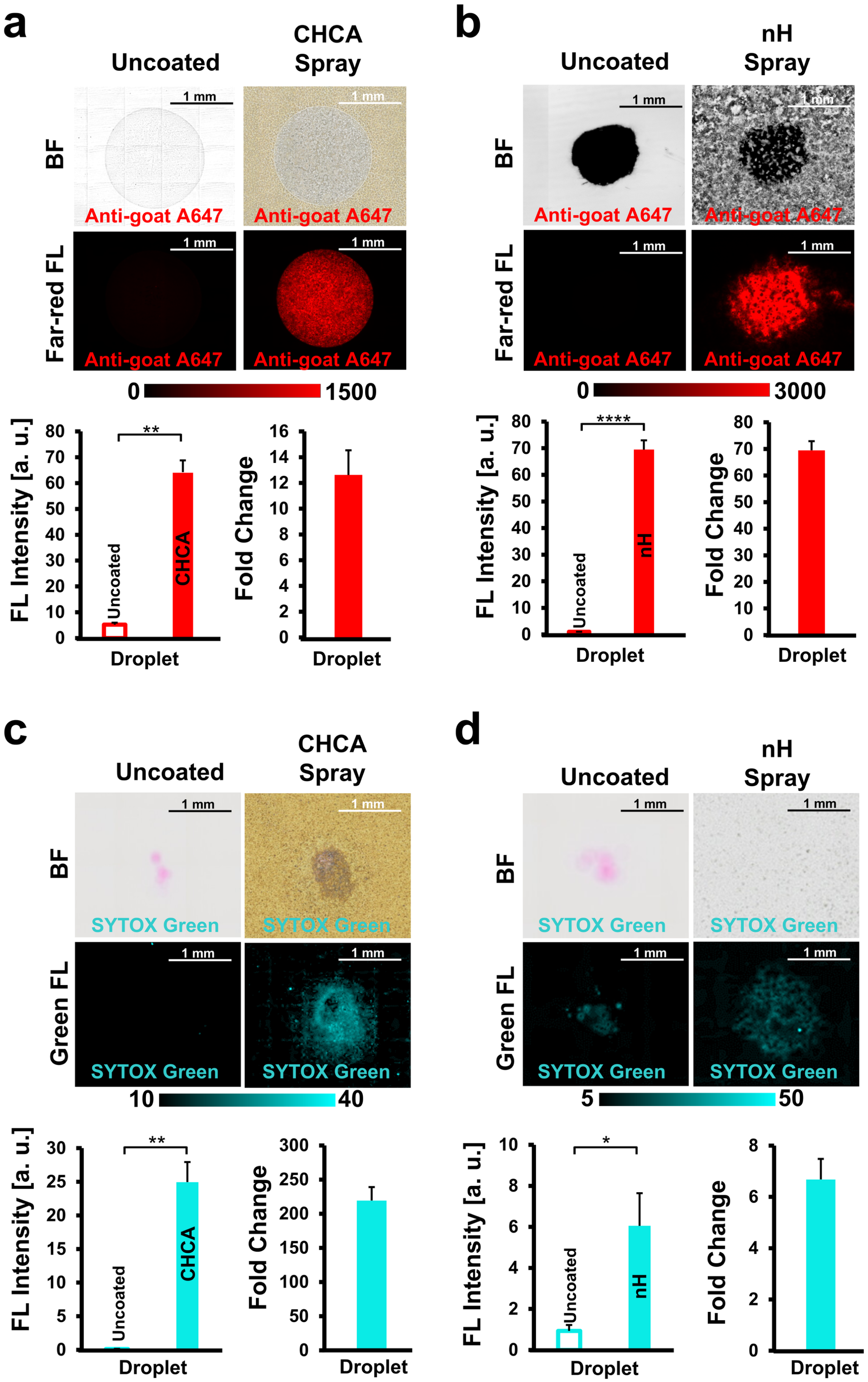
a
Uncoated
CHCA Spray
BF
Far-red FL
1 mm
Anti-goat A647
0
1500
FL Intensity [a. u.]
**
Uncoated
CHCA
Droplet
Fold Change
Droplet
b
Uncoated
nH Spray
BF
Far-red FL
1 mm
Anti-goat A647
0
3000
FL Intensity [a. u.]
****
Uncoated
nH
Droplet
Fold Change
Droplet
c
Uncoated
CHCA Spray
BF
Green FL
1 mm
SYTOX Green
10
40
FL Intensity [a. u.]
**
Uncoated
CHCA
Droplet
Fold Change
Droplet
d
Uncoated
nH Spray
BF
Green FL
1 mm
SYTOX Green
5
50
FL Intensity [a. u.]
*
Uncoated
nH
Droplet
Fold Change
Droplet

**<u>Supplementary Fig. S2:</u> Matrix coating effects on fluorescent dried droplet detection. Top panels** (**a-d**): Brightfield (BF, top) and fluorescence (FL, bottom) images of uncoated (left) and CHCA (**a**, **c**) or nH (**b**, **d**) spray-coated (right) pure dried A647-conjugated anti-goat antibody (20 µg/mL; **a**, **b**) or SYTOX Green (1 µM; **c**, **d**) droplets. Scale bars: 1 mm. Color scales indicate fluorescence intensity. **Bottom panels** (**a-d**): Quantification of fluorescence intensity (left graphs) and fold change (right graphs) of dried droplets comparing uncoated versus matrix-coated conditions. Error bars represent standard deviation (n = 3 droplets).

**a**

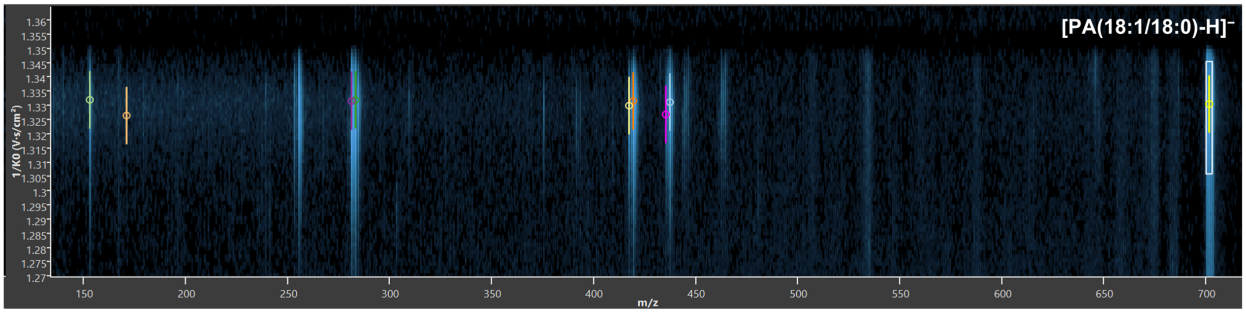

| *m/z* | *m/z* width [± ppm] | $1/K_0$ [V·s/cm²] | $1/K_0$ width [±] | Intensity [Regions] | Chemical formula | Adduct ion | Type | Isolation window |
|---|---|---|---|---|---|---|---|---|
| 152.9957 | 4.8 | 1.332 | 0.01 | 866.5955 | $C_3H_5O_5P^{\bullet}$ | $[M]^-$ | Fragment | 701.5159 *m/z*, $1/K_0$ 1.3257 |
| 171.0065 | 4.8 | 1.3265 | 0.01 | 19.10557 | $C_3H_8O_6P^{\bullet}$ | $[M]^-$ | Fragment | 701.5159 *m/z*, $1/K_0$ 1.3257 |
| 281.2488 | 4.8 | 1.3316 | 0.01 | 3966.749 | $C_{18}H_{33}O_2^{\bullet}$ | $[M]^-$ | Fragment | 701.5159 *m/z*, $1/K_0$ 1.3257 |
| 283.2643 | 4.8 | 1.332 | 0.01 | 4268.236 | $C_{18}H_{35}O_2^{2\bullet}$ | $[M]^-$ | Fragment | 701.5159 *m/z*, $1/K_0$ 1.3257 |
| 417.2415 | 4.8 | 1.33 | 0.01 | 838.3915 | $C_{21}H_{39}O_6P^{\bullet}$ | $[M-H]^-$ | Fragment | 701.5159 *m/z*, $1/K_0$ 1.3257 |
| 419.257 | 4.8 | 1.3316 | 0.01 | 2650.547 | $C_{21}H_{41}O_6P$ | $[M-H]^-$ | Fragment | 701.5159 *m/z*, $1/K_0$ 1.3257 |
| 435.2523 | 4.8 | 1.3269 | 0.01 | 255.256 | $C_{21}H_{40}O_7P^{\bullet}$ | $[M]^-$ | Fragment | 701.5159 *m/z*, $1/K_0$ 1.3257 |
| 437.2677 | 4.8 | 1.3312 | 0.01 | 1024.014 | $C_{21}H_{42}O_7P^{\bullet}$ | $[M]^-$ | Fragment | 701.5159 *m/z*, $1/K_0$ 1.3257 |
| 701.5159 | 4.8 | 1.3305 | 0.01 | 1843.338 | $C_{39}H_{75}O_8P$ | $[M-H]^-$ | Precursor | 701.5159 *m/z*, $1/K_0$ 1.3257 |

**b**

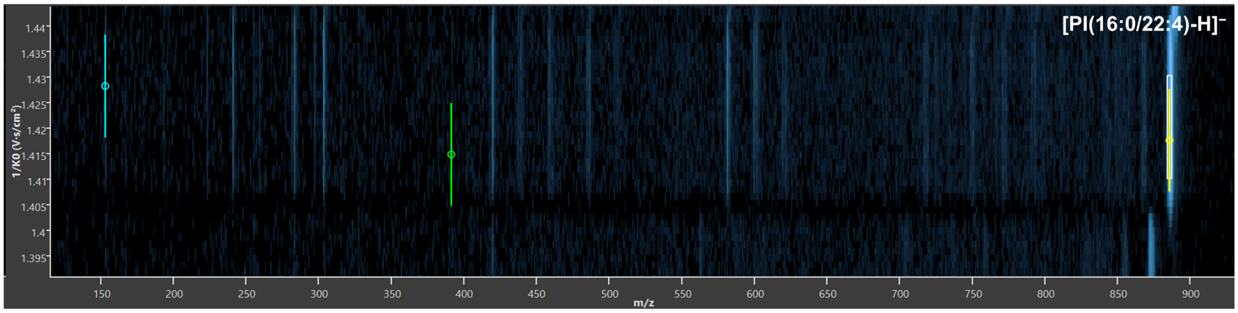

| *m/z* | *m/z* width [± ppm] | $1/K_0$ [V·s/cm²] | $1/K_0$ width [±] | Intensity [Regions] | Chemical formula | Adduct ion | Type | Isolation window |
|---|---|---|---|---|---|---|---|---|
| 152.9974 | 4.8 | 1.4283 | 0.01 | 3.687683 | $C_3H_6O_5P^{\bullet}$ | $[M]^-$ | Fragment | 885.5537 *m/z*, $1/K_0$ 1.4203 |
| 391.2253 | 4.8 | 1.4149 | 0.01 | 1.005492 | $C_{19}H_{37}O_6P^{\bullet}$ | $[M-H]^-$ | Fragment | 885.5537 *m/z*, $1/K_0$ 1.4203 |
| 885.5537 | 4.8 | 1.4176 | 0.01 | 2800.504 | $C_{47}H_{83}O_{13}P$ | $[M-H]^-$ | Precursor | 885.5537 *m/z*, $1/K_0$ 1.4203 |

**c**

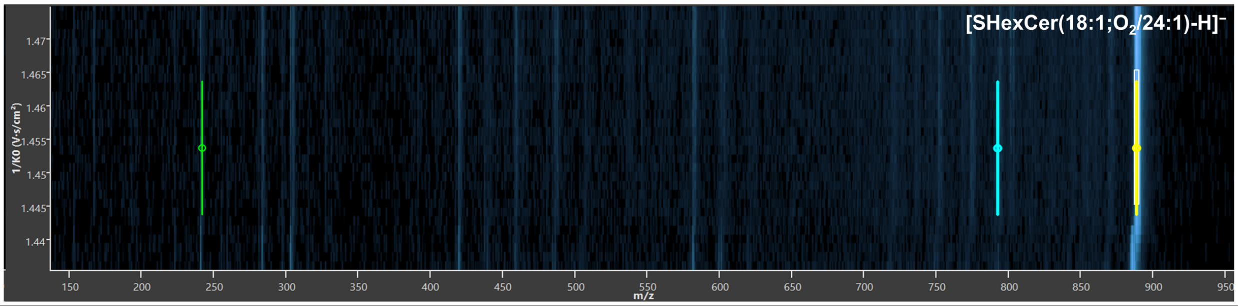

| *m/z* | *m/z* width [± ppm] | $1/K_0$ [V·s/cm²] | $1/K_0$ width [±] | Intensity [Regions] | Chemical formula | Adduct ion | Type | Isolation window |
|---|---|---|---|---|---|---|---|---|
| 242.0121 | 4.8 | 1.4537 | 0.01 | 5.02304137 | $C_6H_{10}O_8S^{\bullet}$ | $[M]^-$ | Fragment | 888.6277 *m/z*, $1/K_0$ 1.4553 |
| 792.5284 | 4.8 | 1.4537 | 0.01 | 5.46711326 | $C_{48}H_{90}NO_7^{\bullet}$ | $[M]^-$ | Fragment | 888.6277 *m/z*, $1/K_0$ 1.4553 |
| 888.6277 | 4.8 | 1.4537 | 0.01 | 7032.025 | $C_{48}H_{91}NO_{11}S$ | $[M-H]^-$ | Precursor | 888.6277 *m/z*, $1/K_0$ 1.4553 |

**Supplementary Fig. S3: Multiplexed lipid precursor identification using negative ion mode iprm-PASEF profiling of dye- and IF-stained human hippocampal brain tissue section.** Three distinct precursor ions were simultaneously isolated within a single iprm-PASEF window in negative-ion mode MALDI and successfully resolved using ion mobility separation. (**a**) The precursor at *m/z*⁻ 701.52 ($1/K_0$ ~1.33 V·s/cm²) was identified as phosphatidic acid PA(18:1/18:0) based on characteristic fragment ions including *m/z*⁻ 437.26, 435.25, 419.25, 417.24, 283.26, 281.24, 171.00, and 152.99. (**b**) The precursor at *m/z*⁻ 885.55 ($1/K_0$ ~1.42 V·s/cm²) was identified as phosphatidylinositol PI(16:0/22:4) with diagnostic fragments at *m/z*⁻ 391.22 and 152.99. (**c**) The precursor at *m/z*⁻ 888.63 ($1/K_0$ ~1.45 V·s/cm²) was identified as sulfatide SHexCer(18:1;$O_2$/24:1) based on fragments at *m/z*⁻ 792.52 and 242.01. Heat maps display ion mobility ($1/K_0$) versus *m/z* with corresponding fragment ion tables showing mass accuracy, mobility values, intensities, chemical formulas, adduct types, and isolation window assignments. Colored markers indicate precursor (yellow and boxed) and fragment ions used for structural identification.

**a**

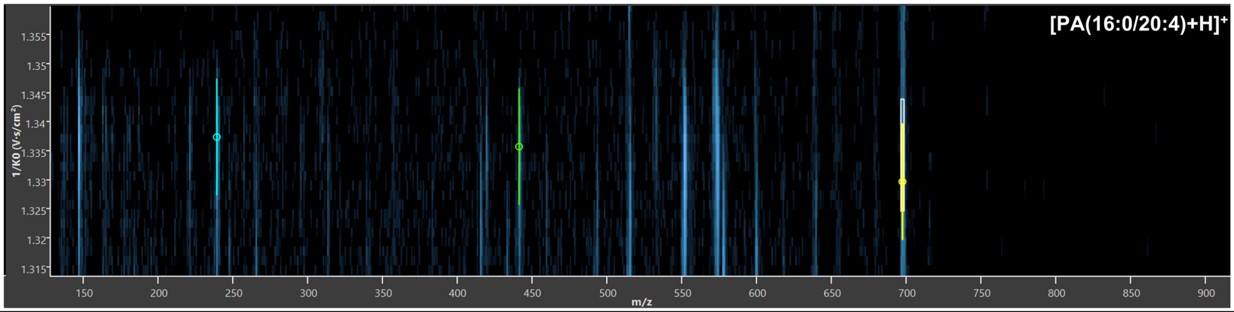

| *m/z* | *m/z* width [± ppm] | $1/K_0$ [V·s/cm²] | $1/K_0$ width [±] | Intensity [Regions] | Chemical formula | Adduct ion | Type | Isolation window |
|---|---|---|---|---|---|---|---|---|
| 239.2369 | 4.2 | 1.3374 | 0.01 | 1.001253 | $C_{16}H_{31}O^{\cdot}$ | $[M]^+$ | Fragment | 697.4817 m/z, $1/K_0$ 1.3343 |
| 441.2346 | 4.2 | 1.3357 | 0.01 | 1062.84 | $C_{23}H_{38}O_6P^{\cdot}$ | $[M]^+$ | Fragment | 697.4817 m/z, $1/K_0$ 1.3343 |
| 697.4817 | 4.2 | 1.3297 | 0.01 | 201.4082 | $C_{39}H_{69}O_8P$ | $[M+H]^+$ | Precursor | 697.4817 m/z, $1/K_0$ 1.3343 |

**b**

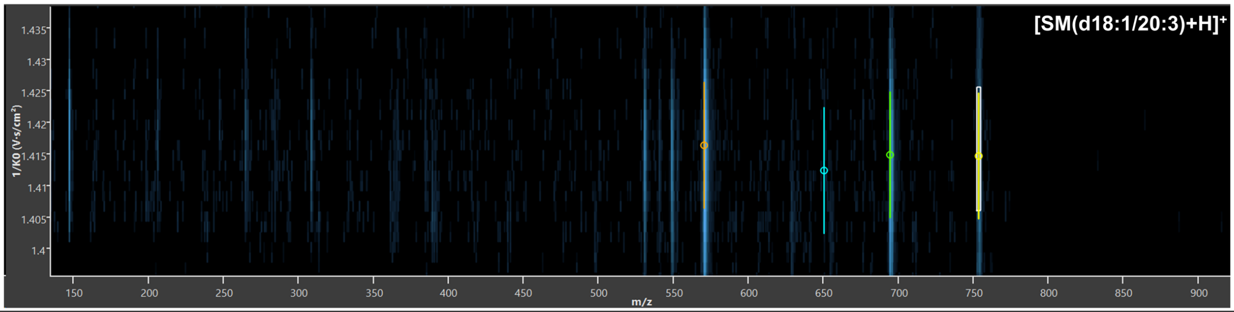

| *m/z* | *m/z* width [± ppm] | $1/K_0$ [V·s/cm²] | $1/K_0$ width [±] | Intensity [Regions] | Chemical formula | Adduct ion | Type | Isolation window |
|---|---|---|---|---|---|---|---|---|
| 570.5187 | 4.2 | 1.4164 | 0.01 | 34945.12 | $C_{38}H_{68}NO_2^{\cdot -}$ | $[M]^+$ | Fragment | 753.5922 m/z, $1/K_0$ 1.4158 |
| 650.4813 | 4.2 | 1.4124 | 0.01 | 12.31803 | $C_{38}H_{68}NO_5P^{-}$ | $[M+H]^+$ | Fragment | 753.5922 m/z, $1/K_0$ 1.4158 |
| 694.511 | 4.2 | 1.4149 | 0.01 | 11558.5 | $C_{40}H_{72}NO_6P^{\cdot -}$ | $[M+H]^+$ | Fragment | 753.5922 m/z, $1/K_0$ 1.4158 |
| 753.5922 | 4.2 | 1.4147 | 0.01 | 208.8554 | $C_{43}H_{81}N_2O_6P$ | $[M+H]^+$ | Precursor | 753.5922 m/z, $1/K_0$ 1.4158 |

**Supplementary Fig. S4: Multiplexed lipid precursor identification using positive ion mode iprm-PASEF profiling of dye- and IF-stained human hippocampal brain tissue section.** Two distinct precursor ions were simultaneously isolated within a single iprm-PASEF window in positive-ion mode MALDI and successfully resolved using ion mobility separation. (**a**) The precursor at $m/z^+$ 697.48 ($1/K_0$ ~1.33 V·s/cm²) was identified as phosphatidic acid PA(16:0/20:4) based on characteristic fragment ions including $m/z^+$ 441.23 and 239.24. (**b**) The precursor at $m/z^+$ 753.59 ($1/K_0$ ~1.41 V·s/cm²) was identified as sphingomyelin SM(d18:1/20:3) with diagnostic fragments at $m/z^+$ 694.51, 650.48, and 570.52. Heat maps display ion mobility ($1/K_0$) versus *m/z* with corresponding fragment ion tables showing mass accuracy, mobility values, intensities, chemical formulas, adduct types, and isolation window assignments. Colored markers indicate precursor (yellow and boxed) and fragment ions used for structural identification.

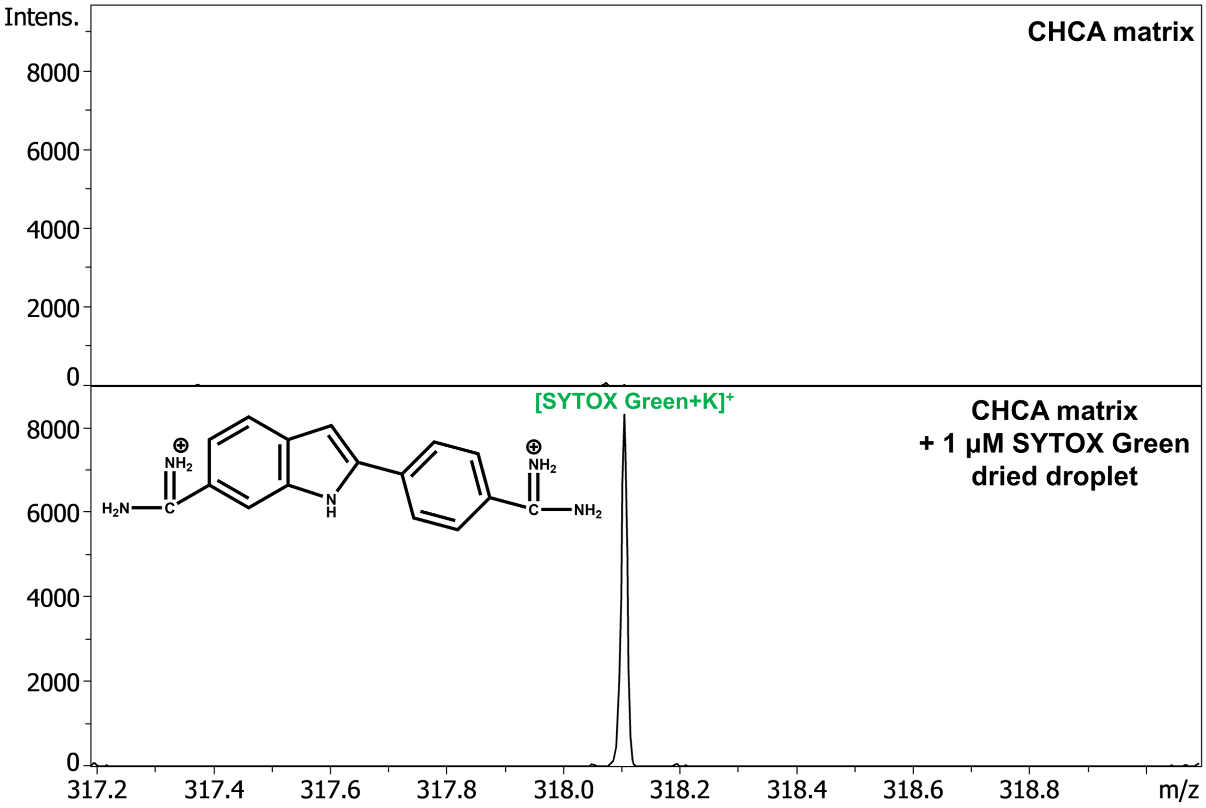

**<u>Supplementary Fig. S5:</u> Validation of SYTOX Green detection by MALDI-MS by dried droplet spot testing.** Mass spectra comparing CHCA matrix alone (top) versus CHCA matrix sprayed onto a 1 µM SYTOX Green dried droplet (bottom) in the *m/z* range of 317.2–319.0 Da. A SYTOX Green droplet was spotted onto the target plate and subsequently coated with CHCA matrix at 0.9 µg/mm² via robotic spray deposition prior to MALDI-MS acquisition. The potassium adduct of SYTOX Green ($[M+K]^+$, $m/z^+$ 318.1) was detected exclusively in the SYTOX Green droplet, confirming that this ion signal originated specifically from the nuclear stain and not from matrix-related interference. This validation supports the SYTOX Green assignment of $m/z^+$ 318.1 signals observed in FluoMALDI tissue imaging experiments.

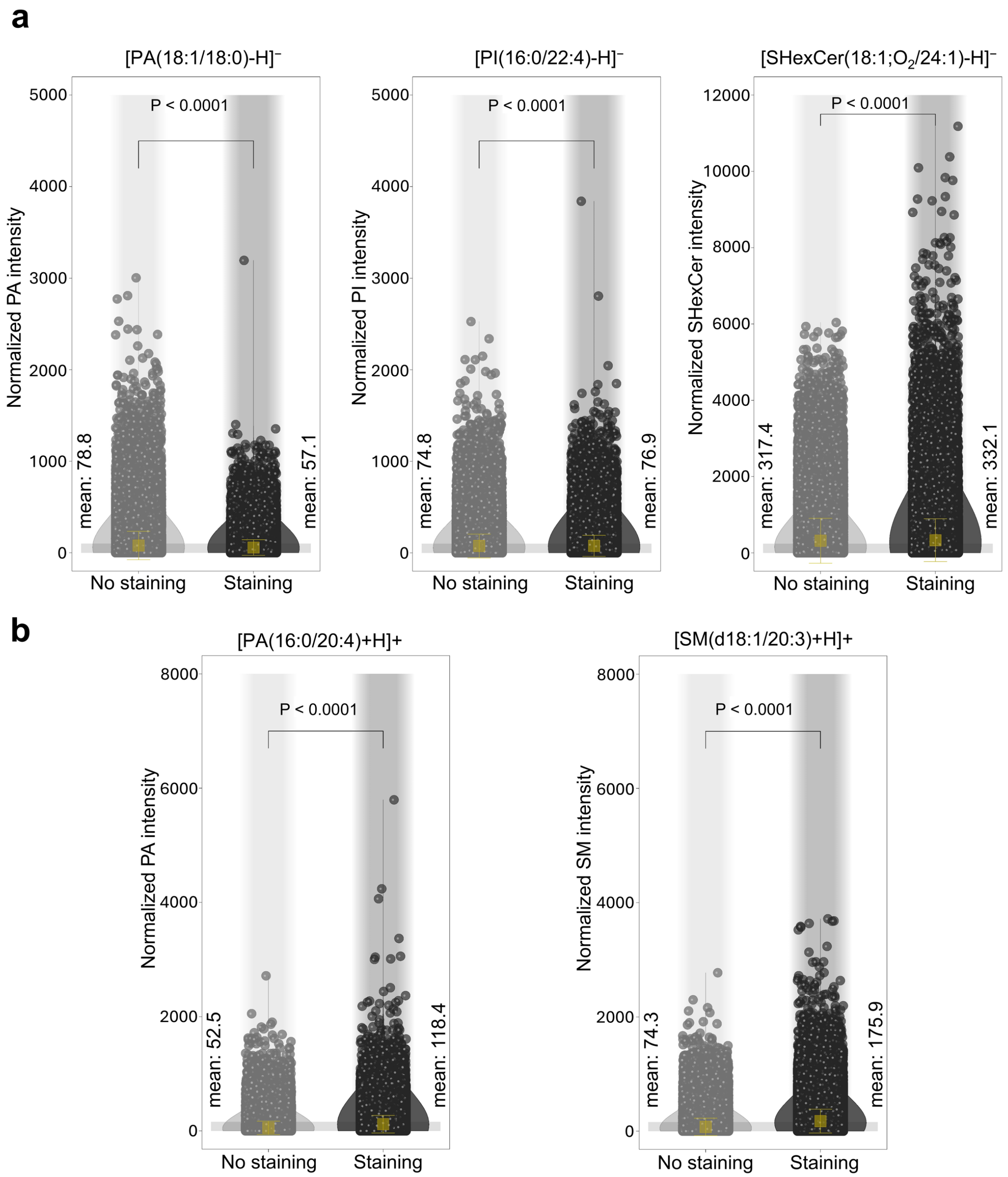

**<u>Supplementary Fig. S6:</u> Immunofluorescence staining preserves lipid molecular integrity for dual-modality FluoMALDI imaging.** Violin plots comparing normalized signal intensities of lipid species between unstained (No Staining) and immunofluorescence-stained (Staining) adjacent arterial regions in human hippocampal tissue sections. Data are presented on a per-pixel basis, with each open circle plot representing an individual pixel at a spatial resolution of 5 µm. (**a**) Lipid species detected in negative ion mode: phosphatidic acid $[PA(18:1/18:0)-H]^{-}$, phosphatidylinositol $[PI(16:0/22:4)-H]^{-}$, and sulfatide $[SHexCer(18:1;O_2/24:1)-H]^{-}$. Despite statistically significant differences (Mann–Whitney U test, $p < 0.0001$), mean signal intensities remained comparable between stained and unstained conditions (PA: 57.1 versus 78.8; PI: 76.9 versus 74.8; SHexCer: 332.1 versus 317.4). (**b**) Lipid species detected in positive ion mode: phosphatidic acid $[PA(16:0/20:4)+H]^{+}$ and sphingomyelin $[SM(d18:1/20:3)+H]^{+}$. Stained tissue sections showed higher mean intensities compared to unstained sections (PA: 118.4 versus 52.5; SM: 175.9 versus 74.3; Mann–Whitney U test, $p < 0.0001$). All results are also presented as mean ± standard deviation (SD), where the mean value for each group is indicated by yellow squares.

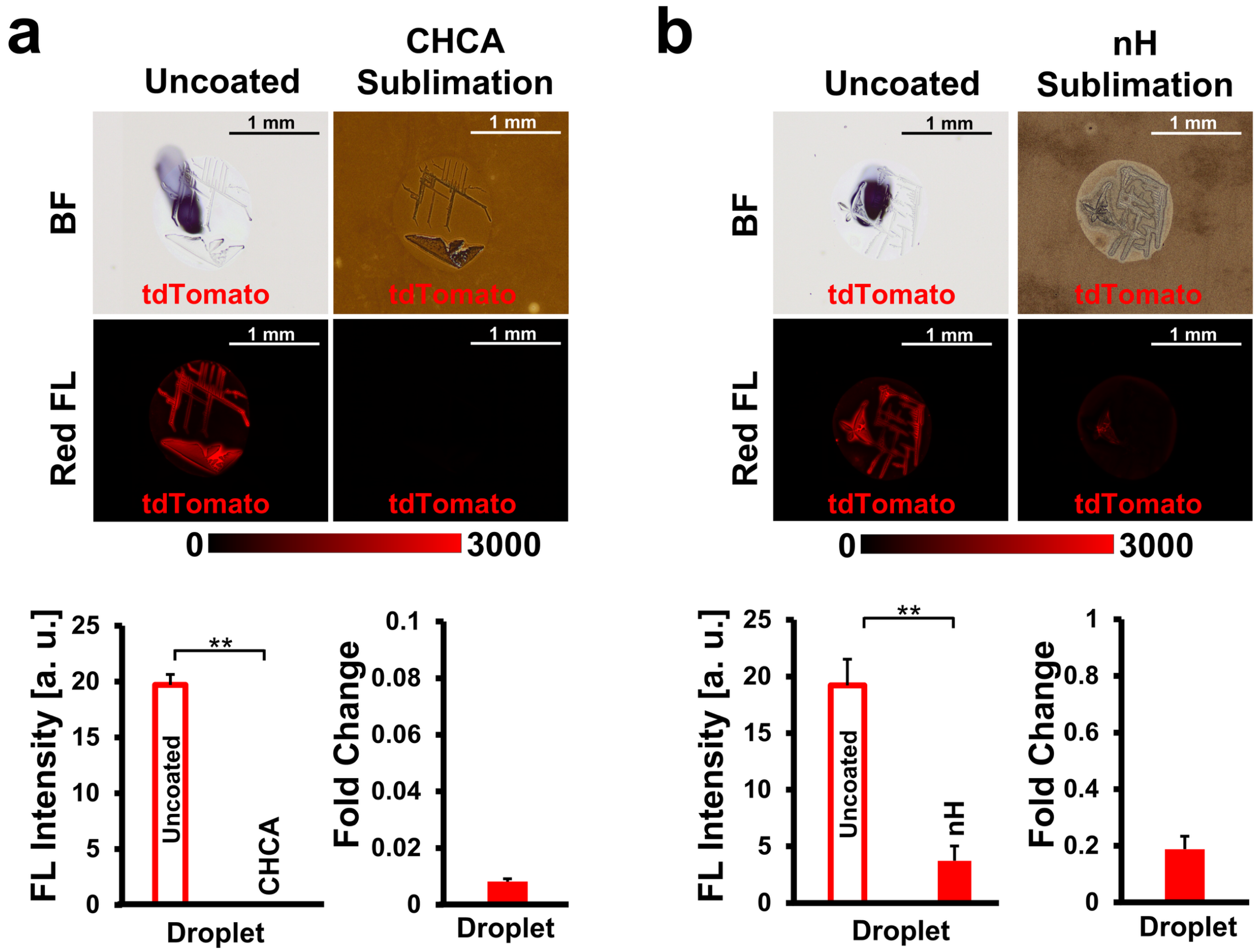

<u>**Supplementary Fig. S7:**</u> **MALDI matrix sublimation coating reduces fluorescence detection of tdTomato fluorescent protein.** (**a**) Effect of CHCA matrix coating on tdTomato fluorescence. Top: Brightfield (BF) and red fluorescence (Red FL) images of dried tdTomato droplets (1 µM) before (left, Uncoated) and after (right) CHCA sublimation coating. Bottom: Quantification of fluorescence intensity (left) and fold change relative to uncoated control (right) comparing uncoated versus CHCA-coated conditions. (**b**) Effect of nH matrix coating on tdTomato fluorescence. Top: Brightfield (BF) and red fluorescence (Red FL) images of dried tdTomato droplets (1 µM) before (left, Uncoated) and after (right) nH sublimation coating. Bottom: Quantification of fluorescence intensity (left) and fold change relative to uncoated control (right) comparing uncoated versus nH-coated conditions. Error bars represent standard deviation (n = 3 droplets).

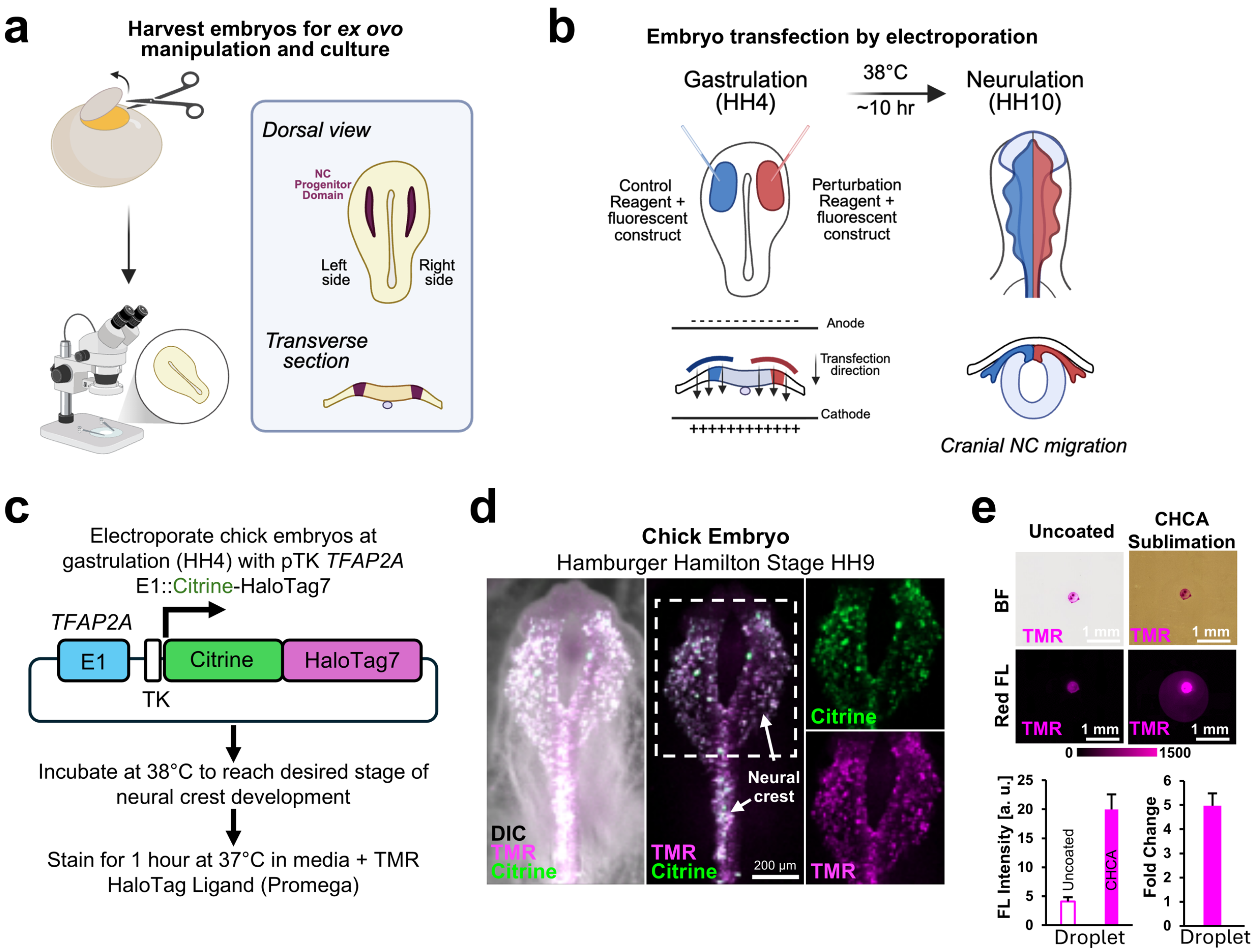

**Supplementary Fig. S8: HaloTag-TMR-based genetic labeling of neural crest cells in chick embryos and fluorescence enhancement by CHCA matrix coating.** (**a**) Chick embryos are harvested at gastrulation and cultured *ex ovo* for manipulation under a stereomicroscope. Right, dorsal view and transverse section of a gastrula-stage embryo showing the left and right sides and the neural crest (NC) progenitor domains (maroon). (**b**) *Ex ovo* electroporation at gastrulation (Hamburger–Hamilton stage 4, HH4). A control reagent with a fluorescent construct is delivered to the left side (blue) and a perturbation reagent with a fluorescent construct to the right side (red), as in the *SMPD3* gain-of-function experiments in Fig. 3, so that each embryo contains its own control. The transfection direction between the electrodes is indicated. Embryos are incubated at 38 °C until neurulation (HH10), when cranial NC cells on both sides migrate (bottom right, transverse section). (**c**) The pTK *TFAP2A* E1::Citrine-HaloTag7 construct drives a Citrine-HaloTag7 fusion under the NC-specific *TFAP2A* E1 enhancer (E1) and the thymidine kinase (TK) minimal promoter. Electroporated embryos are incubated at 38 °C to the desired stage of NC development and stained with TMR HaloTag ligand (Promega) for 1 h at 37 °C. (**d**) Validation of NC labeling in a chick embryo at HH9. Left, merged differential interference contrast (DIC), TMR (magenta) and Citrine (green) channels. Middle, merged TMR and Citrine fluorescence, in which the two signals colocalize in NC cells (arrows). Right, individual Citrine and TMR channels of the boxed region. Scale bar: 200 µm. (**e**) CHCA matrix coating enhances TMR fluorescence. Top, brightfield (BF) and red fluorescence (Red FL, magenta) images of dried TMR droplets before (Uncoated) and after CHCA sublimation coating. The color bar shows the display range (0 to 1500, arbitrary units, a.u.) of both Red FL images. Scale bars: 1 mm. Bottom, fluorescence (FL) intensity (a.u.) and fold change of the droplets, showing an approximately 5-fold increase after CHCA coating. Data are mean ± s.d. (n = 3 droplets).

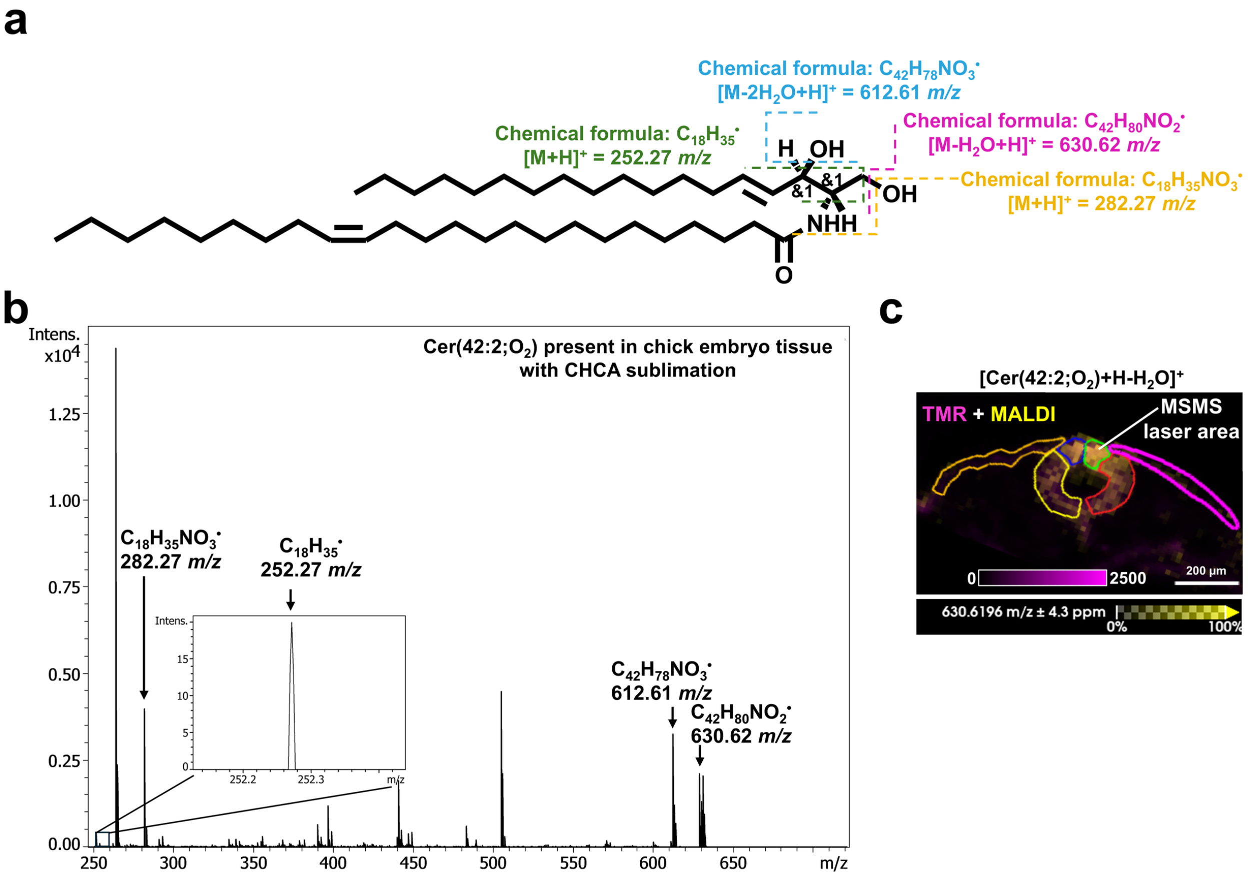

**<u>Supplementary Fig. S9:</u> On-tissue MS/MS identification of Cer(42:2;O$_2$) from chick embryo tissue.** (**a**) Chemical structure of Cer(42:2;O$_2$) with the precursor ion at *m/z*$^+$ 630.62. Annotations indicate characteristic fragmentation sites and (**b**) their corresponding fragment ions: precursor ion at *m/z*$^+$ 630.62, fragments at *m/z*$^+$ 612.61, 282.27, and sphingoid base fragment at *m/z*$^+$ 252.27. (**c**) FluoMALDI-MSI overlay image of TMR fluorescence and MALDI ion image showing the MS/MS laser interrogation area (highlighted in green) targeting the GOF neural crest region with the highest Cer(42:2;O$_2$) abundance, corresponding to the distribution shown in Figure 3. Scale bar: 200 µm.

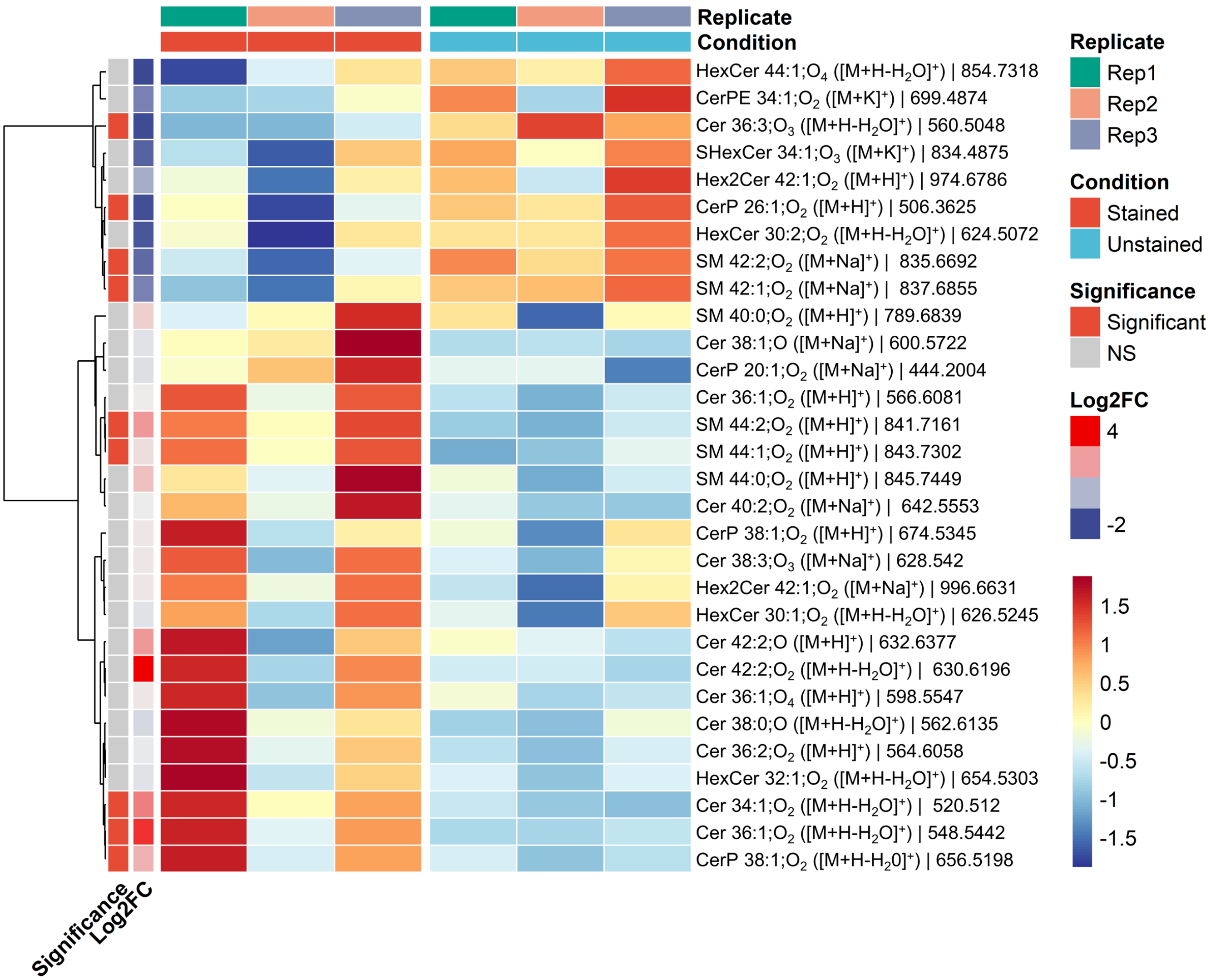

**<u>Supplementary Fig. S10:</u> Heatmap showing comparable ceramide-related lipid profiles between HaloTag-stained and unstained embryo samples.** Lipids with more than two missing values across the six samples were excluded prior to analysis. Remaining values below the limit of detection were inferred by using half-minimum substitution with a small uniform jitter added to avoid numerical ties. The displayed top 30 lipids were selected by ranking their π-score, a composite metric combining statistical significance and fold-change magnitude. Intensities were $\log_2$-transformed and row-wise Z-score normalized (red is high relative abundance, blue is low relative abundance). Row annotations indicate statistical significance determined by Empirical Bayes linear modeling (limma; unadjusted $p < 0.05$ AND $|\log_2 FC| \geq 1$; red is significant, gray is not significant) alongside the $\log_2$ fold change (red is enriched in HaloTag-stained, blue is enriched in unstained). Column annotations denote the experimental condition ($n = 3$ per group; red is HaloTag-stained, cyan is unstained) and replicate identity. Lipids (rows) were hierarchically clustered using the Ward.D2 agglomeration method based on Pearson correlation distance. Lipid class abbreviations: Cer, ceramide; CerP, ceramide-1-phosphate; CerPE, ceramide phosphoethanolamine; SM, sphingomyelin; HexCer, hexosylceramide; Hex2Cer, dihexosylceramide; SHexCer, sulfatide.

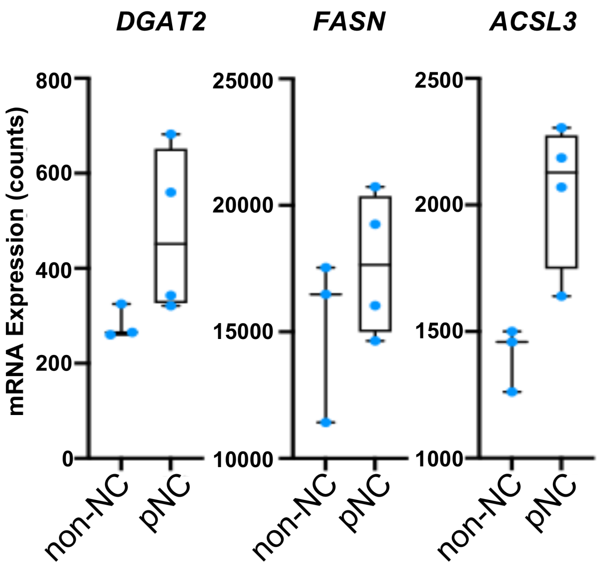

**Supplementary Fig. S11: Upregulation of lipid metabolic genes in premigratory neural crest cells revealed by bulk RNA sequencing.** Bulk mRNA sequencing analysis of isolated pre-migratory neural crest (pNC) cells compared with surrounding non–neural crest (non-NC) tissue revealed an increased expression of lipid metabolism–related genes. Data analysis was performed using previously published sequencing datasets[1]. mRNA levels for *DGAT2*, *FASN*, and *ACSL3* are displayed as box plots with individual data points representing technical replicates. pNC samples (n=4 technical replicates) exhibited higher transcript abundance of these genes relative to non-NC samples (n=3 technical replicates), consistent with enhanced fatty acid synthesis and lipid remodeling in pre-migratory neural crest cells.

a

b

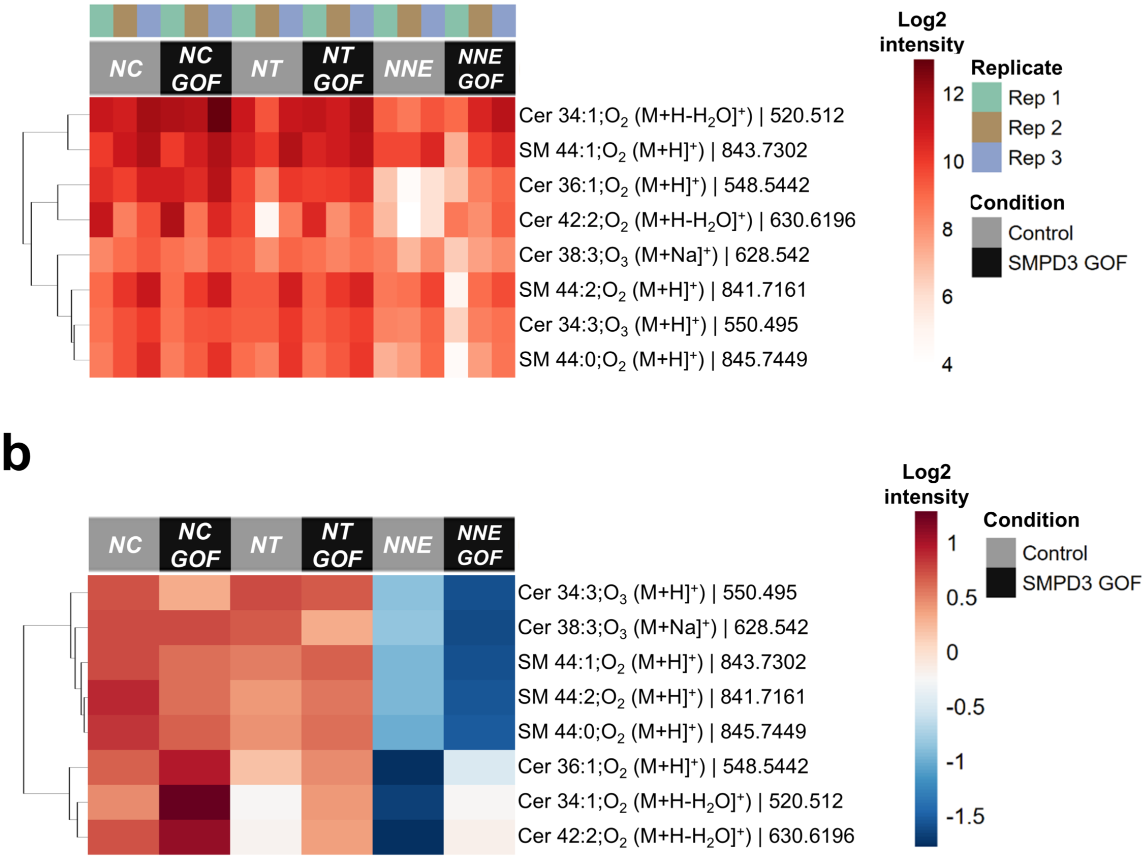

**<u>Supplementary Fig. S12:</u> Sphingolipid remodeling in *SMPD3* gain-of-function embryonic tissues.** Heatmaps displaying log2-transformed intensities of sphingomyelin (SM) and ceramide (Cer) species in control versus *SMPD3* gain-of-function (GOF) conditions across three embryonic tissue types: neural crest (NC), neural tube (NT), and non-neural ectoderm (NNE). (**a**) Individual technical replicates (n = 3). (**b**) Averaged values across 3 replicates. In both panels, columns are organized by tissue type with control (gray) and *SMPD3* GOF (black) conditions paired. Rows, representing individual lipid species (identified by lipid class, acyl chain composition, detected ion adduct, and *m/z* value) are hierarchically clustered based on abundance patterns. Consistent with elevated sphingomyelinase activity, SM species show reproducible depletion in *SMPD3* GOF samples across all tissues and replicates.

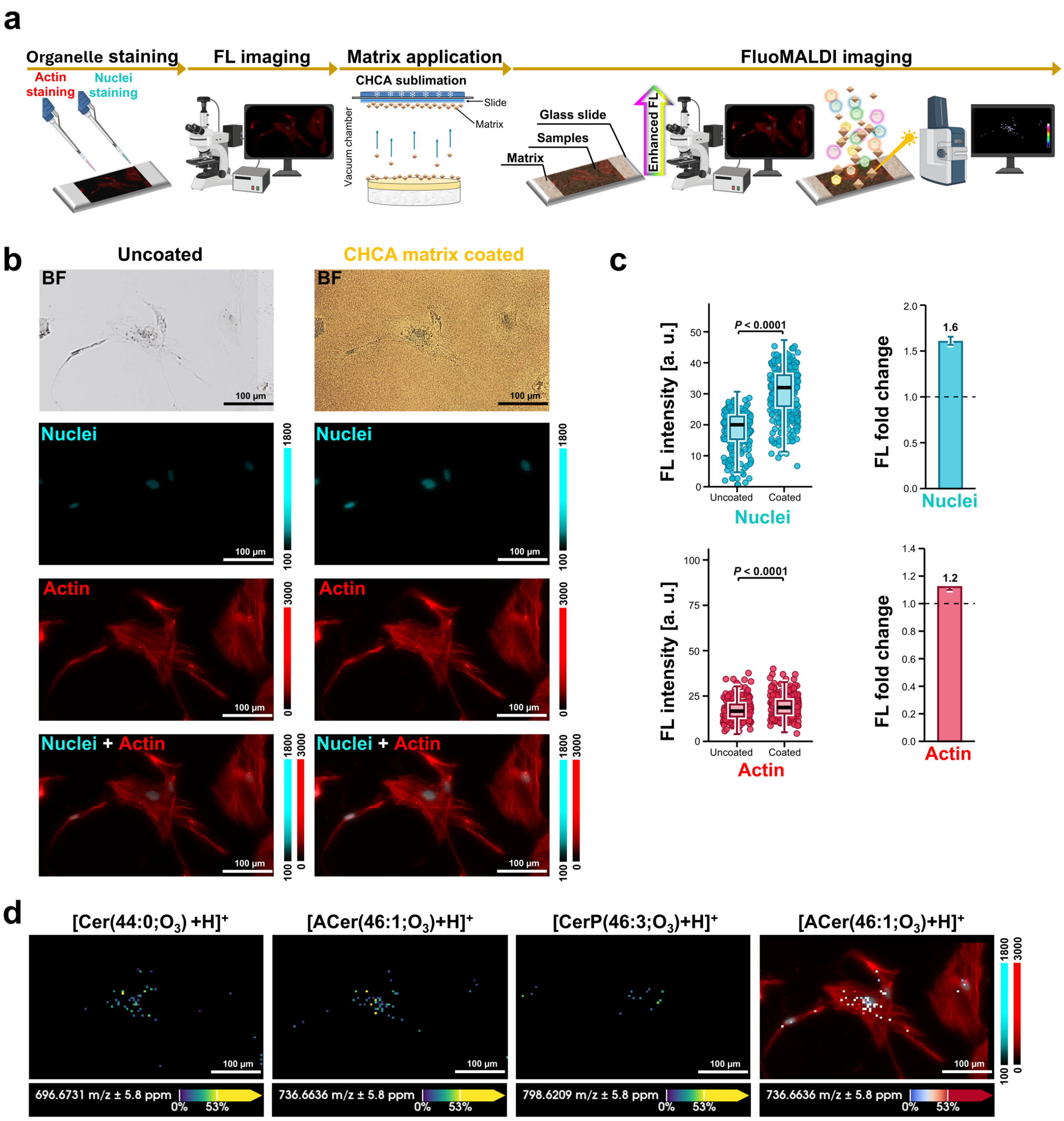

**<u>Supplementary Fig. S13:</u> Dual-channel dye-FluoMALDI imaging of human iPSC-derived astrocytes with subcellular organelle staining.** (**a**) Schematic workflow for dye-FluoMALDI-MSI analysis focusing on fluorescence enhancement. Human iPSC-derived astrocytes were cultured in chamber slides and labeled with organelle-specific fluorescent dyes. Optical microscopy was performed to compare the fluorescence of uncoated and coated cells. CHCA matrix was applied by sublimation. (**b**) Representative brightfield (BF) and fluorescence images of human iPSC-derived astrocytes before (left column, uncoated) and after (right column, CHCA matrix-coated) matrix deposition. Fluorescence channels are displayed as follows: nuclei (SYTOX Green) in cyan and filamentous actin (Phalloidin-Alexa Fluor 635) in red. Bottom row shows merged multichannel images. Scale bars: 100 µm. (**c**) Fluorescence intensity (a.u.) of nuclei and actin before and after CHCA sublimation (left) and corresponding fold change (right, ratio of medians, 95% bootstrap confidence interval, the dashed line marks 1). Each point is one pixel. Boxes show the median and interquartile range, and whiskers show 1.5 times the interquartile range. Statistical comparisons used two-sided Mann-Whitney U tests, $P < 0.0001$ for all channels. Effect sizes are in Source Data. **(d)** Integrated fluorescence and MALDI-MSI of human iPSC-derived astrocytes. MALDI ion images show the spatial distribution of a ceramide species (Cer, $[Cer(44:0;O_3) + H]^+$ at 696.6731 *m/z*), an acyl ceramide species (ACer, $[ACer(46:1;O_3) + H]^+$ at 736.6636 *m/z*), and a ceramide phosphate species (CerP, $[CerP(46:3;O_3) + H]^+$ at 798.6209 *m/z*). The rightmost panel displays a co-registered overlay of fluorescence microscopy and MALDI-MSI data.

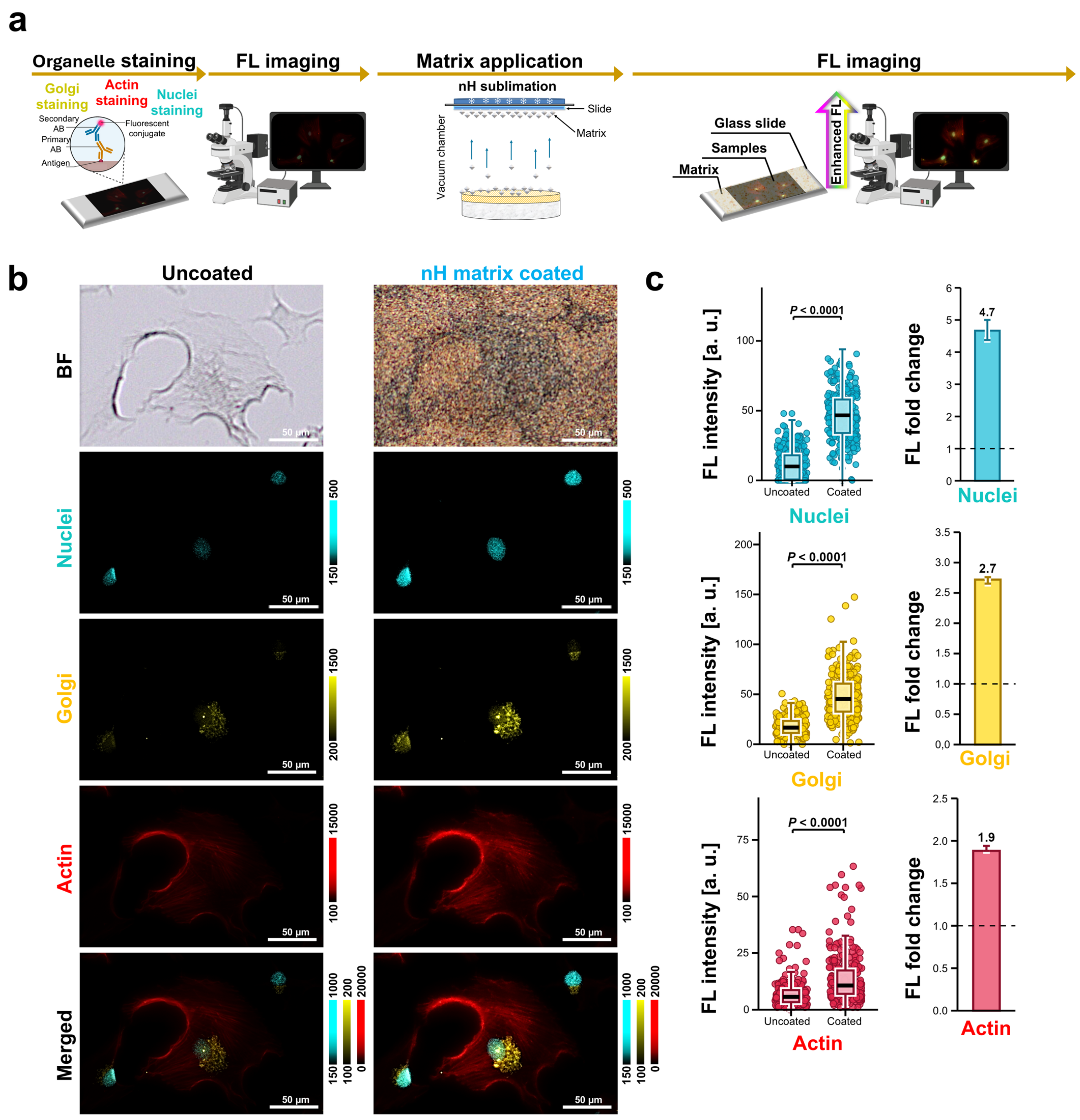

**Supplementary Fig. S14:** **Three-channel dye- and immuno-FluoMALDI imaging of human iPSC-derived astrocytes: fluorescence enhancement of nuclei, Golgi, and filamentous actin following nH matrix coating**. (**a**) Schematic workflow for dye- and immuno-FluoMALDI fluorescence enhancement analysis. Human iPSC-derived astrocytes were cultured in chamber slides and labeled with organelle-specific fluorescent dyes. Optical microscopy was performed to compare the fluorescence of uncoated and coated cells. nH matrix was applied by sublimation. (**b**) Representative brightfield (BF) and fluorescence images of astrocytes before (left column, uncoated) and after (right column, nH matrix-coated) matrix deposition. Fluorescence channels are displayed as follows: nuclei (SYTOX Green) in cyan, Golgi (A568-conjugated antibody) in yellow, and filamentous actin (Phalloidin-Alexa Fluor 647) in red. Bottom row shows merged multichannel images. Scale bars: 50 µm. (**c**) Fluorescence intensity (a.u.) of nuclei, Golgi, and actin before and after nH sublimation (left) and corresponding fold change (right, ratio of medians, 95% bootstrap confidence interval, the dashed line marks 1). Each point is one pixel. Boxes show the median and interquartile range, and whiskers show 1.5 times the interquartile range. Statistical comparisons used two-sided Mann-Whitney U tests, $P < 0.0001$ for all channels. Effect sizes are in Source Data.

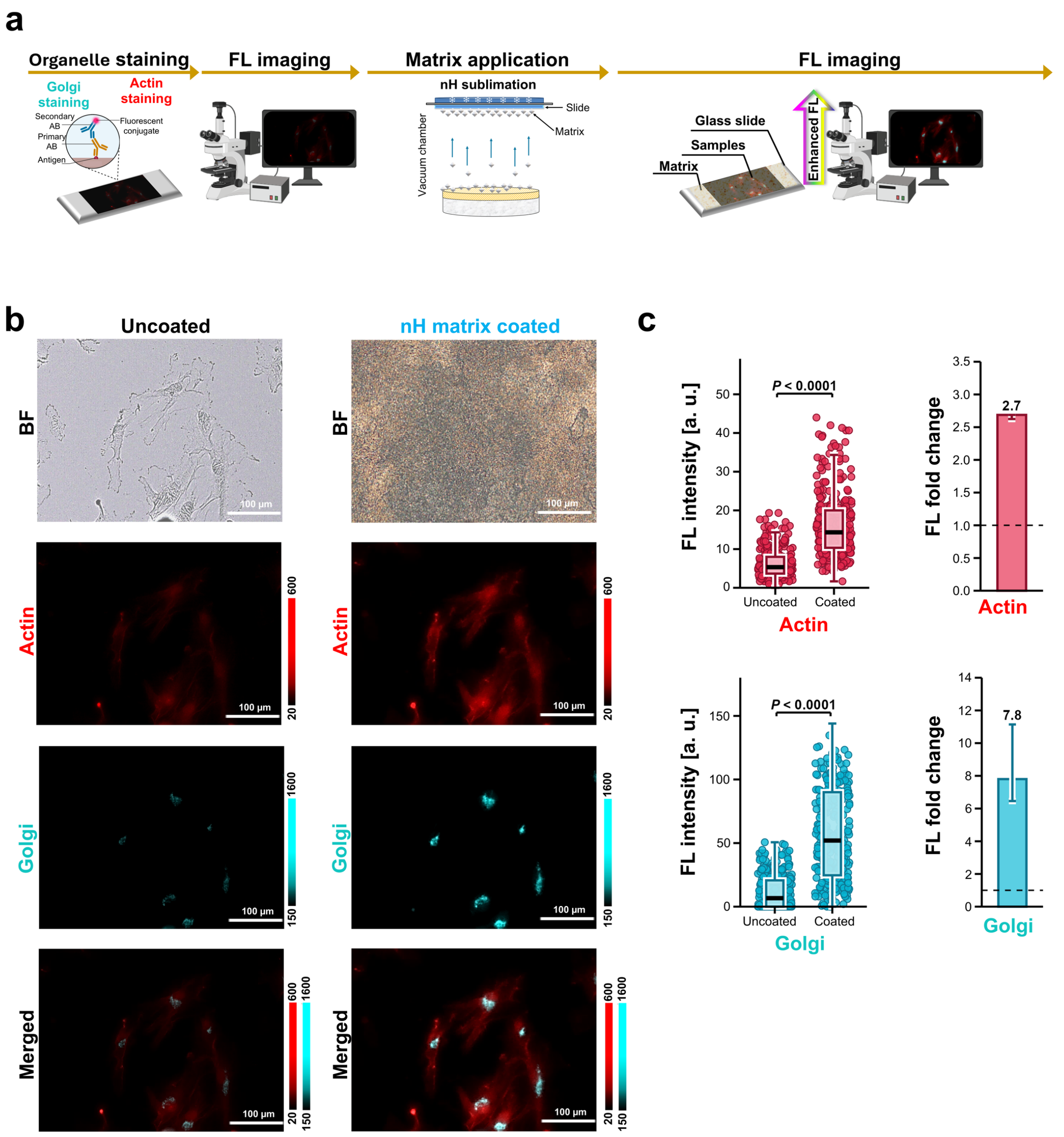

**Supplementary Fig. S15: Dual-channel dye- and immuno-FluoMALDI imaging of human iPSC-derived astrocytes: fluorescence enhancement of Golgi and filamentous actin following nH matrix coating.** (**a**) Schematic workflow for dye- and immuno-FluoMALDI fluorescence enhancement analysis. Human iPSC-derived astrocytes were cultured in chamber slides and labeled with organelle-specific fluorescent dyes. Optical microscopy was performed to compare the fluorescence of uncoated and coated cells. nH matrix was then applied by sublimation. (**b**) Representative brightfield (BF) and fluorescence images of astrocytes before (left column, uncoated) and after (right column, nH matrix-coated) matrix deposition. Fluorescence channels are displayed as follows: Golgi (A488-conjugated antibody) in cyan, and actin (Phalloidin-Alexa Fluor 635) in red. Bottom row shows merged multichannel images. Scale bars: 100 μm. (**c**) Fluorescence intensity (a.u.) of Golgi and actin before and after nH sublimation (left) and corresponding fold change (right, ratio of medians, 95% bootstrap confidence interval, the dashed line marks 1). Each point is one pixel. Boxes show the median and interquartile range, and whiskers show 1.5 times the interquartile range. Statistical comparisons used two-sided Mann-Whitney U tests, $P < 0.0001$ for all channels. Effect sizes are in Source Data.

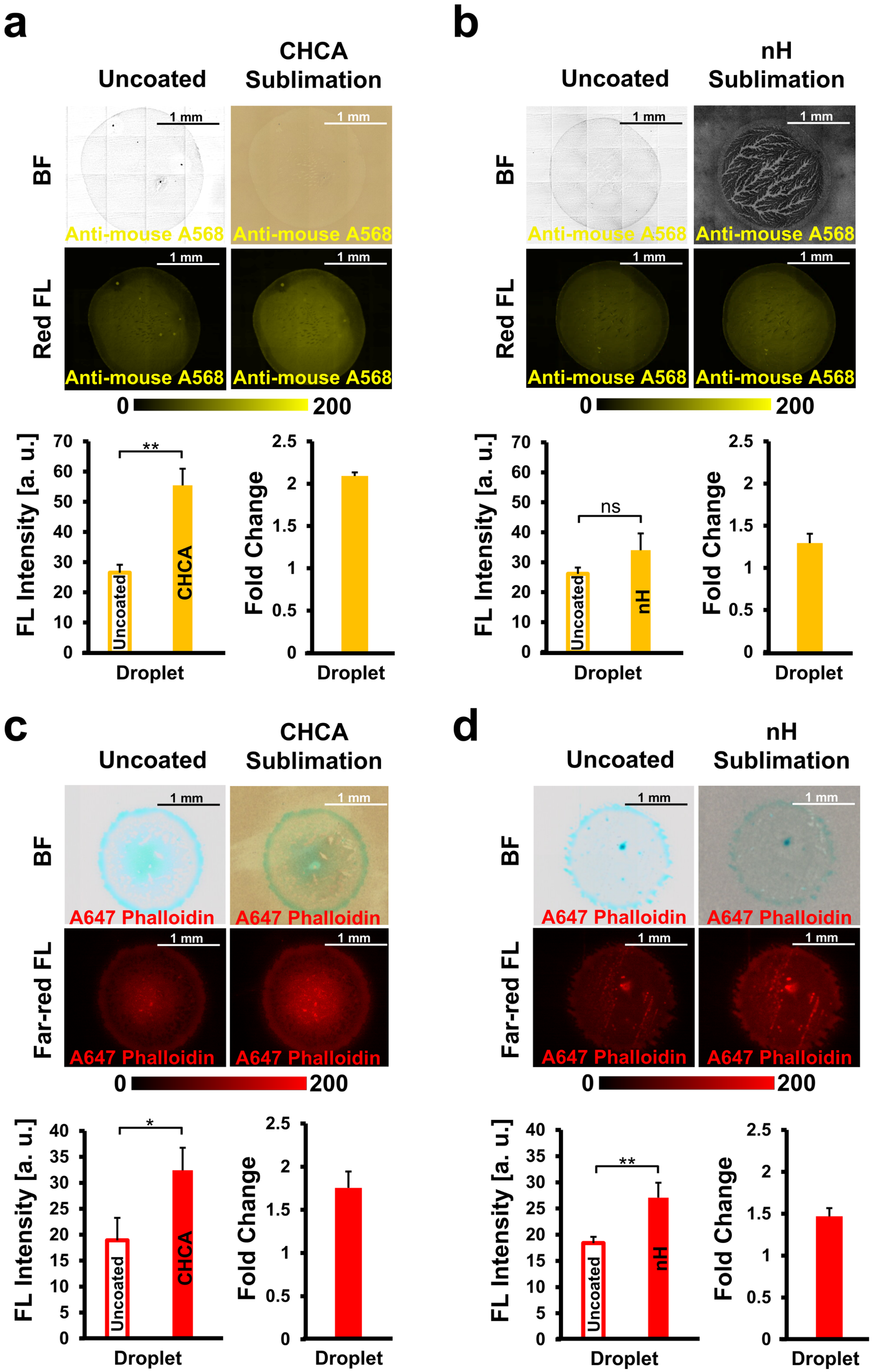
a
Uncoated
CHCA Sublimation
BF
Red FL
1 mm
Anti-mouse A568
0
200
FL Intensity [a. u.]
Fold Change
Droplet
**
Uncoated
CHCA
b
Uncoated
nH Sublimation
ns
nH
c
Uncoated
CHCA Sublimation
Far-red FL
A647 Phalloidin
*
d
Uncoated
nH Sublimation
**

**Supplementary Fig. S16: Matrix sublimation coating effects on fluorescent dried droplet detection of A568-conjugated antibody and A647-conjugated phalloidin. Top panels** (**a-d**): Brightfield (BF, top) and fluorescence (FL, bottom) images of uncoated (left) and CHCA (**a**, **c**) or nH (**b**, **d**) sublimation coated (right) pure dried A568-conjugated anti-mouse antibody (20 µg/mL; **a**, **b**) or A647-conjugated phalloidin (2 µM; **c**, **d**) droplets. Scale bars: 1 mm. Color scales indicate fluorescence intensity. **Bottom panels** (**a-d**): Quantification of fluorescence intensity (left graphs) and fold change (right graphs) of dried droplets comparing uncoated versus matrix-coated conditions. Error bars represent standard deviation (n = 3 droplets).

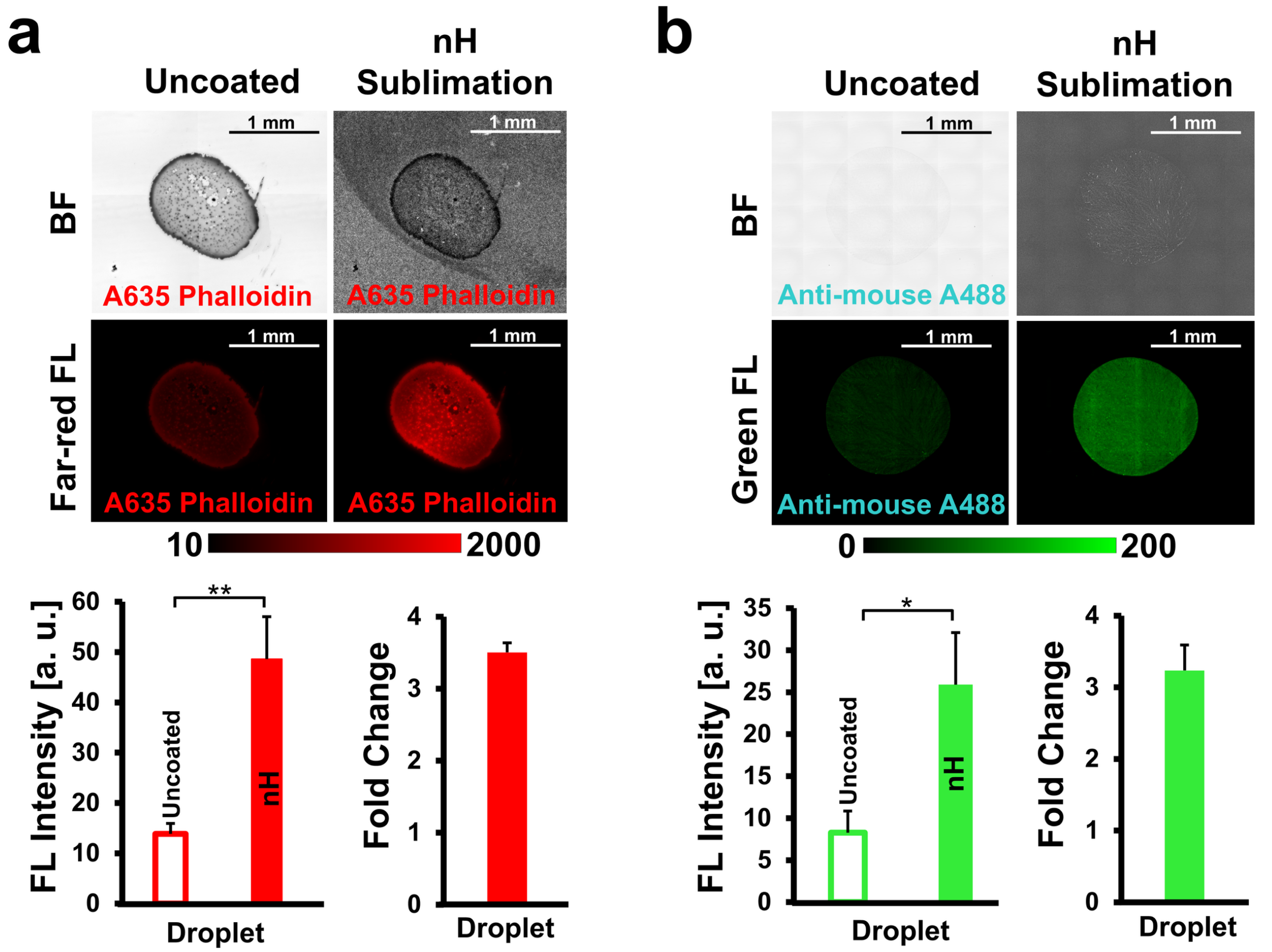

<u>Supplementary Fig. S17:</u> **Matrix sublimation coating effects on fluorescent dried droplet detection of A635-conjugated phalloidin and A488-conjugated antibody. Top panels** (**a, b**): Brightfield (BF, top) and fluorescence (FL, bottom) images of uncoated (left) and nH (**a**, **b**) sublimation coated (right) pure dried A635-conjugagted phalloidin (2 µM; **a**) or A488-conjugated anti-mouse antibody (20 µg/mL; **b**) droplets. Scale bars: 1 mm. Color scales indicate fluorescence intensity. **Bottom panels** (**a**, **b**): Quantification of fluorescence intensity (left graphs) and fold change (right graphs) of dried droplets comparing uncoated versus matrix-coated conditions. Error bars represent standard deviation (n = 3 droplets).

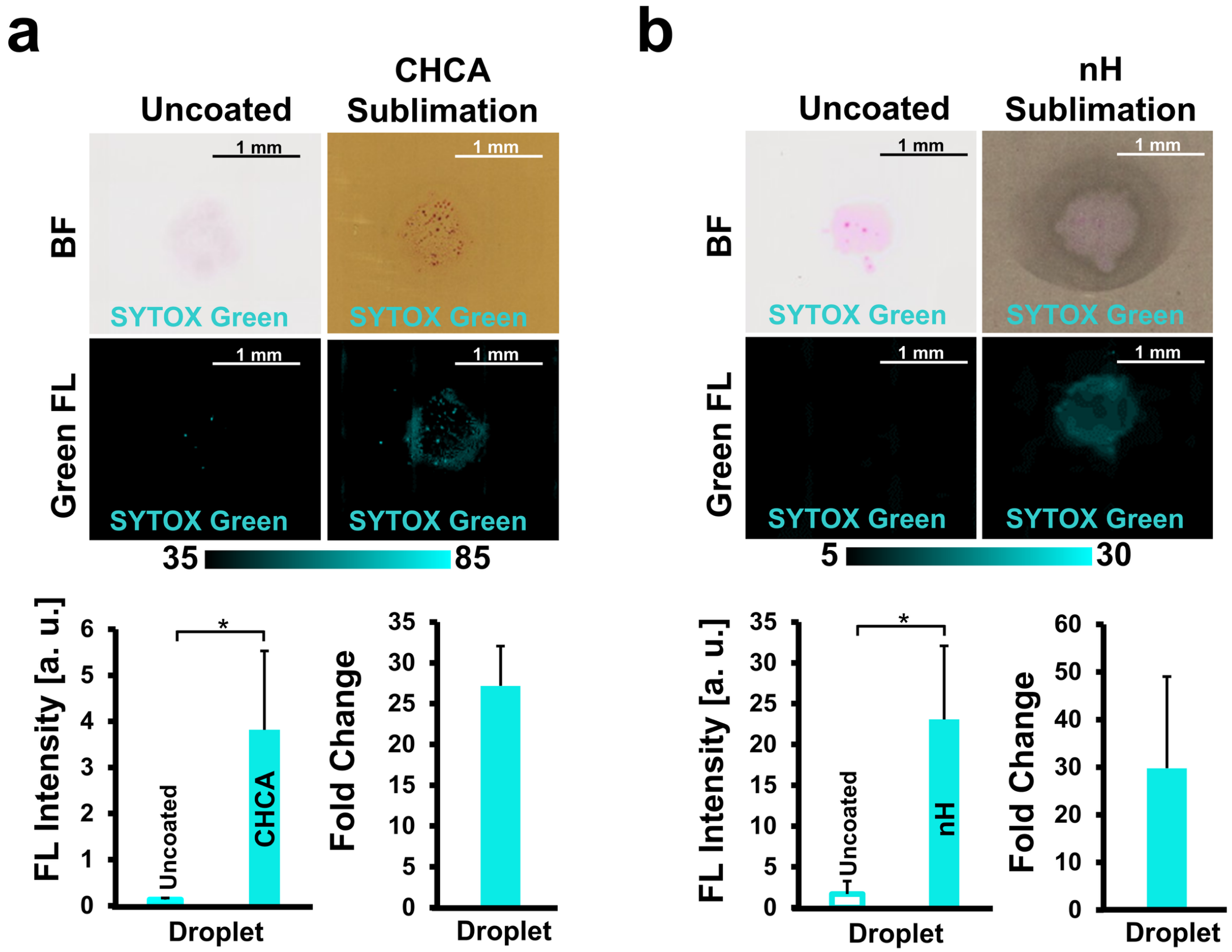

**<u>Supplementary Fig. S18:</u> Matrix sublimation coating effects on fluorescent dried droplet detection of SYTOX Green. Top panels** (**a, b**): Brightfield (BF, top) and fluorescence (FL, bottom) images of uncoated (left) and CHCA (**a**) or nH (**b**) sublimation coated (right) pure dried SYTOX Green (1 µM; **a**, **b**) droplets. Scale bars: 1 mm. Color scales indicate fluorescence intensity. **Bottom panels** (**a**, **b**): Quantification of fluorescence intensity (left graphs) and fold change (right graphs) of dried droplets comparing uncoated versus matrix-coated conditions. Error bars represent standard deviation (n = 3 droplets).

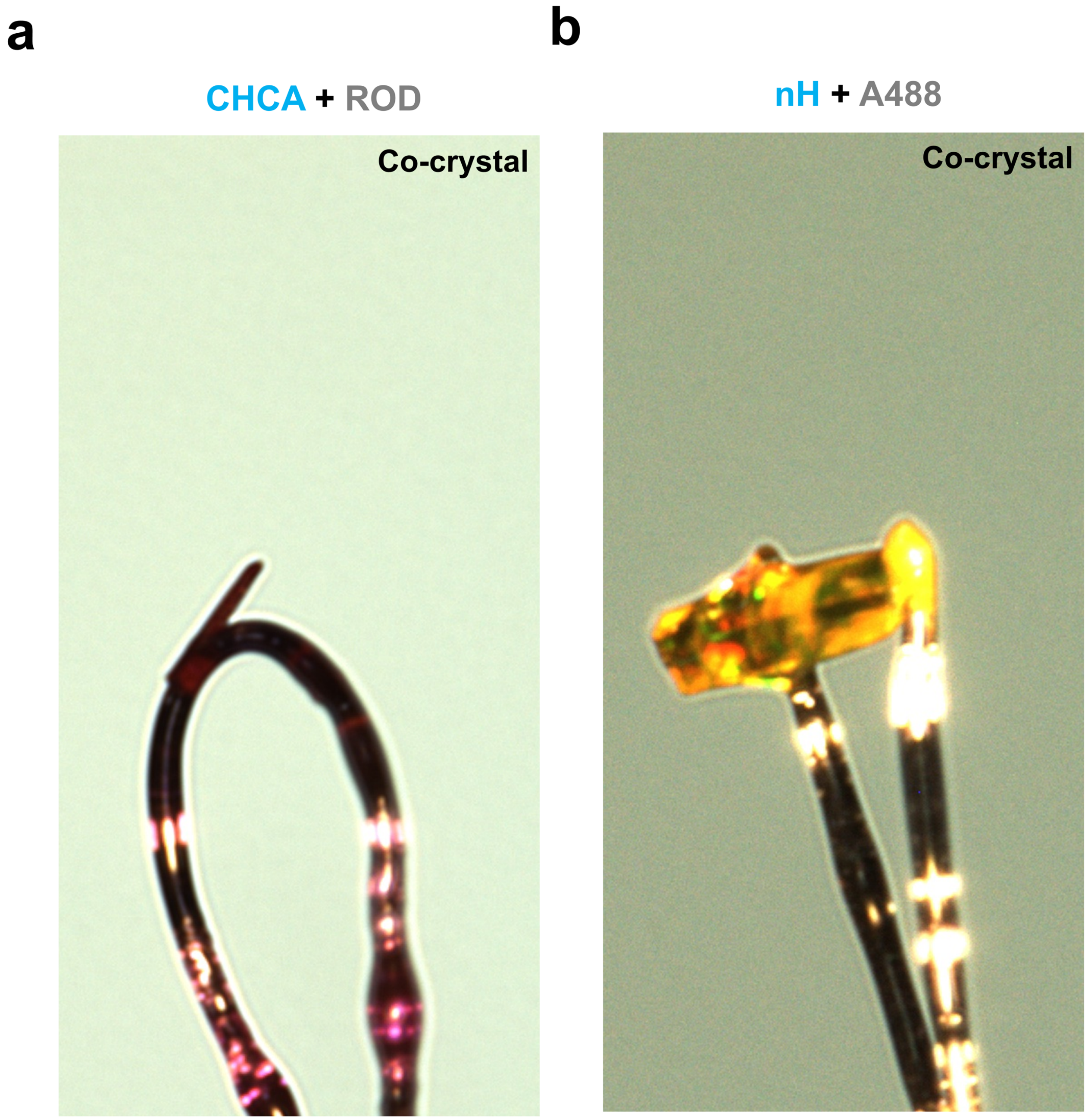

**<u>Supplementary Fig. S19:</u> Representative single co-crystals used for X-ray diffraction.** Optical micrographs of (**a**) the CHCA–rhodamine B (ROD) co-crystal with needle-like crystal morphology, and (**b**) the nH–Alexa Fluor 488 (A488) co-crystal with rod-like crystal morphology, mounted for diffraction data collection. Crystal coloration is consistent with fluorophore incorporation: the CHCA–ROD crystal is deep red-violet, while the nH–A488 crystal is amber-orange.

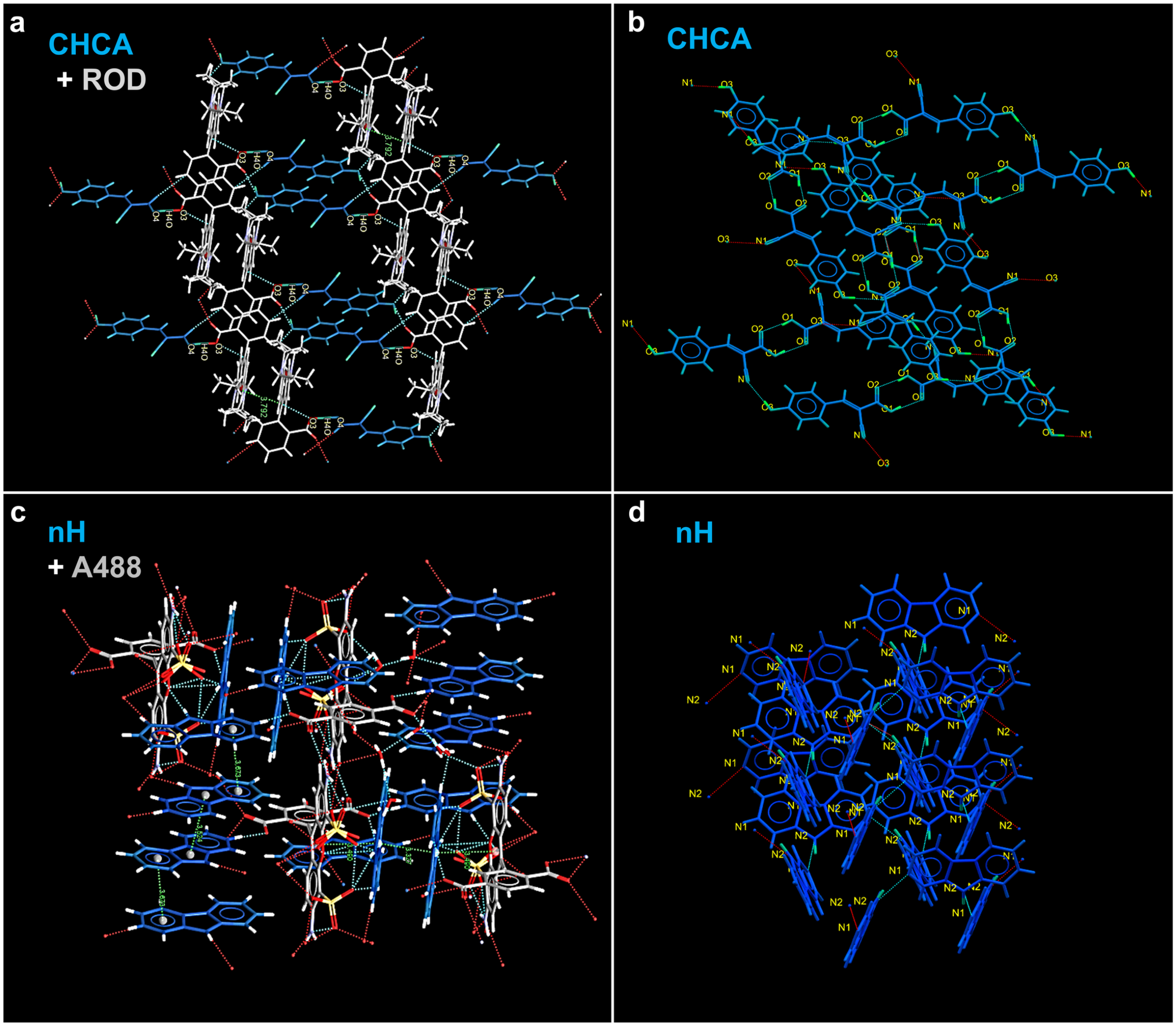

**<u>Supplementary Fig. S20:</u> Crystal packing of matrix–fluorophore co-crystals and matrix-only controls, determined by single-crystal X-ray diffraction (SCXRD).** Capped-stick representations from single-crystal X-ray diffraction. (**a**) CHCA–rhodamine B (ROD) co-crystal: a localized π–π interaction between a ROD pair (interplanar distance, ca. below 3.8 Å; least-squares planes calculated for C1–C13 and O1) bridged by CHCA through O–H···O hydrogen bonds (O4···O3, 2.4610(16) Å, O4–H···O3, 174(2)°; O6···O2, 2.6318(18) Å, O6–H···O2, 160(2)°). (**b**) CHCA alone (negative control): O–H···O hydrogen-bonded network (dashed blue lines), no π–π stacking. (**c**) nH–A488 co-crystal: hydrogen bonds (dashed blue lines, 2.65 Å) between the nH pyridine nitrogen and A488 carboxyl groups; a localized π–π interaction (centroid-to-centroid distance, <3.7 Å) between the nH indole ring and A488 xanthene core. (**d**) nH alone (negative control): hydrogen-bonded network (dashed blue lines), no π–π stacking. Structures rendered in Mercury[2].

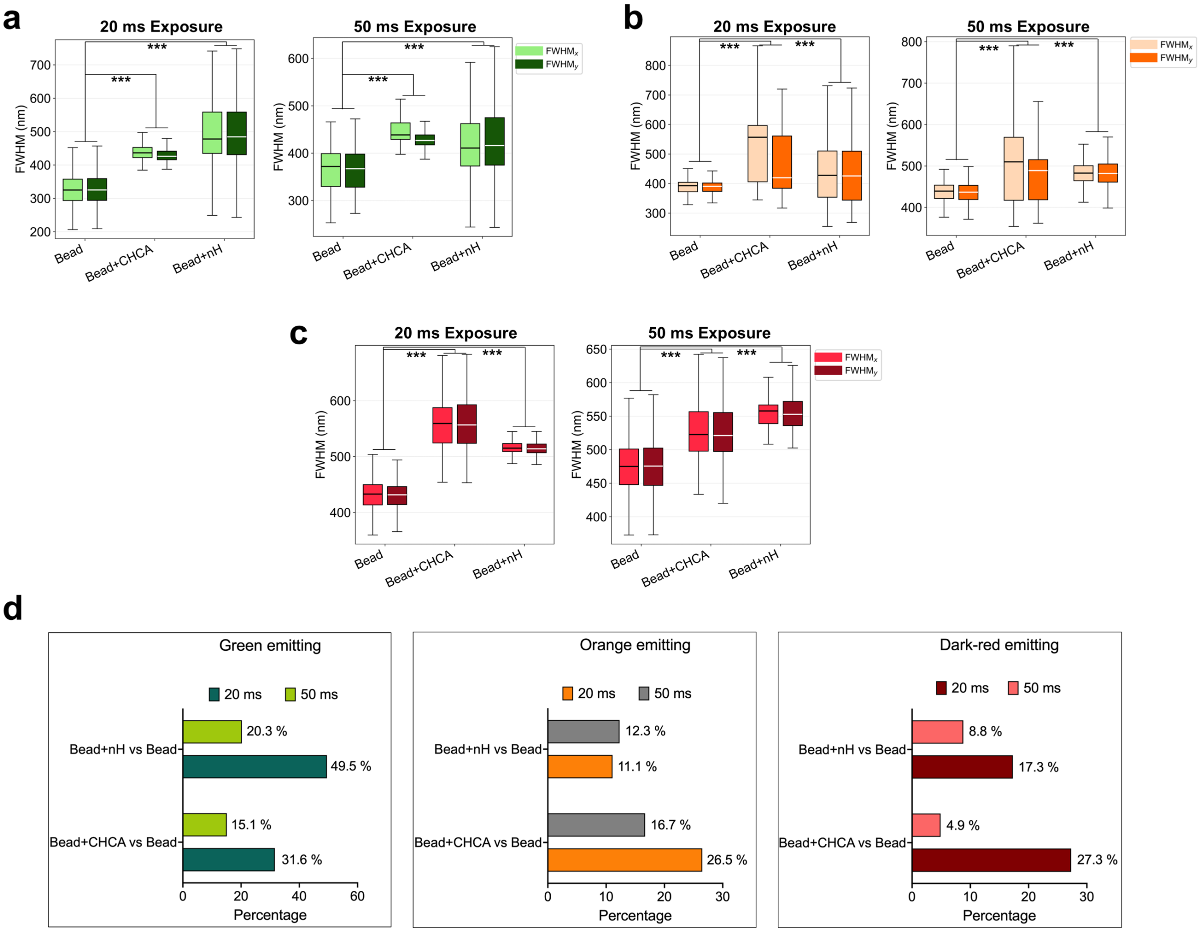

**<u>Supplementary Fig. S21:</u> Point spread function (PSF) measurements.** Full width at half maximum (FWHM) measurements of 200-nm-diameter fluorescent microspheres without matrix (Bead) and after sublimation coating of CHCA (Bead+CHCA) or nH (Bead+nH). (**a-c**) Distributions of lateral FWHM along the x and y directions for (**a**) green-emitting, (**b**) red-emitting, and (**c**) far-red-emitting beads acquired at 20-ms and 50-ms exposures. (**d**) Percentage increases in mean bead FWHM relative to the corresponding uncoated control at the same exposure time, calculated as 100 × (mean FWHM_matrix / mean FWHM_bead − 1), with per-bead FWHM defined as (FWHM_x + FWHM_y)/2. CHCA-associated increases at 20 ms and 50 ms were 31.6% and 15.1% (green), 26.5% and 16.7% (orange), and 27.3% and 4.9% (dark-red). The corresponding nH-associated increases were 49.5% and 20.3%, 11.1% and 12.3%, and 17.3% and 8.8%. Statistical comparisons used two-sided Mann-Whitney U tests on per-bead mean FWHM, comparing each matrix condition with the uncoated control within each exposure, with Holm correction. Boxes show the interquartile range (25th-75th percentiles) of the bead measurements.

#### Supplementary Tables

##### Supplementary Table S1. Crystallographic Data of CHCA_Rhodamine B

| | **CHCA_Rhodamine B** |
|---|---|
| Crystal data | |
| Chemical formula | $C_{28}H_{30}N_2O_3 \cdot C_{10}H_7NO_3$ |
| $M_r$ | 631.70 |
| Crystal system, space group | Triclinic, *P*-1 |
| Temperature (K) | 110 |
| *a*, *b*, *c* (Å) | 11.5312 (3), 12.0186 (3), 12.6873 (4) |
| α, β, γ (°) | 109.995 (2), 91.245 (2), 93.597 (2) |
| *V* (Å$^3$) | 1647.33 (8) |
| *Z* | 2 |
| Radiation type | Cu *K*α |
| μ (mm$^{-1}$) | 0.70 |
| Crystal size (mm) | 0.16 × 0.07 × 0.03 |
| Data collection | |
| Diffractometer | XtaLAB Synergy R, HyPix |
| Absorption correction | Analytical<br>*CrysAlis PRO* 1.171.42.80a (Rigaku Oxford Diffraction, 2023) Analytical numeric absorption correction using a multifaceted crystal model based on expressions derived by R.C. Clark & J.S. Reid. (Clark, R. C. & Reid, J. S. (1995). Acta Cryst. A51, 887-897) Empirical absorption correction using spherical harmonics, implemented in SCALE3 ABSPACK scaling algorithm. |
| $T_{min}$, $T_{max}$ | 0.934, 0.987 |
| No. of measured, independent and observed [$I > 2\sigma(I)$] reflections | 32312, 6460, 5364 |
| $R_{int}$ | 0.029 |

| | |
|---|---|
| $(\sin \theta/\lambda)_{max}$ (Å$^{-1}$) | 0.617 |
| Refinement | |
| $R[F^2 > 2\sigma(F^2)]$, $wR(F^2)$, $S$ | 0.047, 0.144, 1.05 |
| No. of reflections | 6460 |
| No. of parameters | 454 |
| No. of restraints | 33 |
| H-atom treatment | H atoms treated by a mixture of independent and constrained refinement |
| $\Delta r_{max}$, $\Delta r_{min}$ (e Å$^{-3}$) | 0.36, -0.25 |

**Computer programs:** *CrysAlis PRO* 1.171.42.49 (Rigaku OD, 2022), *SHELXT2018*/2 (Sheldrick, 2018), *SHELXL2019*/3 (Sheldrick, 2018), *SHELXTL* v6.10 (Sheldrick, 2008)[3].

**Supplementary Table S2. Crystallographic Data of nH_Alexa Fluor 488**

| | **nH_Alexa Fluor 488** |
|---|---|
| Crystal data | |
| Chemical formula | $C_{21}H_{11}N_2O_{11}S_2 \cdot 3(C_{11}H_9N_2) \cdot 7(H_2O)$ |
| $M_r$ | 1165.15 |
| Crystal system, space group | Triclinic, *P*-1 |
| Temperature (K) | 110 |
| *a*, *b*, *c* (Å) | 13.39220 (17), 14.7543 (2), 15.45243 (17) |
| α, β, γ (°) | 87.6615 (10), 71.7236 (11), 66.0207 (13) |
| *V* (Å$^3$) | 2635.79 (6) |
| *Z* | 2 |
| Radiation type | Cu *K*α |
| μ (mm$^{-1}$) | 1.65 |
| Crystal size (mm) | 0.34 × 0.11 × 0.06 |
| Data collection | |
| Diffractometer | XtaLAB Synergy R, HyPix |
| Absorption correction | Analytical<br>*CrysAlis PRO* 1.171.42.95a (Rigaku Oxford Diffraction, |

| | |
|---|---|
| | 2023) Analytical numeric absorption correction using a multifaceted crystal model based on expressions derived by R.C. Clark & J.S. Reid. (Clark, R. C. & Reid, J. S. (1995). Acta Cryst. A51, 887-897) Empirical absorption correction using spherical harmonics, implemented in SCALE3 ABSPACK scaling algorithm. |
| $T_{min}$, $T_{max}$ | 0.721, 0.921 |
| No. of measured, independent and observed [$I > 2\sigma(I)$] reflections | 83236, 10325, 9995 |
| $R_{int}$ | 0.022 |
| $(\sin \theta/\lambda)_{max}$ ($Å^{-1}$) | 0.617 |
| Refinement | |
| $R[F^2 > 2\sigma(F^2)]$, $wR(F^2)$, $S$ | 0.041, 0.110, 1.04 |
| No. of reflections | 10325 |
| No. of parameters | 834 |
| No. of restraints | 136 |
| H-atom treatment | H atoms treated by a mixture of independent and constrained refinement |
| **$\Delta\rho_{max}$, $\Delta\rho_{min}$ (e $Å^{-3}$)** | **0.85, -0.53** |

**Computer programs:** *CrysAlis PRO* 1.171.42.49 (Rigaku OD, 2022), *SHELXT2018*/2 (Sheldrick, 2018), *SHELXL2019*/3 (Sheldrick, 2018), *SHELXTL* v6.10 (Sheldrick, 2008)[3].

**Supplementary Results**

**Supplementary Single-Crystal X-Ray Diffraction Data for Co-Crystals from CHCA and Rhodamine B**

A mixture of 1:1.25 CHCA and rhodamine B dissolved in 50% toluene was prepared in a single pot. Subsequently, this mixture was allowed to evaporate at room temperature. After one week, several red crystals were obtained in block form. All reflection intensities were measured at 110.00(10) K using a Rigaku XtaLAB Synergy R (Rigaku) equipped with a rotating-anode X-ray source and HyPix-6000HE detector with Cu *K*α radiation (λ = 1.54178 Å) using CrysAlisPro (Version CrysAlisPro 1.171.42.49, Rigaku OD, 2022). The temperature of the data collection was controlled using a Cryostream 1000 system (Oxford Cryosystems, Oxford, United Kingdom). CrysAlisPro was also used to refine the cell dimensions and for data reduction. The structure was solved with the SHELXT-2018/2 software (Sheldrick, 2018) and was refined on $F^2$ with SHELXL-2019/3 (Sheldrick, 2018). Analytical numeric absorption correction using a multifaceted crystal was applied using CrysAlisPro. The H atoms were placed at calculated positions (unless otherwise specified) using the instructions AFIX 23, AFIX 43 or AFIX 137 with isotropic displacement parameters having values 1.2 or 1.5 $U_{eq}$ of the attached C atoms. The H atom attached to O3/O4 and O6 were found from difference Fourier maps. The coordinates for H6O (attached to O6) were refined pseudo-freely using the DFIX instruction to keep the O-H distance within an acceptable range. The structure is partly disordered. The moiety C14-C15 is disordered over two orientations, and the occupancy factor of the major component of the disorder refines to 0.743(6).

Response to Alert Level A (from checkCIF): PLAT355 ALERT 3 A Long O-H (X0.82, N0.98A) O4-H4O. 1.16 Å. The H atom H4O lies at a position somewhat intermediate between O3 and O4 (this is observed from the contoured difference Fourier map drawn in the plane defined by O3, O4 and H4O) as there is a strong hydrogen bond interaction found between the carboxylic acid – carboxylate pair (O3···O4: 2.4610(16) Å). The coordinates for H4O were refined freely in the final refinement. Most likely, CHCA and Rhodamine B can act both as carboxylic acid and carboxylate without any preference in the crystal. Hence, the unusually long O-H distance is an experimental artifact resulting from a space average over all possible positions of the O-H in the crystal.

Specified hydrogen bonds (with esds except fixed and riding H)

| D-H | H···A | D···A | <(DHA) | |
|---|---|---|---|---|
| 1.15(3) | 1.31(3) | 2.4610(16) | 174(2) | O4-H4O...O3 |
| 0.95(2) | 1.72(2) | 2.6318(18) | 160(2) | O6-H6O...O2_$3 |

**Supplementary Single-Crystal X-Ray Diffraction Data for Co-Crystals from nH and Alexa Fluor 488**

A mixture of 1:1 nH and Alexa Fluor 488 dissolved in 50% dichloromethane was prepared in a single-pot. Subsequently, this mixture was allowed to evaporate at room temperature. After three weeks, several amber-orange crystals were obtained in block form. All reflection intensities were measured at 110.00(10) K using a Rigaku XtaLAB Synergy R (equipped with a rotating-anode X-ray source and HyPix-6000HE detector) with Cu *K*α radiation (λ = 1.54178 Å) under the program CrysAlisPro (Version CrysAlisPro 1.171.42.49, Rigaku OD, 2022). The same program was used to refine the cell dimensions and for data reduction. The structure was solved with the program SHELXT-2018/2 (Sheldrick, 2018) and was refined on $F^2$ with SHELXL-2019/3 (Sheldrick, 2018). Analytical numeric absorption correction using a multifaceted crystal model was applied using CrysAlisPro. The temperature of the data collection was controlled using the system Cryostream 1000 from Oxford Cryosystems. The H atoms were placed at calculated positions (unless otherwise specified) using the instructions AFIX 43 with isotropic displacement parameters having values 1.2 $U_{eq}$ of the attached C atoms. The H atoms attached to N1, N2, N1X, N2X (X = A-C), and O*n*W (*n* = 1-7, lattice water molecules) were found from difference Fourier maps and their coordinates were refined pseudofreely using the DFIX instruction in order to keep the N−H, O−H and H···H distances within some acceptable ranges.

The structure is mostly ordered. The S2/O5−O7 moiety is disordered over two orientations, and the occupancy factor of the major component of the disorder refines to 0.817(12).

**Supplementary References**

1. Piacentino, M. L., Fasse, A. J., Camacho-Avila, A., Grabylnikov, I. & Bronner, M. E. SMPD3 expression is spatially regulated in the developing embryo by SOXE factors. *Dev. Biol.* **506**, 31–41 (2024).
2. Macrae, C. F. et al. Mercury CSD 2.0: New features for the visualization and investigation of crystal structures. *J. Appl. Crystallogr.* **41**, 466–470 (2008).
3. Sheldrick, G. M. Crystal structure refinement with SHELXL. *Acta Crystallogr. C Struct. Chem.* **71**, 3–8 (2015).
